# Co-Occurrence of Enzyme Domains Guides the Discovery of an Oxazolone Synthetase

**DOI:** 10.1101/2020.06.11.147165

**Authors:** Tristan de Rond, Julia E. Asay, Bradley S. Moore

**Author notes:** Author contributions: TdR and BSM designed research; TdR and JEA performed research; TdR analyzed data; TdR, JEA and BSM wrote the paper. The authors declare no competing interest.

## Abstract

Multidomain enzymes are cellular machines that orchestrate two or more catalytic activities to carry out metabolic transformations with increased control and speed. Our understanding of these enzymes’ capabilities drives progress in fundamental metabolic research, biocatalysis, and human health. Here, we report the development of a new genome mining approach for the targeted discovery of novel biochemical transformations through the analysis of co-occurring enzyme domains (CO-ED) in a single protein. CO-ED was designed to identify unannotated multifunctional enzymes for functional characterization and discovery based on the premise that linked enzyme domains have evolved to function collaboratively. Guided by CO-ED, we targeted an unannotated predicted ThiF-nitroreductase di-domain enzyme found in more than 50 proteobacteria. Through heterologous expression and biochemical reconstitution, we discovered a series of new natural products containing the rare oxazolone (azlactone) heterocycle and characterized the di-domain enzyme as the first reported oxazolone synthetase in biology. This enzyme has the potential to become a valuable biocatalyst for the production of versatile oxazolone synthetic intermediates. This proof-of-principle experiment validates CO-ED-guided genome mining as a new method with potential broad utility for both the discovery of novel enzymatic transformations and the functional gene annotation of multidomain enzymes.

**TOC graphic:** 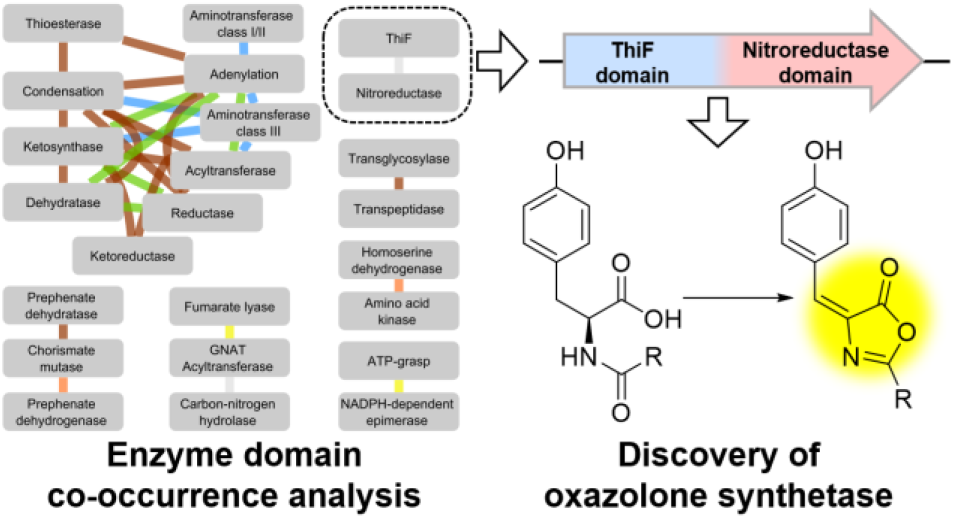

## Introduction

Our knowledge of nature’s diversity of enzymatic transformations is crucial to advancing research in a multitude of disciplines. For instance, our ability to predict metabolic capacity from genome sequences enables new insights in human health and ecology^1,2^, while the development of bioprocesses to produce chemicals in an environmentally benign fashion relies on the availability of a well-stocked biocatalytic toolbox^3^. Billions of years of evolution has resulted in immense natural genetic diversity, which we are rapidly starting to uncover using modern sequencing technologies. However, the functional assignment of the enzymes encoded by this sequence diversity is lagging^4–6^. Many enzymes catalyzing chemical transformations previously not known to occur in nature are still being discovered^7^, and there is no end in sight of unannotated and mis-annotated enzymes in genomic databases^8^. The search for novel enzymes in this genomic wilderness can fulfill the dual purpose of functional gene annotation and biocatalyst discovery.

The computational searches for biosynthetic enzymes that underlie “genome mining” campaigns are most commonly based on gene homology to known core biosynthetic genes of well-established classes of medicinal natural products such as polyketides, non-ribosomal peptides and terpenoids^9^. This approach, while likely to identify gene clusters that produce bio-active products, is prone to the re-discovery of known enzymology. Therefore, we set out to develop a genome mining strategy that could guide enzyme discovery in a complimentary manner.

One notable feature amongst specialized metabolism is an overabundance of enzymes harboring multiple catalytic domains (Figure S1). Multidomain enzymes are thought to arise through gene fusion of two or more single-domain enzymes, affording catalytic advantages such as coupled temporal and spatial regulation, a fixed active site stoichiometry, and channeling of reactive intermediates^10,11^. Over time, a multidomain enzyme may even evolve to orchestrate the reactivity of its constituent active sites to such an extent that it acquires a new function^12^. Classic examples of multidomain enzymes are the fatty acid synthase, polyketide synthase (PKS), and nonribosomal peptide synthetase (NRPS) assembly-line enzymes^13^, some of which consist of dozens of domains. More recently, multidomain enzymes have been shown to catalyze epimerization^14,15^ and *N-*nitrosation^16,17^ reactions.

Here, we present a genome mining paradigm where we leverage the evolved co-occurrence of enzymatic domains (CO-ED) in proteins to inform enzyme discovery. Through CO-ED analysis, we discovered a series of new oxazolone-containing natural products, as well as the first reported oxazolone synthetase, a novel bifunctional cyclodehydratase–oxidoreductase.

## Results

### CO-ED–guided identification of a di-domain enzyme candidate

The CO-ED workflow takes a query set of proteins and generates a network representation of co-occurring enzymatic domains (Figure 1). A node is drawn for each Pfam^18^ domain found in the query proteins and matching a curated set of enzymatic domains, and an edge is drawn between two nodes for each two-domain combination of domains found together in at least one query protein. In parallel, a set of characterized enzymes, derived from the BRENDA^19^, MIBiG^20^, and Uniprot^21^ databases, is subjected to the same analysis. If an edge in the network represents a domain pair found in a previously characterized multidomain enzyme, the edge is color coded according to the originating database. Remaining uncolored edges, therefore, represent enzyme domain combinations that are likely uncharacterized.

**Figure 1 (one-column).**
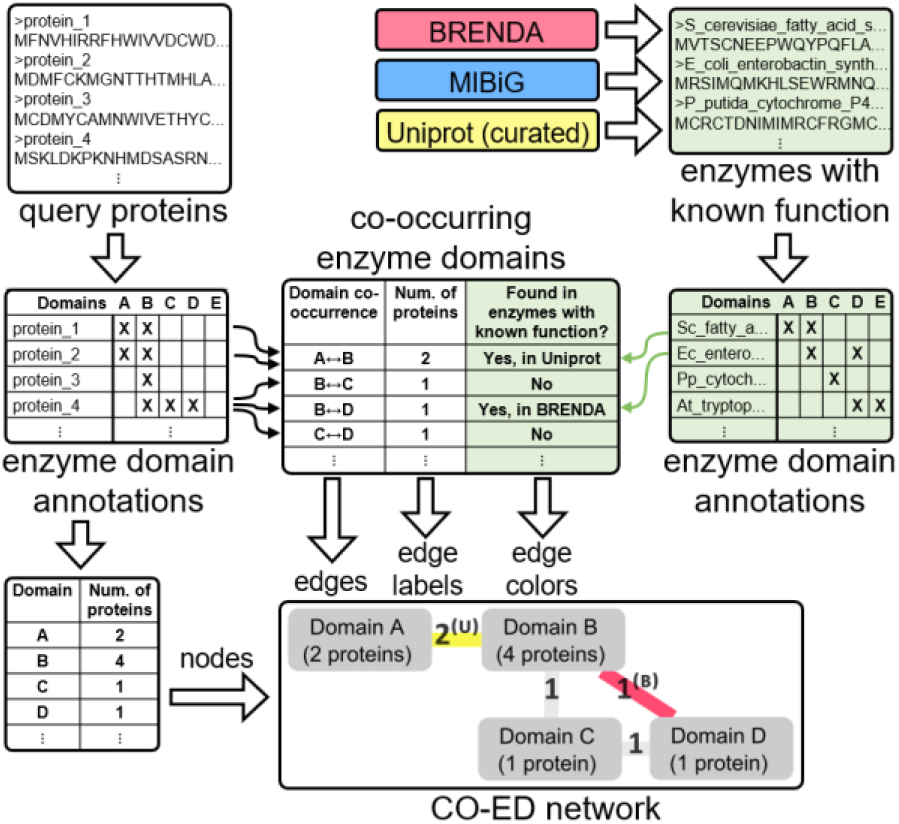
Outline of the CO-ED workflow. A CO-ED network consists of nodes representing a curated set of enzymatic domains present in a query set of proteins, and edges connecting two domains if they co-occur in the same protein. Edges are colored based on whether any known enzymes contain the pair of domains the edge represents. Known enzymes were compiled from BRENDA^19^ (www.brenda-enzymes.org), MIBiG^20^ (mibig.secondarymetabolites.org) and a curated set entries from Uniprot^21^ (www.uniprot.org), the edge color indicating the originating database(s) for the annotation. For details, please refer to the methods in the SI or the publicly-available code.

To validate the CO-ED workflow, we first applied it to the genome of the model organism *Escherichia coli* K12. The *E. coli* network (Figs. 2a,b, S2, S3, Dataset S1) revealed 19 co-occurring pairs of domains, including some well-studied multifunctional enzymes including penicillin-binding proteins^22^, the aldehyde dehydrogenase–alcohol dehydrogenase AdhE^23^, the UDP-L-Ara4FN biosynthesis enzyme ArnA^24^, the amino acid biosynthesis enzymes TrpCF^25^, TrpGD^26^ and HisIE^27^, and the enterobactin NRPS EntF^28^. Every edge is colored, suggesting that the function of every multidomain enzyme in *E. coli* detected by CO-ED has likely already been established.

**Figure 2 (two-column).**
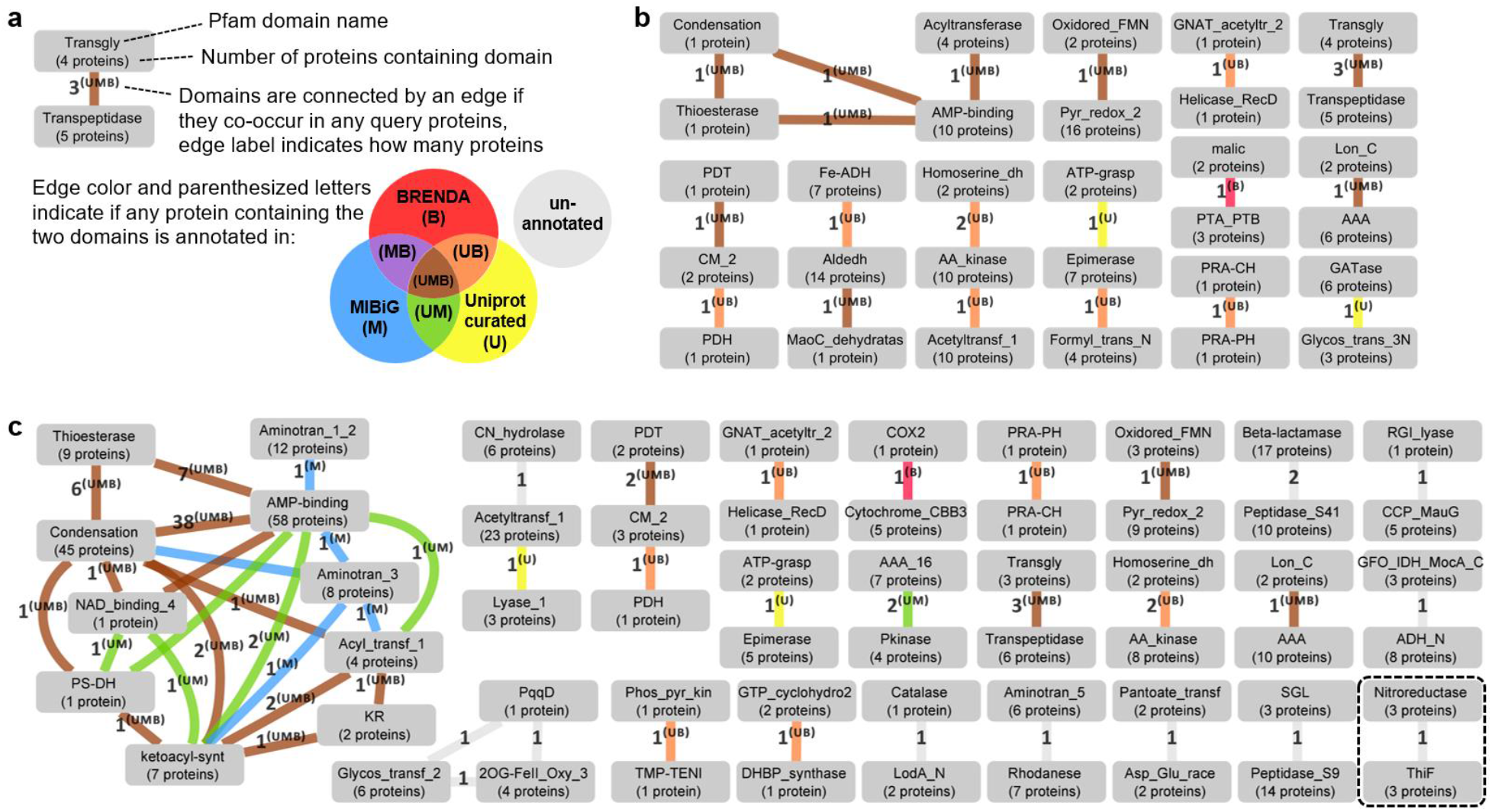
CO-ED networks can help guide enzyme discovery. **a:** Legend for the CO-ED networks shown in **b** and **c**. **b:** CO-ED network for *E. coli* K12 (NCBI ASM584v2^77^). All edges are colored, suggesting *E. coli* is of low priority for CO-ED–guided genome mining. **c:** CO-ED network for *Pseudoalteromonas rubra* DSM 6842 (NCBI “Prub_2.0”^78^). Some overlapping edge labels are omitted for clarity. Many edges are unannotated (gray), suggesting there is much potential for exploring multidomain enzymes for novel biosynthetic capacity in this organism. The ThiF-nitroreductase domain pair investigated further in this study is highlighted Networks were rendered in Cytoscape^79^. Only the connected components are shown.

We next applied CO-ED analysis to the genome of the biosynthetically talented^29,30^, non-model marine gammaproteobacterium *Pseudoalteromonas rubra* DSM 6842, chosen because of our experience heterologously expressing enzymes from this and closely-related species^31–33^. The *P. rubra* CO-ED network (Figs. 2c, S4, Dataset S1) revealed numerous NRPSs, as can be inferred from the edge indicating that the “AMP-binding” (also known as adenylation) and “Condensation” Pfam domains co-occur in thirty-eight proteins, including two hybrid PKS/NRPS enzymes, as indicated by the two proteins containing both “ketoacyl-synt” and “Condensation” Pfam domains. The network also shows a putative bifunctional RibBA, known to be involved in riboflavin biosynthesis in other species^34^, as well as some of the same bifunctional primary metabolic enzymes observed in *E. coli*. However, unlike the *E. coli* network, twelve domain pairs are unannotated, suggesting that P. rubra may harbor multidomain enzymes that catalyze novel enzymatic transformations.

A *P. rubra* protein harboring a “ThiF”-“nitroreductase” domain pair (highlighted in bottom right of Fig. 2c) caught our attention, because each of these domains is known to catalyze a diverse set of chemical transformations (Fig. 3a), yet their combination is unprecedented. Enzymes of the ThiF protein family catalyze various vital reactions that proceed through carboxylate adenylation, including the activation of ubiquitin and ubiquitin-like proteins by E1 enzymes^35^, the cyclization of *N*^6^-threonylcarbamoyladenosine in tRNA maturation^36^, and the post-translational modification of ribosomally-synthesized peptide antibiotics^37,38^. ThiF family enzymes also play a role in the sulfur incorporation machinery for thiamin, the molybdenum cofactor and some natural products^39,40^. To date, all characterized substrates of ThiF family enzymes have been polypeptides or tRNA. The nitroreductase family consists of flavoenzymes that catalyze a remarkable variety of redox reactions^41^, such as the reductive deiodination of thyroid hormones^42^, the dehydrogenation of oxazolines and thiazolines^43,44^, the fragmentation of flavin mononucleotide to form the vitamin B12 lower ligand 5,6-dimethylbenzimidazole^45^, enol methylene reduction in cofactor F420 biosynthesis^46–48^, diketopiperazine desaturation in albonoursin biosynthesis^49^, and the reductive detoxification of various nitro functional groups^50^.

**Figure 3 (two-column).**
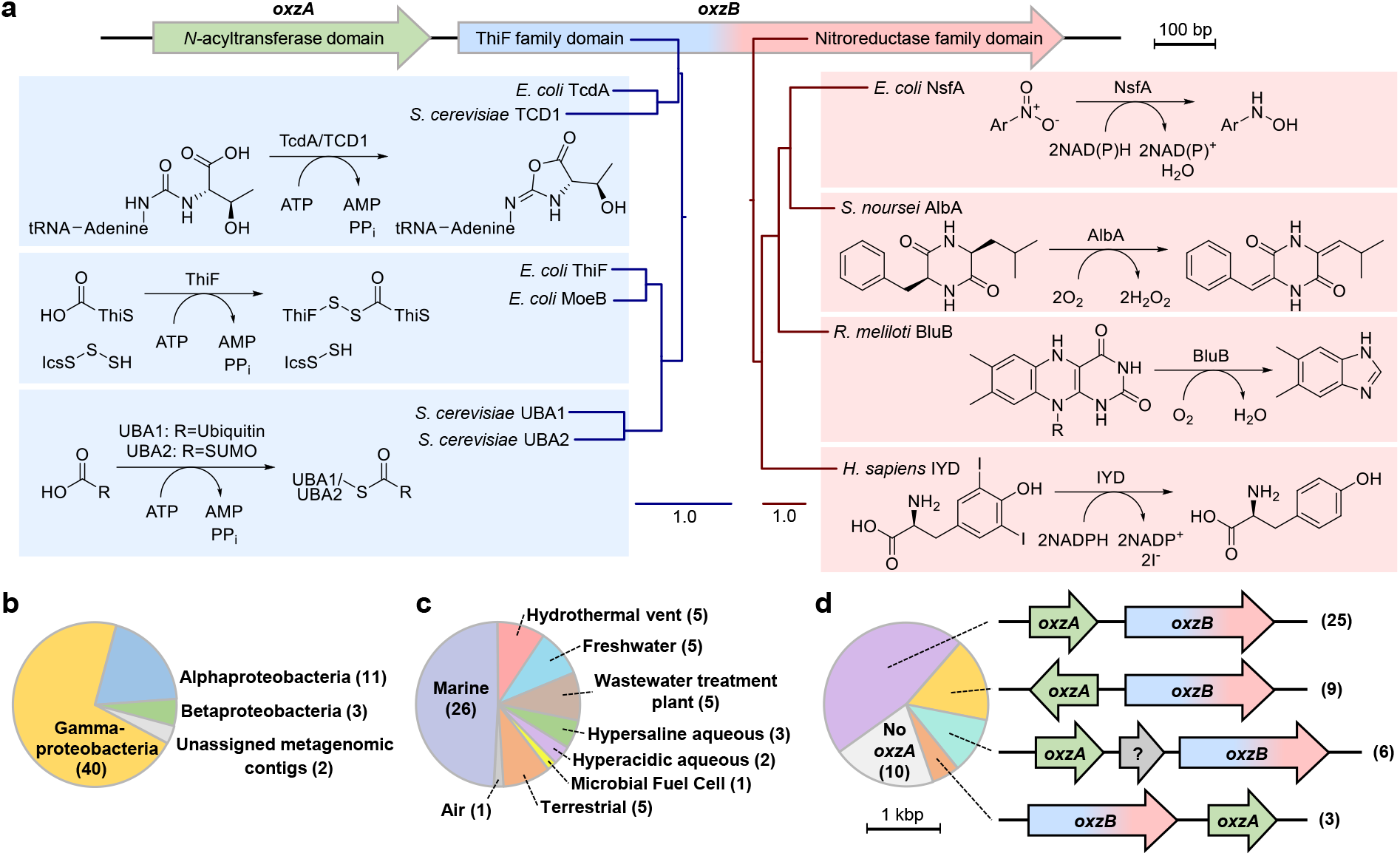
Survey of transformations known to be catalyzed by the individual domains of a di-domain enzyme identified by CO-ED, and properties of organisms harboring homologs of this enzyme. **a:** OxzB exhibits homology to both the ThiF and nitroreductase enzyme families. Shown is a selection of characterized members of these enzyme families along with the transformations they catalyze, along with midpoint-rooted gene trees showing the relationships between each other and OxzB. ThiF family enzymes are known to catalyze ATP-dependent carboxylate activating reactions, while nitroreductase family enzymes catalyze a variety of redox reactions. Inferred phylogenies were generated from protein sequences using neighbor joining on the MAFFT web server^80^ and midpoint-rooted. Scale bars designate 1 substitution per site. **b,c:** Phylogenetic distribution (**b**) and habitat (**c**) of the host organisms harboring genes encoding OxzB proteins represented in the Uniprot database. Organisms with unknown habitats are not included. **d:** Genomic context of *oxzB* genes as determined using the Enzyme Function Initiative Genome Neighborhood Tool^60^. Most *oxzB* homologs are accompanied by *oxzA*, which codes for an *N*-acyltransferase. The arrow labeled “?” represents genes unrelated to *oxzA* or *oxzB*. Organisms with unknown *oxzB* genomic context (e.g., at the edge of a contig) are not included.

Querying the Uniprot database with the di-domain ThiF-nitroreductase enzyme from *P. rubra,* which we named *oxzB*, revealed 56 proteins originating from alpha-, beta-, and gammaproteobacteria (Figs. 3b, S5, Table S1) isolated from a variety of (predominantly aquatic) environments (Fig. 3c). In those cases where the genomic context of the *oxzB* homolog was known, it is typically accompanied by an *N*-acyltransferase homolog (*oxzA*). In *P. rubra* and many other species, *oxzA* is immediately upstream of *oxzB* on the same strand, forming an apparent two-gene *oxzAB* operon (Fig. 3d).

### Heterologous expression of *oxzAB* leads to production of novel oxazolone metabolites

To investigate the function of the *oxzAB* genes, we PCR amplified *oxzAB* from *P. rubra*, *Rheinheimera pacifica*, *Colwellia chukchiensis* (all gammaproteobacteria), *Skermanella aerolata* (alphaproteobacterium), and *Undibacterium pigrum* (betaproteobacterium), and heterologously expressed the gene pairs in *E. coli*. Colonies on solid media and cell pellets after growth in liquid media both turned visibly yellow (Fig. S6). The color would briefly intensify when incubated above pH 9, suggesting the possibility of a phenolic, base-labile product (Figs. S6, S7). HPLC-UV-MS analysis of extracts of the pellets revealed several chromatographic peaks with absorbance in the 300-400 nm range which were not produced by *E. coli* natively (Fig. 4a). The product profiles of heterologously-expressed *oxzAB* genes originating from different species shared many constituents, and the inclusion of genes found up- or downstream from *oxzAB* did not substantially change the product profiles (Fig. S8).

**Figure 4 (two-column).**
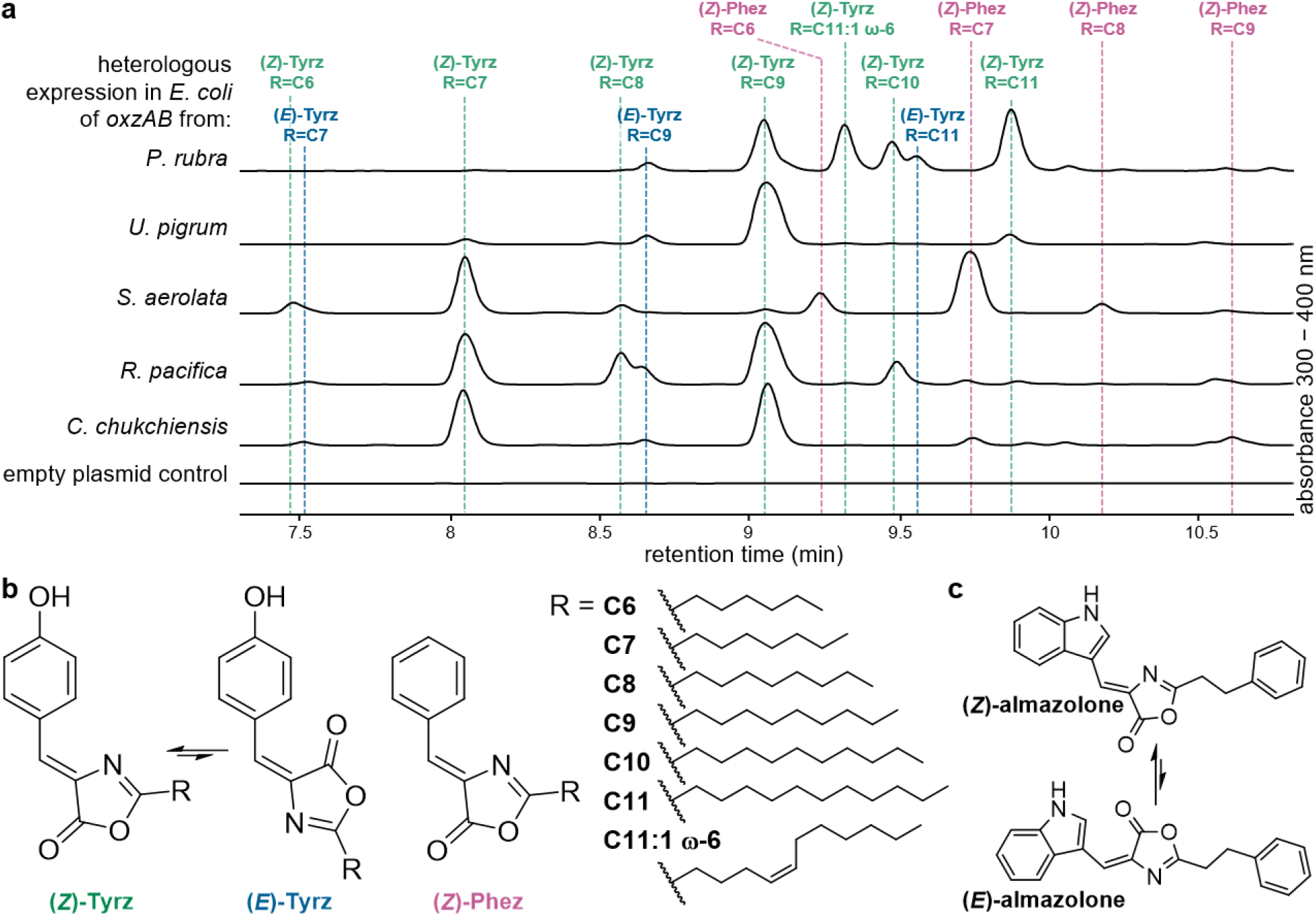
Heterologous expression of *oxzAB* results in the production of a series of oxazolones. **a:** Reversed phase HPLC profiles of extracts of *E. coli* expressing *oxzAB* from various proteobacteria. An absorbance range of 300-400 nm was chosen to detect all oxazolones reported in this manuscript (under our chromatography conditions (H_2_O/acetonitrile with 0.1% formic acid), the λ_*max*_ of the (*E*)-tyrazolones is 360 nm, that of the (*Z*)-tyrazolones is 358 nm, and that of the (*Z*)-phenazolones is 330 nm). **b:** Structures of the tyrazolones and phenazolones, which are found as a series with varying alkyl tails. (*Z*)-Tyrz, R = C9, C10, C11 and C11:1 ω-6 and (*Z*)-Phez, R = C7 were characterized by NMR, while all others were inferred by exact mass and MS/MS fragmentation patterns. The tyrazolones exist as (*E*)/(*Z*) isomers in equilibrium. **c:** Almazolone, an oxazolone natural product previous isolated from an alga, also occurs as an (*E*)/(*Z*) equilibrium.

Through a combination of mass spectrometry and spectroscopic techniques, with notable insight provided by 1,1-ADEQUATE NMR, ^1^H-^15^N HMBC NMR, and the identity of degradation products formed in basic methanol^51,52^, we were able to assign the products of *oxzAB* as a series of oxazolones (Figs. 4b, S9, S10). We named these seemingly tyrosine- and phenylalanine-derived molecules tyrazolones and phenazolones respectively. The major products were heptyl-nonyl-, undecyl-, and ω-6-undecenyltyrazolone (Tyrz, R = C7, C9, C11 and C11:1 ω-6), which were found to occur in an equilibrium of (*E*) and (*Z*) isomers with the latter being predominant, and (*Z*)-heptylphenazolone (Phez, R = 7). None of these products have previously been reported before in the literature, however an oxazolone that appears to be derived from tryptophan, almazolone (Fig. 4c), was isolated from a red alga^53^ and its structure confirmed through synthesis. Almazolone similarly exists as an (*E*)/(*Z*) equilibrium, forms degradation products analogous to those formed by the tyrazolones and phenazolones, and has similar spectroscopic properties, lending credence to our structural assignments.

### *In vitro* reconstitution of oxazolone biosynthesis by OxzAB

Considering the enzymatic domains contained within *oxzAB*, we hypothesized oxazolone biosynthesis to proceed as follows: OxzA forms an *N*-acyl amino acid using an acyl-CoA derived from the natural fatty acyl-CoA beta oxidation pool. In order to form the oxazolone, OxzB then catalyzes cyclization and oxidation of the *N*-acyl amino acid, in either order (Fig. S11). The ThiF family enzyme TcdA is known to catalyze cyclodehydration of an amino acid moiety, and the nitroreductase family enzyme AlbA has been shown to oxidize amino acid-derived substrates (Fig. 2a). It is thus conceivable that OxzB, which harbors both these domains, could act to form the oxazolone heterocycle.

To test our biosynthetic hypothesis, we heterologously expressed and purified *P. rubra* OxzA and OxzB in *E. coli* as N-terminal His6 and MBP fusion proteins, respectively, and assayed their activity by HPLC (Fig. 5). When incubated with L-tyrosine and decanoyl-CoA, OxzA catalyzed the formation of *N*-decanoyl-L-tyrosine. OxzB was able to catalyze the formation of nonyltyrazolone both from synthetic *N*-decanoyltyrosine as well as when combined with OxzA, tyrosine and decanoyl-CoA. OxzB activity was only observed in the presence of ATP, as is typical for ThiF family enzymes. (*Z*)-nonyltyrazolone, the favored isomer of the (*E*)/(*Z*) equilibrium, was the major product upon extended incubation of the enzymatic reactions, however, shorter incubation times revealed a proportionally greater amount of (*E*)-tyrazolone (Fig. S12), suggesting that the immediate product of OxzB is (*E*)-nonyltyrazolone.

**Figure 5 (one-column).**
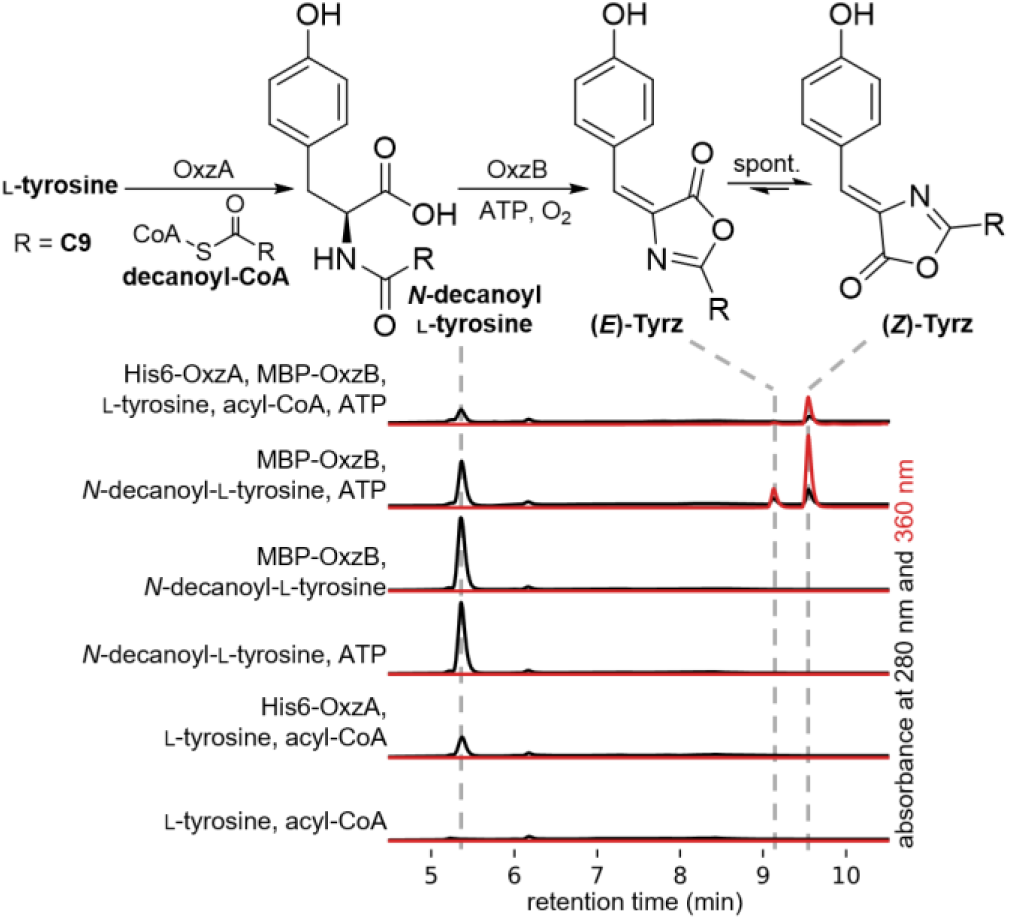
*In vitro* characterization of oxazolone biosynthesis by *P. rubra* OxzA and OxzB. Black HPLC chromatograms show absorbance at 280 nm (*N*-decanoyl-L-tyrosine); red chromatograms show absorbance at 360 nm (tyrazolones) and are scaled down 4.5-fold in relation to the 280 nm signal.

### Analysis of oxazolone production in the native hosts

To determine if the oxazolones are true natural products or merely artifacts of heterologous expression, we attempted to detect these molecules in the five bacteria whose *oxzAB* gene pairs we tested heterologously above. Under standard culturing conditions, oxazolones could be reliably detected only in *C. chukchiensis*. Encouraged by reports of chemical elicitation methods to activate silent gene clusters^54^, we evaluated a series of antibiotics to induce oxazolone production in *P. rubra* or *C. chulkchiensis* at sublethal concentrations (Figs. 6, S13). Gratifyingly, we could elicit oxazolone production in *P. rubra* with erythromycin or chloramphenicol, both known inhibitors of protein synthesis^55^, with oxazolone peak areas similar to those of uninduced *C. chukchiensis*. Oxazolone production in *C. chukchiensis* could be further induced by several antibiotics, only some of which inhibit protein synthesis. We were unable to detect tyrazolones or phenazolones in extracts of the other three species whose *oxzAB* gene pair we validated heterologously.

**Figure 6 (one-column).**
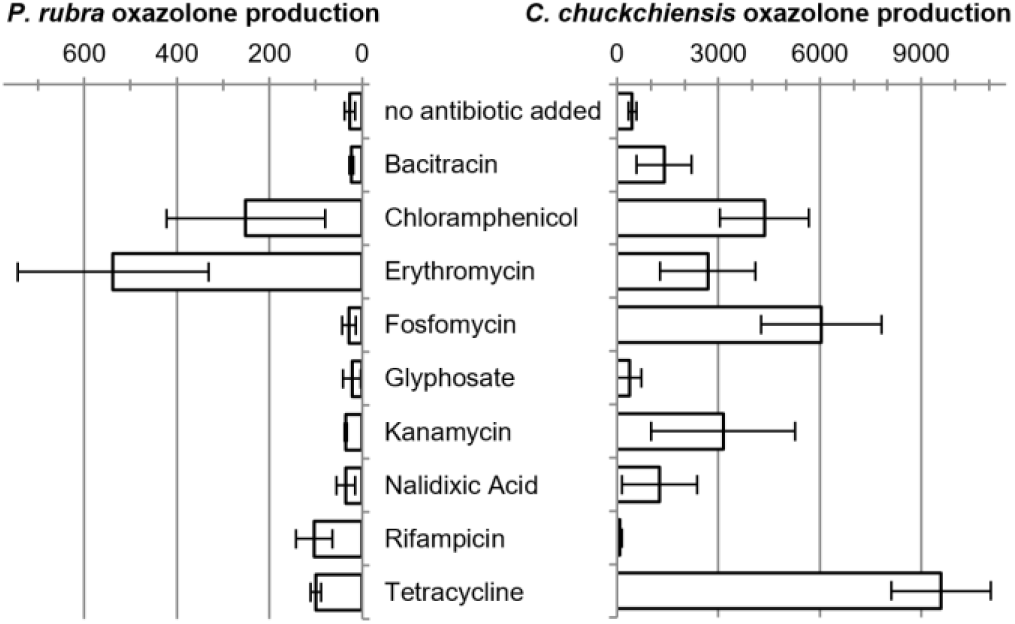
Induction of oxazolone production in *P. rubra* and *C. chukchiensis* by various antibiotics. Bacteria were grown as a lawn on Marine Agar 2216 with a drop of antibiotic. A consistent amount of biomass adjacent to the zone of inhibition was harvested, extracted and analyzed by HPLC. Bars indicate summed peak areas between wavelengths of 300 nm and 400 nm. Error bars represent standard deviation of three biological replicates.

## Discussion

### CO-ED as a method to prioritize multidomain enzyme for genome mining

Using a CO-ED–guided genome mining strategy, we discovered a series of novel oxazolone natural products as well as a unique oxazolone synthetase harboring ThiF and nitroreductase domains. This proof-of-principle experiment showcases the promise of CO-ED in selecting orphan multidomain enzymes for targeted functional discovery. Many computational workflows for the prioritization of unannotated genes for genome mining rely on identifying gene clusters based on sequence homology to known biosynthetic genes, as is the case for e.g., PRISM^56^ and AntiSMASH^57^. Downstream analysis tools, such as BiGSCAPE^58^, in turn often depend on AntiSMASH gene cluster annotations. This approach risks overlooking genes that do not either themselves have homology to known biosynthetic genes, or are clustered with such genes. Alternatively, there exist workflows that allow the user to explore enzyme families related to a query sequence or domain, such as EnzymeMiner^59^ and the tools hosted by the Enzyme Function Initiative^60^, but these might miss protein families that the user did not consider searching for. CO-ED is complementary to both the aforementioned approaches in that it considers unannotated enzyme domain co-occurrences in a relatively unbiased fashion.

The analysis of co-occurring protein domains has previously been applied in contexts other than genome mining, such as in evolutionary studies^61,62^, to help identify antifungal drug targets^63^, and in functional gene annotation (where it is sometimes called the “Rosetta Stone” method)^64,65^. Parallel to this, efforts have been made to connect protein domains to enzymatic reactions^60,66–69^. CO-ED represents, to our knowledge, the first time these concepts have been united to specifically guide the discovery of enzymes catalyzing novel transformations. We plan to develop a user-friendly web tool to facilitate CO-ED analysis on any set of query sequences in the near future.

While CO-ED analysis has revealed that all multidomain enzymes in *E. coli* likely have known functions, we wished to explore whether this is the case for other model bacteria, particularly those that are promising sources of novel biocatalysts. To this end, we applied CO-ED to the genomes of *Streptomyces coelicolor* A3(2), *Salinispora tropica* CNB-440, and *Pseudomonas fluorescens* Pf-5 – model organisms whose remarkable biosynthetic capacities are well-studied^70–72^. We found over a dozen unannotated domain pairs in each of these species (Figs. S14-S16). Investigating the functions of these multidomain enzymes may reveal yet more biochemical novelty even in these well-studied organisms.

CO-ED analysis of all proteins in the Uniprot^21^ database (Fig. S17 and Datatset S1) shows that while the most abundantly distributed co-occurring domain pairs have enzymes annotated in BRENDA, MIBiG, or Uniprot, there is a still a plethora of less-widespread multidomain enzymes whose functions are unassigned: Of the 252 enzymatic domain pairs found in at least 1000 proteins, 78.6% are annotated, whereas 82.8% of the 3381 domain pairs found in at least 10 proteins are unannotated. These annotation rates may be an under-estimate because of a lag in the appearance of new results in the databases, or an over-estimate if the same combination of enzymatic domains catalyzes different transformations in different proteins. Nevertheless, there are many more multidomain enzymes awaiting examination for new fundamental biochemical insight and as potential biocatalysts.

### A newly-discovered oxazolone synthetase

Oxazolones – also known as azlactones – are versatile synthetic intermediates^73,74^, but no biocatalytic methods for their production have been reported. Biocatalytic approaches have the potential to exhibit better functional group tolerance than synthetic methods, which could shorten synthetic routes that rely on oxazolones. Besides OxzB, the only other known oxazolone-forming enzyme is MbnBC, which is involved in the biosynthesis of the chalkophore methanobactin^75^. MbnBC catalyzes a 4-electron oxidative rearrangement – a fundamentally different reaction from OxzB – and acts on a polypeptide substrate, which may pose challenges to developing of this enzyme as a biocatalyst. Considering that *P. rubra* OxzB is a soluble enzyme that acts on small molecule substrates, it and its homologs have the potential to become valuable biocatalysts.

Characterization of the product profiles of five *oxzAB* gene pairs led to the discovery of several novel oxazolone natural products. Dozens more *oxzAB* gene pairs are present in sequence databases, ready to be explored for further biosynthetic novelty. Given the structural similarity of almazolone to the oxazolones described here, we expect a yet-undiscovered OxzB-like enzyme selective for a tryptophan derivative to be involved in almazolone biosynthesis. Some *oxzB* genes are clustered with predicted glycosyltransferases, PKSs and NRPSs, suggesting they may be part of biosynthetic gene clusters coding for more complex oxazolone-containing natural products.

The reaction catalyzed by OxzB can formally be understood to consist of a cyclization and an oxidation, presumably catalyzed by the ThiF-like and nitroreductase domains respectively. Ongoing mechanistic studies aim to determine the order in which these reactions occur, and whether any soluble intermediates are formed. OxzB is also the first characterized ThiF family enzyme that does not act on a macromolecular substrate but on a small molecule, potentially making it a useful model to gain insight into this important enzyme family^76^.

### The biological role of oxazolones

We have shown that *P. rubra* and *C. chukchiensis* can produce oxazolone natural products, but it is unclear what the biological role of these molecules is. The oxazolone-containing methanobactins are chalkophores, but this function is mediated by thiolates that are not present in the tyrazolones or phenazolones. The observation that production of these molecules can be induced by antibiotics suggests that they may act as signals in response to stress. Like many signaling molecules, the oxazolones are somewhat labile in water; at pH 8 (e.g., seawater) they hydrolyze with a half-life on the order of hours. We have thus far observed no obvious phenotypes when bacteria (including the five species whose *oxzAB* genes we studied heterologously) are exposed to these molecules, both in the presence and absence of antibiotics. As we continue to explore the distribution and repertoire of oxazolone biosynthetic genes, we hope to learn more about the biological functions of these newly discovered bacterial natural products.

## Associated Content

The Supporting Information contains Supplementary Figures, a Supplementary Table containing Uniprot accession numbers for the proteins investigated, Methods, and Spectral data.

Jupyter notebooks containing Python code for CO-ED analysis and to generate the statistics shown in Figure S1 are publicly available at https://github.com/tderond/CO-ED.

Pre-generated CO-ED networks are available as Dataset S1 and can be opened with Cytoscape

## Acknowledgements

We would like to thank our UCSD colleagues Jie Li, Vikram Shende, and Brendan Duggan for helpful discussions. This work was supported by National Institutes of Health awards F32GM129960 to TdR and R01GM085770 to BSM, as well as the American Society for Pharmacognosy Undergraduate Research Award and the UC San Diego “Eureka” Undergraduate Research Scholarship to JEA.

## References

(1) O’Brien, E. J.; Monk, J. M.; Palsson, B. O. Using Genome-scale Models to Predict Biological Capabilities. Cell 2015, 161, 971–987.

(2) Gu, C.; Kim, G. B.; Kim, W. J.; Kim, H. U.; Lee, S. Y. Current Status and Applications of Genome-scale Metabolic Models. Genome Biol 2019, 20, 121.

(3) Bornscheuer, U. T. The Fourth Wave of Biocatalysis Is Approaching. Philos Transact A Math Phys Eng Sci 2018, 376.

(4) Gerlt, J. A.; Allen, K. N.; Almo, S. C.; Armstrong, R. N.; Babbitt, P. C.; Cronan, J. E.; Dunaway-Mariano, D.; Imker, H. J.; Jacobson, M. P.; Minor, W.; et al. The Enzyme Function Initiative. Biochemistry 2011, 50, 9950–9962.

(5) Hanson, A. D.; Pribat, A.; Waller, J. C.; de Crécy-Lagard, V. Unknown Proteins and Orphan Enzymes: The Missing Half of the Engineering Parts List -;and How to Find It. Biochem J 2009, 425, 1–11.

(6) Schnoes, A. M.; Brown, S. D.; Dodevski, I.; Babbitt, P. C. Annotation Error in Public Databases: Misannotation of Molecular Function in Enzyme Superfamilies. PLoS Comput Biol 2009, 5, e1000605.

(7) Scott, T. A.; Piel, J. The Hidden Enzymology of Bacterial Natural Product Biosynthesis. Nat. Rev. Chem. 2019, 3, 404–425.

(8) Ellens, K. W.; Christian, N.; Singh, C.; Satagopam, V. P.; May, P.; Linster, C. L. Confronting the Catalytic Dark Matter Encoded by Sequenced Genomes. Nucleic Acids Res 2017, 45, 11495–11514.

(9) Medema, M. H.; Fischbach, M. A. Computational Approaches to Natural Product Discovery. Nat Chem Biol 2015, 11, 639–648.

(10) Michael, A. J. Evolution of Biosynthetic Diversity. Biochem J 2017, 474, 2277–2299.

(11) Hagel, J. M.; Facchini, P. J. Tying the Knot: Occurrence and Possible Significance of Gene Fusions in Plant Metabolism and Beyond. J Exp Bot 2017, 68, 4029–4043.

(12) Bashton, M.; Chothia, C. The Generation of New Protein Functions by the Combination of Domains. Structure 2007, 15, 85–99.

(13) Weissman, K. J. The Structural Biology of Biosynthetic Megaenzymes. Nat Chem Biol 2015, 11, 660–670.

(14) Farrow, S. C.; Hagel, J. M.; Beaudoin, G. A. W.; Burns, D. C.; Facchini, P. J. Stereochemical Inversion of (S)-reticuline by a Cytochrome P450 Fusion in Opium Poppy. Nat Chem Biol 2015, 11, 728–732.

(15) Winzer, T.; Kern, M.; King, A. J.; Larson, T. R.; Teodor, R. I.; Donninger, S. L.; Li, Y.; Dowle, A. A.; Cartwright, J.; Bates, R.; et al. Plant Science. Morphinan Biosynthesis in Opium Poppy Requires a P450-oxidoreductase Fusion Protein. Science 2015, 349, 309–312.

(16) Ng, T. L.; Rohac, R.; Mitchell, A. J.; Boal, A. K.; Balskus, E. P. An N-nitrosating Metalloenzyme Constructs the Pharmacophore of Streptozotocin. Nature 2019, 566, 94–99.

(17) He, H.; Henderson, A. C.; Du, Y. -L.; Ryan, K. S. A Two-enzyme Pathway Links L-arginine to Nitric Oxide in N-nitroso Biosynthesis. J Am Chem Soc 2019, 141, 4026–4033.

(18) El-Gebali, S.; Mistry, J.; Bateman, A.; Eddy, S. R.; Luciani, A.; Potter, S. C.; Qureshi, M.; Richardson, L. J.; Salazar, G. A.; Smart, A.; et al. The Pfam Protein Families Database in 2019. Nucleic Acids Res 2019, 47, D427–D432.

(19) Jeske, L.; Placzek, S.; Schomburg, I.; Chang, A.; Schomburg, D. BRENDA in 2019: a European ELIXIR Core Data Resource. Nucleic Acids Res 2019, 47, D542–D549.

(20) Kautsar, S. A.; Blin, K.; Shaw, S.; Navarro-Muñoz, J. C.; Terlouw, B. R.; van der Hooft, J. J. J.; van Santen, J. A.; Tracanna, V.; Suarez Duran, H. G.; Pascal Andreu, V.; et al. MIBiG 2.0: a Repository for Biosynthetic Gene Clusters of Known Function. Nucleic Acids Res 2020, 48, D454–D458.

(21) The UniProt Consortium. UniProt: The Universal Protein Knowledgebase. Nucleic Acids Res 2018, 46, 2699.

(22) Ishino, F.; Mitsui, K.; Tamaki, S.; Matsuhashi, M. Dual Enzyme Activities of Cell Wall Peptidoglycan Synthesis, Peptidoglycan Transglycosylase and Penicillin-sensitive Transpeptidase, in Purified Preparations of Escherichia Coli Penicillin-binding Protein 1A. Biochem Biophys Res Commun 1980, 97, 287–293.

(23) Goodlove, P. E.; Cunningham, P. R.; Parker, J.; Clark, D. P. Cloning and Sequence Analysis of the Fermentative Alcohol-dehydrogenase-encoding Gene of Escherichia Coli. Gene 1989, 85, 209–214.

(24) Williams, G. J.; Breazeale, S. D.; Raetz, C. R. H.; Naismith, J. H. Structure and Function of Both Domains of ArnA, a Dual Function Decarboxylase and a Formyltransferase, Involved in 4-amino-4-deoxy-L-arabinose Biosynthesis. J Biol Chem 2005, 280, 23000–23008.

(25) Kaufmann, M.; Schwarz, T.; Jaenicke, R.; Schnackerz, K. D.; Meyer, H. E.; Bartholmes, P. Limited Proteolysis of the Beta 2-dimer of Tryptophan Synthase Yields an Enzymatically Active Derivative That Binds Alpha-subunits. Biochemistry 1991, 30, 4173–4179.

(26) Li, S. L.; Hanlon, J.; Yanofsky, C. Separation of Anthranilate Synthetase Components I and II of Escherichia Coli, Salmonella Typhimurium, and Serratia Marcescens and Determination of Their Amino-terminal Sequences by Automatic Edman Degradation. Biochemistry 1974, 13, 1736–1744.

(27) Carlomagno, M. S.; Chiariotti, L.; Alifano, P.; Nappo, A. G.; Bruni, C. B. Structure and Function of the Salmonella Typhimurium and Escherichia Coli K-12 Histidine Operons. J Mol Biol 1988, 203, 585–606.

(28) Rusnak, F.; Sakaitani, M.; Drueckhammer, D.; Reichert, J.; Walsh, C. T. Biosynthesis of the Escherichia Coli Siderophore Enterobactin: Sequence of the entF Gene, Expression and Purification of EntF, and Analysis of Covalent Phosphopantetheine. Biochemistry 1991, 30, 2916–2927.

(29) Offret, C.; Desriac, F.; Le Chevalier, P.; Mounier, J.; Jégou, C.; Fleury, Y. Spotlight on Antimicrobial Metabolites from the Marine Bacteria Pseudoalteromonas: Chemodiversity and Ecological Significance. Mar Drugs 2016, 14.

(30) Busch, J. The Diversity, Distribution, and Biological Activity of Brominated Natural Products in the Genus Pseudoalteromonas. Doctoral dissertation, University of California San Diego, 2018.

(31) De Rond, T.; Stow, P.; Eigl, I.; Johnson, R. E.; Chan, L. J. G.; Goyal, G.; Baidoo, E. E. K.; Hillson, N. J.; Petzold, C. J.; Sarpong, R.; et al. Oxidative Cyclization of Prodigiosin by an Alkylglycerol Monooxygenase-like Enzyme. Nat Chem Biol 2017, 13, 1155–1157.

(32) Agarwal, V.; El Gamal, A. A.; Yamanaka, K.; Poth, D.; Kersten, R. D.; Schorn, M.; Allen, E. E.; Moore, B. S. Biosynthesis of Polybrominated Aromatic Organic Compounds by Marine Bacteria. Nat Chem Biol 2014, 10, 640–647.

(33) Ross, A. C.; Gulland, L. E. S.; Dorrestein, P. C.; Moore, B. S. Targeted Capture and Heterologous Expression of the Pseudoalteromonas Alterochromide Gene Cluster in Escherichia Coli Represents a Promising Natural Product Exploratory Platform. ACS Synth Biol 2015, 4, 414–420.

(34) Herz, S.; Eberhardt, S.; Bacher, A. Biosynthesis of Riboflavin in Plants. The ribA Gene of Arabidopsis Thaliana Specifies a Bifunctional GTP Cyclohydrolase II/3,4-dihydroxy-2-butanone 4-phosphate Synthase. Phytochemistry 2000, 53, 723–731.

(35) Schulman, B. A.; Harper, J. W. Ubiquitin-like Protein Activation by E1 Enzymes: The Apex for Downstream Signalling Pathways. Nat Rev Mol Cell Biol 2009, 10, 319–331.

(36) Miyauchi, K.; Kimura, S.; Suzuki, T. A Cyclic Form of N6-threonylcarbamoyladenosine as a Widely Distributed tRNA Hypermodification. Nat Chem Biol 2013, 9, 105–111.

(37) Regni, C. A.; Roush, R. F.; Miller, D. J.; Nourse, A.; Walsh, C. T.; Schulman, B. A. How the MccB Bacterial Ancestor of Ubiquitin E1 Initiates Biosynthesis of the Microcin C7 Antibiotic. EMBO J 2009, 28, 1953–1964.

(38) Ghodge, S. V.; Biernat, K. A.; Bassett, S. J.; Redinbo, M. R.; Bowers, A. A. Post-translational Claisen Condensation and Decarboxylation En Route to the Bicyclic Core of Pantocin A. J Am Chem Soc 2016, 138, 5487–5490.

(39) Xi, J.; Ge, Y.; Kinsland, C.; McLafferty, F. W.; Begley, T. P. Biosynthesis of the Thiazole Moiety of Thiamin in Escherichia Coli: Identification of an Acyldisulfide-linked Protein--protein Conjugate That Is Functionally Analogous to the ubiquitin/E1 Complex. Proc Natl Acad Sci U S A 2001, 98, 8513–8518.

(40) Godert, A. M.; Jin, M.; McLafferty, F. W.; Begley, T. P. Biosynthesis of the Thioquinolobactin Siderophore: An Interesting Variation on Sulfur Transfer. J Bacteriol 2007, 189, 2941–2944.

(41) Akiva, E.; Copp, J. N.; Tokuriki, N.; Babbitt, P. C. Evolutionary and Molecular Foundations of Multiple Contemporary Functions of the Nitroreductase Superfamily. Proc Natl Acad Sci U S A 2017, 114, E9549–E9558.

(42) Mondal, S.; Raja, K.; Schweizer, U.; Mugesh, G. Chemistry and Biology in the Biosynthesis and Action of Thyroid Hormones. Angew Chem Int Ed Engl 2016, 55, 7606–7630.

(43) Schneider, T. L.; Shen, B.; Walsh, C. T. Oxidase Domains in Epothilone and Bleomycin Biosynthesis: Thiazoline to Thiazole Oxidation During Chain Elongation. Biochemistry 2003, 42, 9722–9730.

(44) Melby, J. O.; Li, X.; Mitchell, D. A. Orchestration of Enzymatic Processing by Thiazole/oxazole-modified Microcin Dehydrogenases. Biochemistry 2014, 53, 413–422.

(45) Taga, M. E.; Larsen, N. A.; Howard-Jones, A. R.; Walsh, C. T.; Walker, G. C. BluB Cannibalizes Flavin to Form the Lower Ligand of Vitamin B12. Nature 2007, 446, 449–453.

(46) Bashiri, G.; Rehan, A. M.; Sreebhavan, S.; Baker, H. M.; Baker, E. N.; Squire, C. J. Elongation of the Poly-γ-glutamate Tail of F420 Requires Both Domains of the F420:γ-glutamyl Ligase (FbiB) of Mycobacterium Tuberculosis. J Biol Chem 2016, 291, 6882–6894.

(47) Bashiri, G.; Antoney, J.; Jirgis, E. N. M.; Shah, M. V.; Ney, B.; Copp, J.; Stuteley, S. M.; Sreebhavan, S.; Palmer, B.; Middleditch, M.; et al. A Revised Biosynthetic Pathway for the Cofactor F420 in Prokaryotes. Nat Commun 2019, 10, 1558.

(48) Braga, D.; Hasan, M.; Kröber, T.; Last, D.; Lackner, G. Redox Coenzyme F420 Biosynthesis in Thermomicrobia Involves Reduction by Stand-Alone Nitroreductase Superfamily Enzymes. Appl Environ Microbiol 2020, 86.

(49) Gondry, M.; Lautru, S.; Fusai, G.; Meunier, G.; Ménez, A.; Genet, R. Cyclic Dipeptide Oxidase from Streptomyces Noursei. Isolation, Purification and Partial Characterization of a Novel, Amino Acyl Alpha,beta-dehydrogenase. Eur J Biochem 2001, 268, 1712–1721.

(50) Zenno, S.; Saigo, K.; Kanoh, H.; Inouye, S. Identification of the Gene Encoding the Major NAD(P)H-flavin Oxidoreductase of the Bioluminescent Bacterium Vibrio Fischeri ATCC 7744. J Bacteriol 1994, 176, 3536–3543.

(51) Fisk, J. S.; Mosey, R. A.; Tepe, J. J. The Diverse Chemistry of oxazol-5-(4H)-ones. Chem Soc Rev 2007, 36, 1432–1440.

(52) Deering, R. W.; Chen, J.; Sun, J.; Ma, H.; Dubert, J.; Barja, J. L.; Seeram, N. P.; Wang, H.; Rowley, D. C. N-Acyl Dehydrotyrosines, Tyrosinase Inhibitors from the Marine Bacterium Thalassotalea Sp. PP2-459. J Nat Prod 2016, 79, 447–450.

(53) Guella, G.; N’Diaye, I.; Fofana, M.; Mancini, I. Isolation, Synthesis and Photochemical Properties of Almazolone, a New Indole Alkaloid from a Red Alga of Senegal. Tetrahedron 2006, 62, 1165–1170.

(54) Seyedsayamdost, M.R. High-throughput Platform for the Discovery of Elicitors of Silent Bacterial Gene Clusters. Proc Natl Acad Sci U S A 2014, 111, 7266–7271.

(55) Walsh, C.; Wencewicz, T. Antibiotics: Challenges, Mechanisms, Opportunities; 2nd ed.; American Society of Microbiology: Washington, DC, 2016.

(56) Skinnider, M. A.; Merwin, N. J.; Johnston, C. W.; Magarvey, N. A. PRISM 3: Expanded Prediction of Natural Product Chemical Structures from Microbial Genomes. Nucleic Acids Res 2017, 45, W49–W54.

(57) Blin, K.; Shaw, S.; Steinke, K.; Villebro, R.; Ziemert, N.; Lee, S. Y.; Medema, M. H.; Weber, T. antiSMASH 5.0: Updates to the Secondary Metabolite Genome Mining Pipeline. Nucleic Acids Res 2019, 47, W81–W87.

(58) Navarro-Muñoz, J. C.; Selem-Mojica, N.; Mullowney, M. W.; Kautsar, S. A.; Tryon, J. H.; Parkinson, E. I.; De Los Santos, E. L. C.; Yeong, M.; Cruz-Morales, P.; Abubucker, S.; et al. A Computational Framework to Explore Large-scale Biosynthetic Diversity. Nat Chem Biol 2020, 16, 60–68.

(59) Hon, J.; Borko, S.; Stourac, J.; Prokop, Z.; Zendulka, J.; Bednar, D.; Martinek, T.; Damborsky, J. EnzymeMiner: Automated Mining of Soluble Enzymes with Diverse Structures, Catalytic Properties and Stabilities. Nucleic Acids Res 2020, 48, W104–W109.

(60) Gerlt, J.A. Genomic Enzymology: Web Tools for Leveraging Protein Family Sequence-function Space and Genome Context to Discover Novel Functions. Biochemistry 2017, 56, 4293–4308.

(61) Wuchty, S.; Almaas, E. Evolutionary Cores of Domain Co-occurrence Networks. BMC Evol Biol 2005, 5, 24.

(62) Wang, Z.; Zhang, X. -C.; Le, M. H.; Xu, D.; Stacey, G.; Cheng, J. A Protein Domain Co-occurrence Network Approach for Predicting Protein Function and Inferring Species Phylogeny. PLoS ONE 2011, 6, e17906.

(63) Barrera, A.; Alastruey-Izquierdo, A.; Martín, M. J.; Cuesta, I.; Vizcaíno, J. A. Analysis of the Protein Domain and Domain Architecture Content in Fungi and Its Application in the Search of New Antifungal Targets. PLoS Comput Biol 2014, 10, e1003733.

(64) Suhre, K. Inference of Gene Function Based on Gene Fusion Events: The Rosetta-stone Method. Methods Mol Biol 2007, 396, 31–41.

(65) Promponas, V. J.; Ouzounis, C. A.; Iliopoulos, I. Experimental Evidence Validating the Computational Inference of Functional Associations from Gene Fusion Events: a Critical Survey. Brief Bioinformatics 2014, 15, 443–454.

(66) Alcántara, R.; Onwubiko, J.; Cao, H.; Matos, P. de; Cham, J. A.; Jacobsen, J.; Holliday, G. L.; Fischer, J. D.; Rahman, S. A.; Jassal, B.; et al. The EBI Enzyme Portal. Nucleic Acids Res 2013, 41, D773–80.

(67) Huang, D. W.; Sherman, B. T.; Lempicki, R. A. Systematic and Integrative Analysis of Large Gene Lists Using DAVID Bioinformatics Resources. Nat Protoc 2009, 4, 44–57.

(68) George, R. A.; Spriggs, R. V.; Thornton, J. M.; Al-Lazikani, B.; Swindells, M. B. SCOPEC: a Database of Protein Catalytic Domains. Bioinformatics 2004, 20 Suppl 1, i130–6.

(69) Alborzi, S. Z.; Devignes, M. -D.; Ritchie, D. W. ECDomainMiner: Discovering Hidden Associations Between Enzyme Commission Numbers and Pfam Domains. BMC Bioinformatics 2017, 18, 107.

(70) Van Keulen, G.; Dyson, P. J. Production of Specialized Metabolites by Streptomyces Coelicolor A3(2). Adv Appl Microbiol 2014, 89, 217–266.

(71) Leisinger, T.; Margraff, R. Secondary Metabolites of the Fluorescent Pseudomonads. Microbiol Rev 1979, 43, 422–442.

(72) Jensen, P. R.; Moore, B. S.; Fenical, W. The Marine Actinomycete Genus Salinispora: a Model Organism for Secondary Metabolite Discovery. Nat Prod Rep 2015, 32, 738–751.

(73) De Castro, P. P.; Carpanez, A. G.; Amarante, G. W. Azlactone Reaction Developments. Chem. Eur. J 2016, 22, 10294–10318.

(74) Marra, I. F. S.; de Castro, P. P.; Amarante, G. W. Recent Advances in Azlactone Transformations. European J Org Chem 2019, 2019, 5830–5855.

(75) Kenney, G. E.; Dassama, L. M. K.; Pandelia, M.-E.; Gizzi, A. S.; Martinie, R. J.; Gao, P.; DeHart, C. J.; Schachner, L. F.; Skinner, O. S.; Ro, S. Y.; et al. The Biosynthesis of Methanobactin. Science 2018, 359, 1411–1416.

(76) Cappadocia, L.; Lima, C. D. Ubiquitin-like Protein Conjugation: Structures, Chemistry, and Mechanism. Chem Rev 2018, 118, 889–918.

(77) Blattner, F. R.; Plunkett, G.; Bloch, C. A.; Perna, N. T.; Burland, V.; Riley, M.; Collado-Vides, J.; Glasner, J. D.; Rode, C. K.; Mayhew, G. F.; et al. The Complete Genome Sequence of Escherichia Coli K-12. Science 1997, 277, 1453–1462.

(78) Xie, B.-B.; Shu, Y.-L.; Qin, Q.-L.; Rong, J.-C.; Zhang, X.-Y.; Chen, X.-L.; Zhou, B.-C.; Zhang, Y.-Z. Genome Sequence of the Cycloprodigiosin-producing Bacterial Strain Pseudoalteromonas Rubra ATCC 29570(T). J Bacteriol 2012, 194, 1637–1638.

(79) Shannon, P.; Markiel, A.; Ozier, O.; Baliga, N. S.; Wang, J. T.; Ramage, D.; Amin, N.; Schwikowski, B.; Ideker, T. Cytoscape: a Software Environment for Integrated Models of Biomolecular Interaction Networks. Genome Res 2003, 13, 2498–2504.

(80) Katoh, K.; Rozewicki, J.; Yamada, K. D. MAFFT Online Service: Multiple Sequence Alignment, Interactive Sequence Choice and Visualization. Brief Bioinformatics 2017, bbx108.

