## Supporting Information for "Co-Occurrence of Enzyme Domains Guides the Discovery of an Oxazolone Synthetase"

Other supplementary materials for this manuscript include the following:

**Dataset S1:** co-occurring_enzyme_domains.cys
A Cytoscape session file containing the directed CO-ED networks depicted in Figures S1-5 and S14a. Generated in Cytoscape 3.8

### 1 Supplementary Figures

#### 1.1 Figure S1: Multi-domain enzymes are over-represented among enzymes involved in secondary metabolism

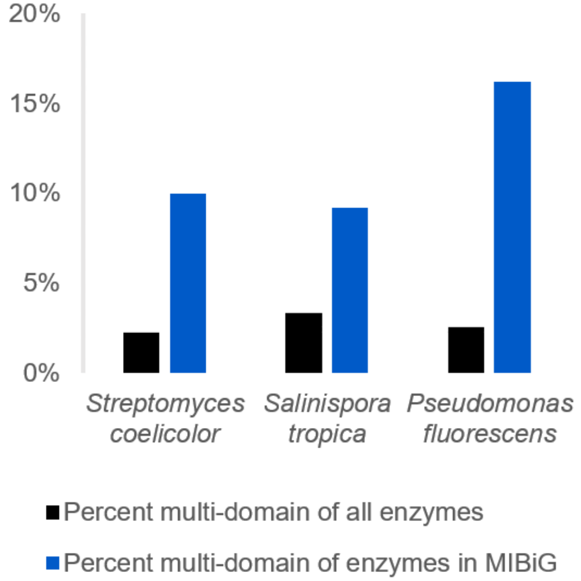

Multi-domain enzymes are over-represented among enzymes known to be involved in secondary metabolism - i.e., those contained in MIBiG[1] - compared to all enzymes, in three organisms well-represented in MIBiG: *S. coelicolor* A3(2) (NCBI ASM20383v1[2], taxid 100226), *S. tropica* CNB-440 (ASM1642v1[3], taxid 369723), and *P. fluorescens* Pf-5 (ASM1226v1[4], taxid 220664)

#### 1.2 Figure S2: Legend for CO-ED networks in Supplementary Figures

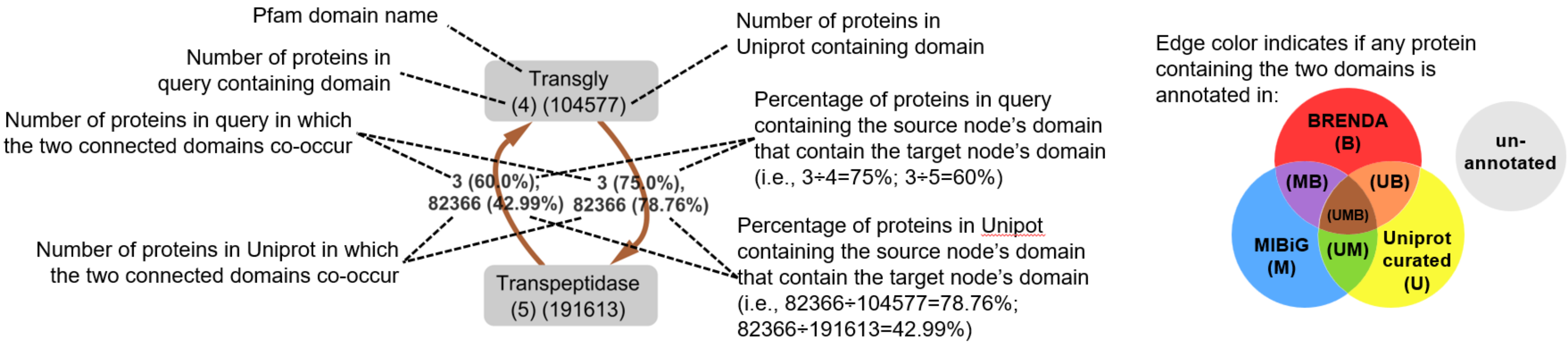

The CO-ED networks shown in the Supplementary Figures are depicted as directed graphs: for each connected pair of domains there are two edges, unlike the networks in the main text which are undirected graphs. The premise is the same: If two domains co-occur in a protein in the query, they are connected by edges. Using two edges per domain pair allows us to display what percentage of proteins containing one domain also contain the other domain. Also included is the frequency of the domains and domain pairs in the Uniprot[5] database. These pieces of information can help the user gauge how "rare" a domain pair is, which can help prioritize proteins for genome mining.

The networks were generated using the Jupyter notebook available at https://github.com/tderond/CO-ED, and visualized in Cytoscape[6]. We strongly encourage the reader to explore these networks interactively as Dataset S1 in Cytoscape, which allows one to re-arrange the nodes, change formatting settings, and see the long-form descriptions annotated for each Pfam domain

For all networks in the Supplementary Figures, only the connected component is shown.

#### 1.3 Figure S3: CO-ED network for *E. coli* K12

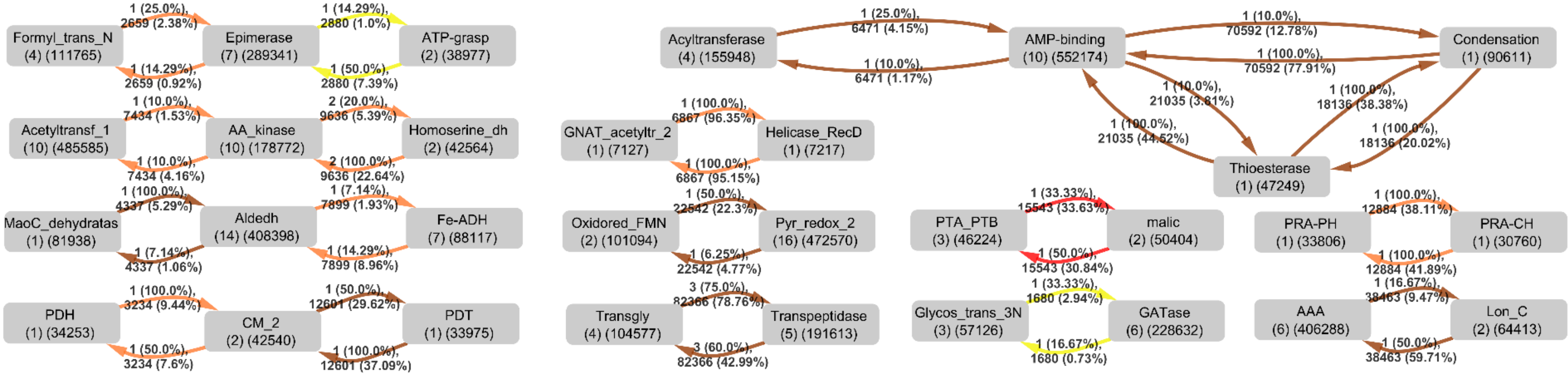

Based on the NCBI ASM584v2[7] genome assembly. Please refer to Fig. S2 above for the legend.

#### 1.4 Figure S4: CO-ED network for *P. rubra* DSM 6842

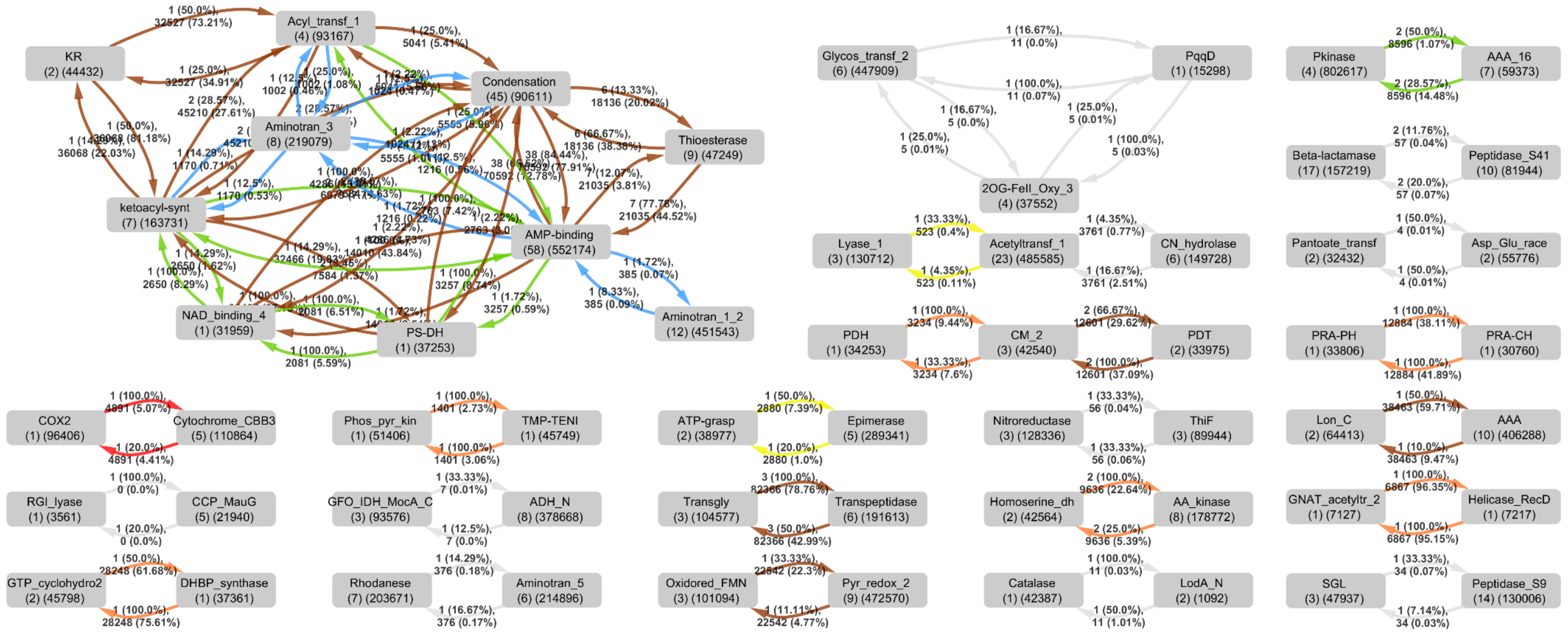

Based on the NCBI "Prub_2.0"[8] genome assembly. Please refer to Fig. S2 above for the legend.

#### 1.5 Figure S5: Phylogeny of organisms that harbor OxzB

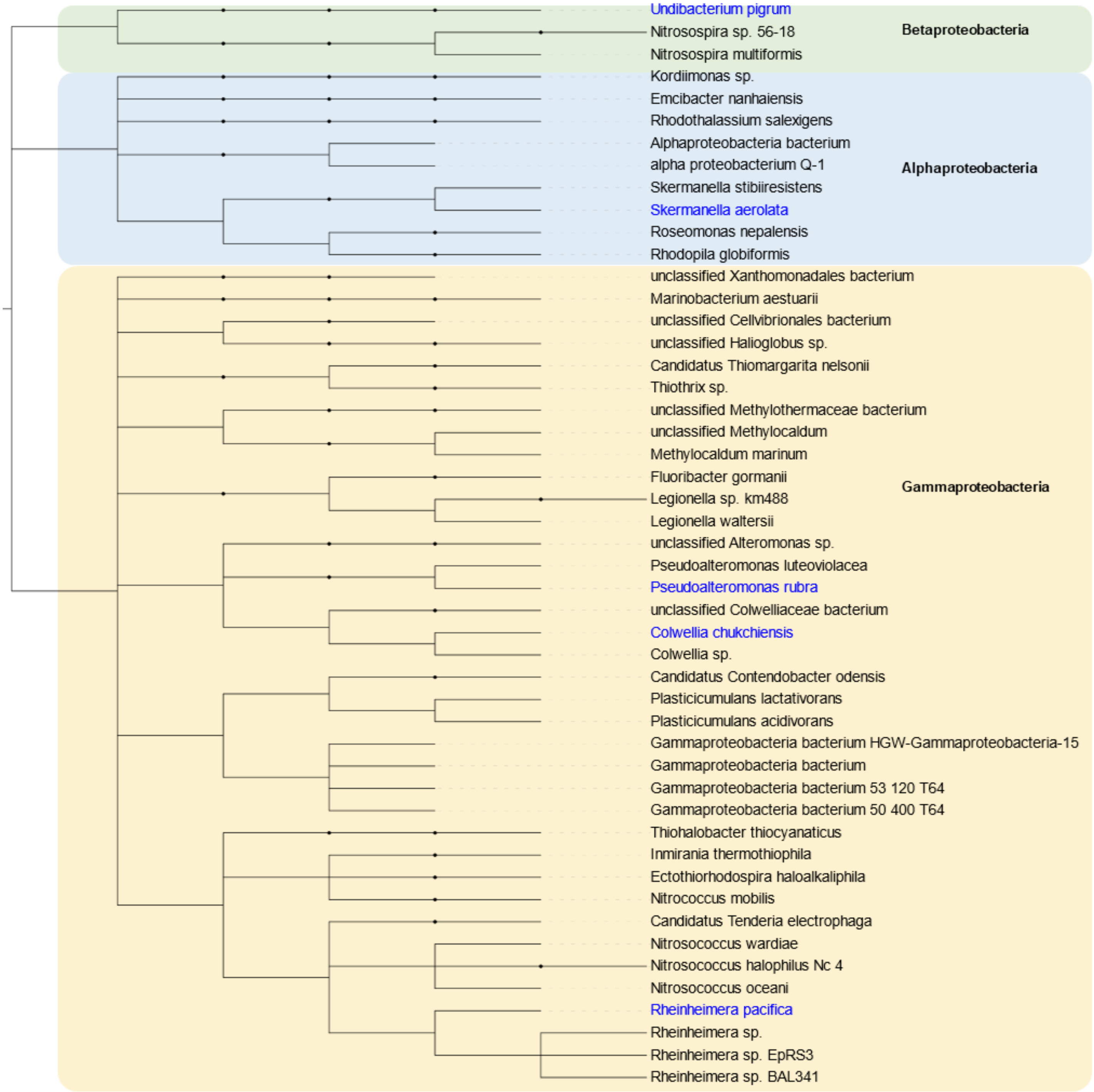

Phylogenetic distribution of organisms harboring OxzB-like ThiF-Nitroreductase di-domain proteins contained in the Uniprot database. Taxonomy was based on NCBI taxonomy. Unassigned metagenomic contigs not shown. Species names written in blue were those whose *oxzAB* clusters were investigated heterologously in this study.

#### 1.6 Figure S6: Color properties of the metabolic products of *oxzAB*

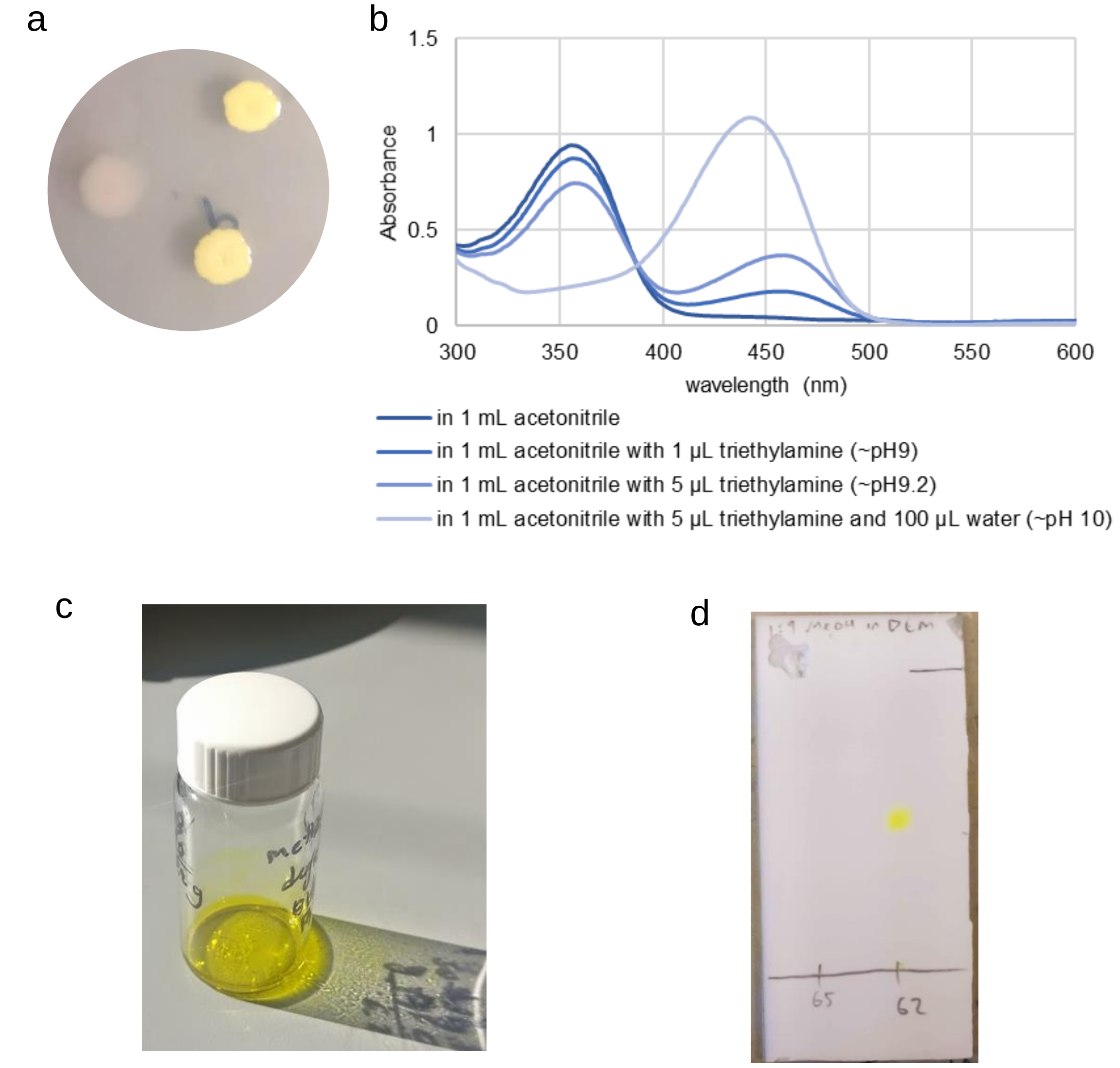

Color properties of the metabolic products of *oxzAB*.

a) *E. coli* colonies containing pTOPO-Pr_*oxzAB* turn yellow after a few days of growth on LB agar. No inducer is added to the media, so presumably the production is due to leaky expression of *oxzAB*.
b) UV/vis spectrum of an ethyl acetate extract of E. *coli* BLR(DE3)/pTOPO-Pr_*oxzAB* dried down and re-dissolved in 1mL acetonitrile, upon addition of 1 µL of triethylamine to this solution, upon the addition for an additional 4 µL to this solution, and upon the addition of water to a final concentration of 10% to this solution.
c) Purified nonyltyrazolone in methanol/$K_2CO_3$
d) Ethyl acetate extract of E. *coli* BLR(DE3)/pTOPO-Pr_*oxzAB* (right lane) and *E. coli* BLR(DE3) harboring an empty plasmid (left lane) developed in 1:9 methanol:DCM and stained with 100mM NaOH in ethanol. The yellow spot is visible for about a minute

#### 1.7 Figure S7: Degradation of the metabolic products of *oxzAB*

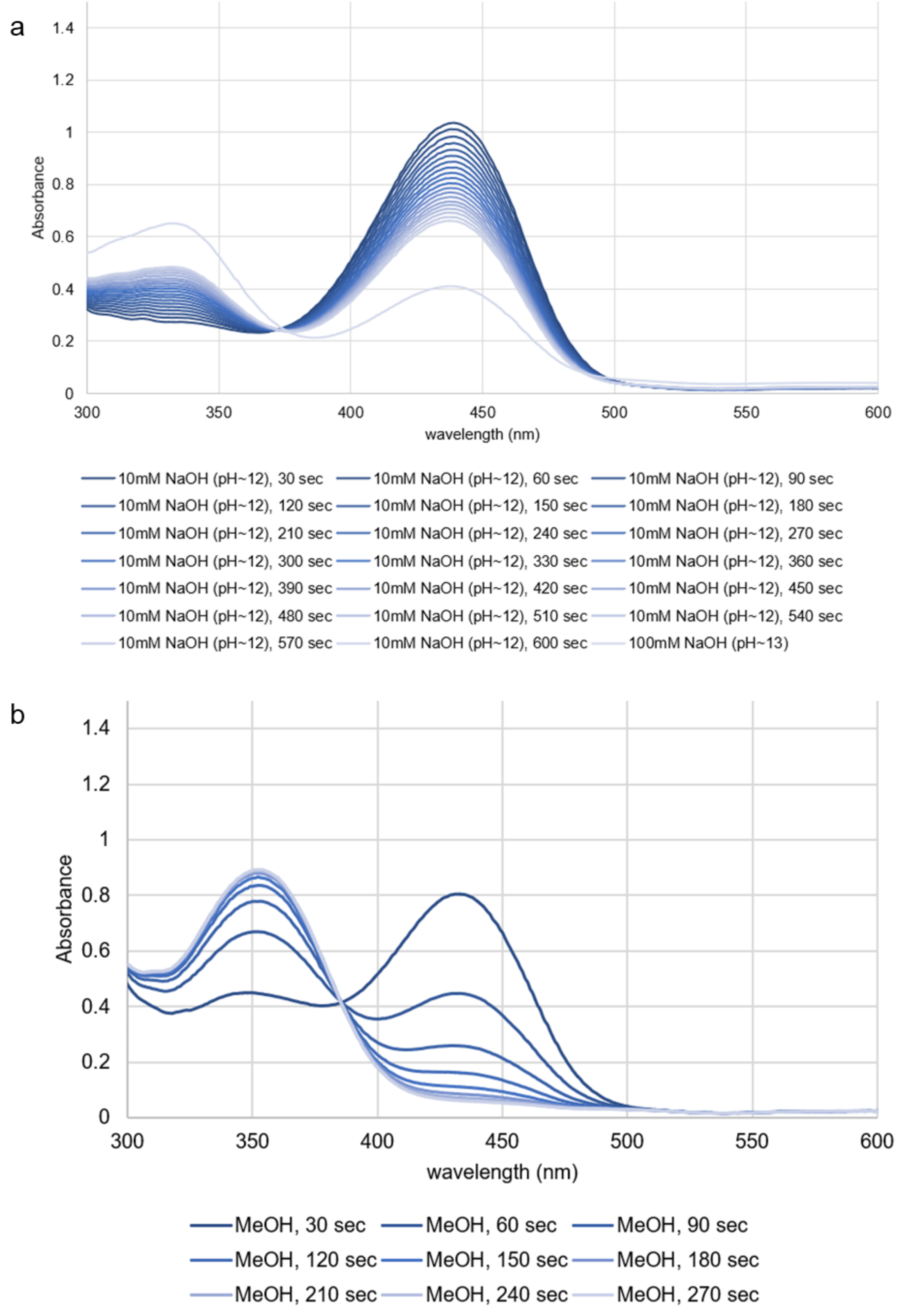

Degradation of an ethyl acetate extract of *E. coli* BLR(DE3)/pTOPO-Pr_*oxzAB* incubated in (**a**) aqueous sodium hydroxide, or (**b**) methanol

#### 1.8 Figure S8: Inclusion of genes up- or downstream from *oxzAB*

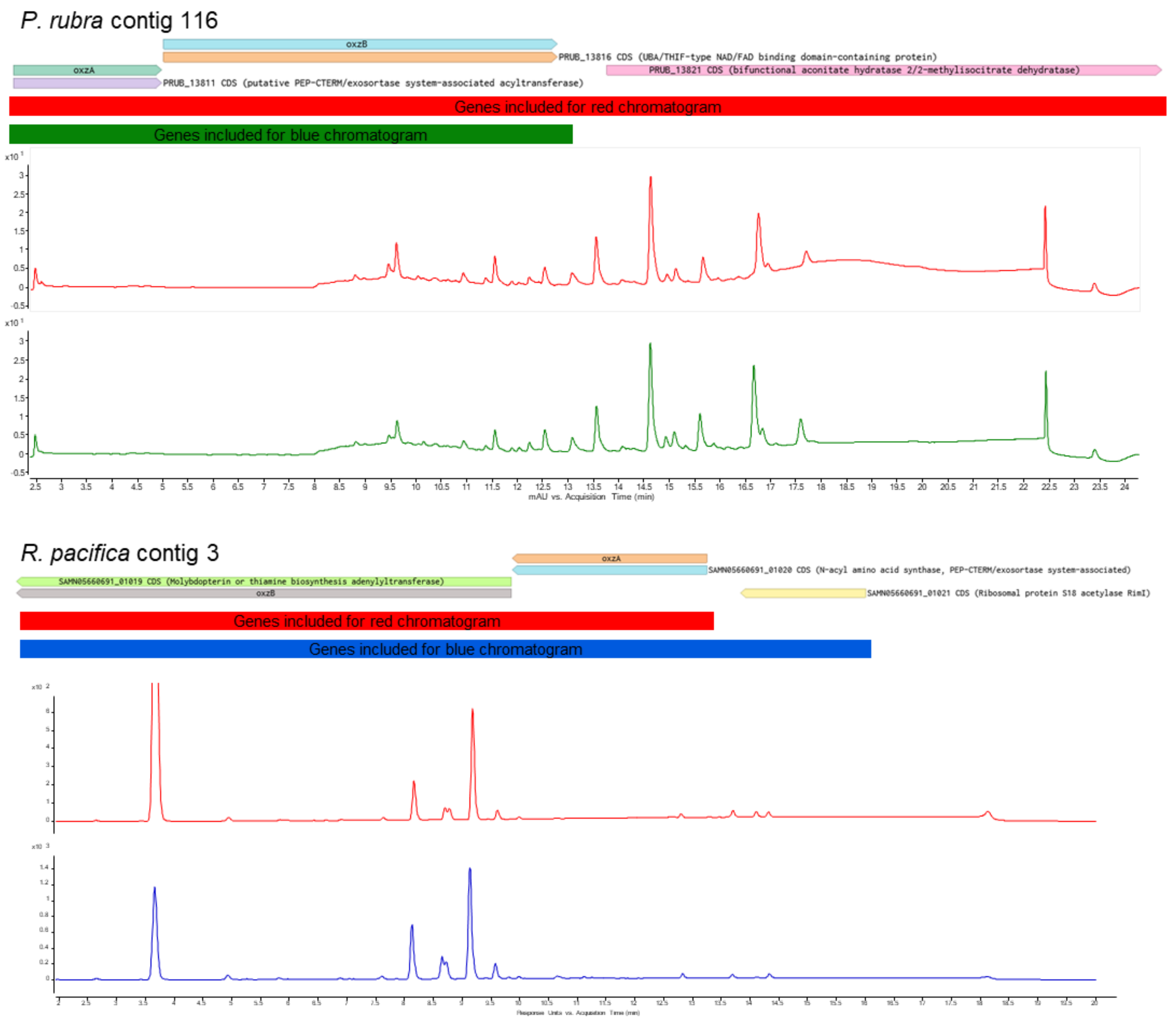

Inclusion of genes up- or downstream from *oxzAB* during heterologous expression in E. coli does not result in an altered metabolite profile. In P. rubra, an aconitase-like gene is downstream of *oxzAB*, while in *R. pacifica* an acyltransferase-like gene is upstream of *oxzAB*.

Note: while the *R. pacifica* chromatograms shown use the same column and gradient as main text Figure 4, but the *P. rubra* chromatograms were acquired before these conditions were established. Instead, the following chromatographic conditions were used: Phenomenex Luna C18(2) (100x4.6mm, 5um pore size), solvent A: water + 0.1% FA, solvent B: acetonitrile + 0.1% FA. gradient 5% B for 5 minutes, ramp to 100% B in 10 minutes, 100% B for 5 minutes, ramp to 5% in 1 minute, 5% B for 5 minutes. The *P. rubra* samples were extracted in methanol, leading to partial degradation of the oxazolones, and thus the relatively complex chromatogram observed.

#### 1.9 Figure S9: Key NMR correlations used for structure elucidation

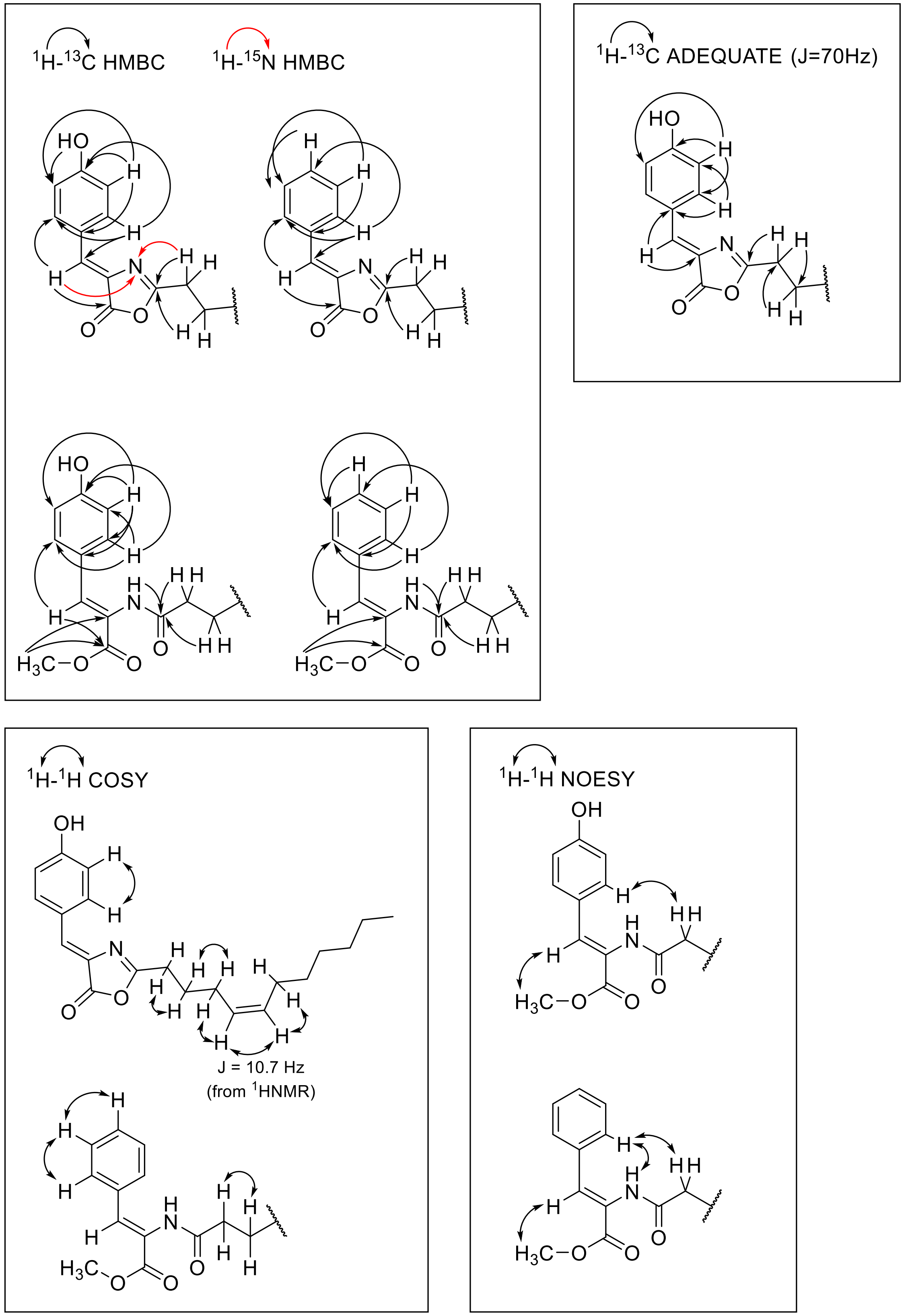

Key NMR correlations used to elucidate the structures of the tyrazolones and phenazolones. *E*/*Z* isomers were assigned by NOESY on their respective ring-opened methanol adducts and assumes the alkene stereochemistry is retained during the methanol addition process (see Fig. S10), and by comparing chemical shifts reported for closely related compounds.

#### 1.10 Figure S10: Adducts formed when incubating oxazolones with methanol under basic conditions

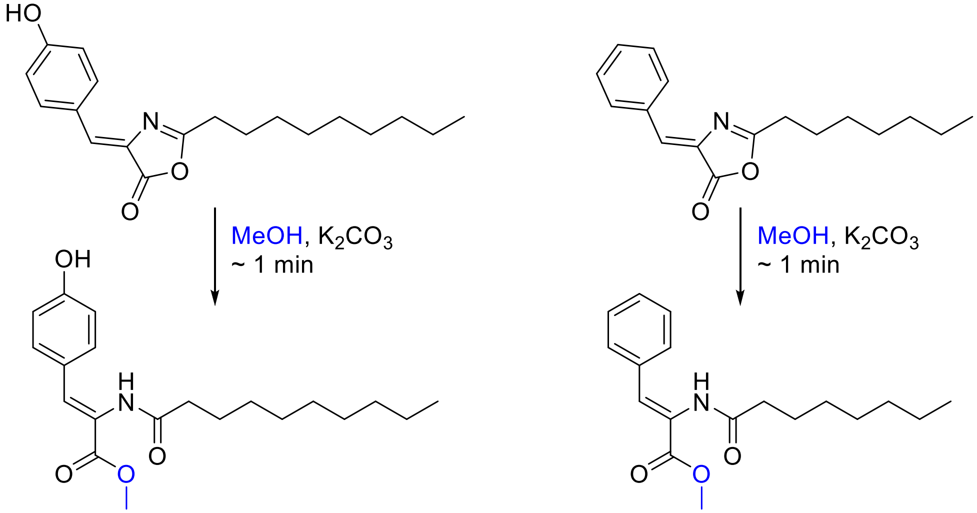

Adducts formed when incubating tyrazolones and phenazolones with methanol under basic conditions. For the purposes of stereochemical assessment of the oxazolones, we assume the alkene stereochemistry is retained during this reaction

#### 1.11 Figure S11: Possible orders of reactions in OxzAB-catalyzed tyrazolone production

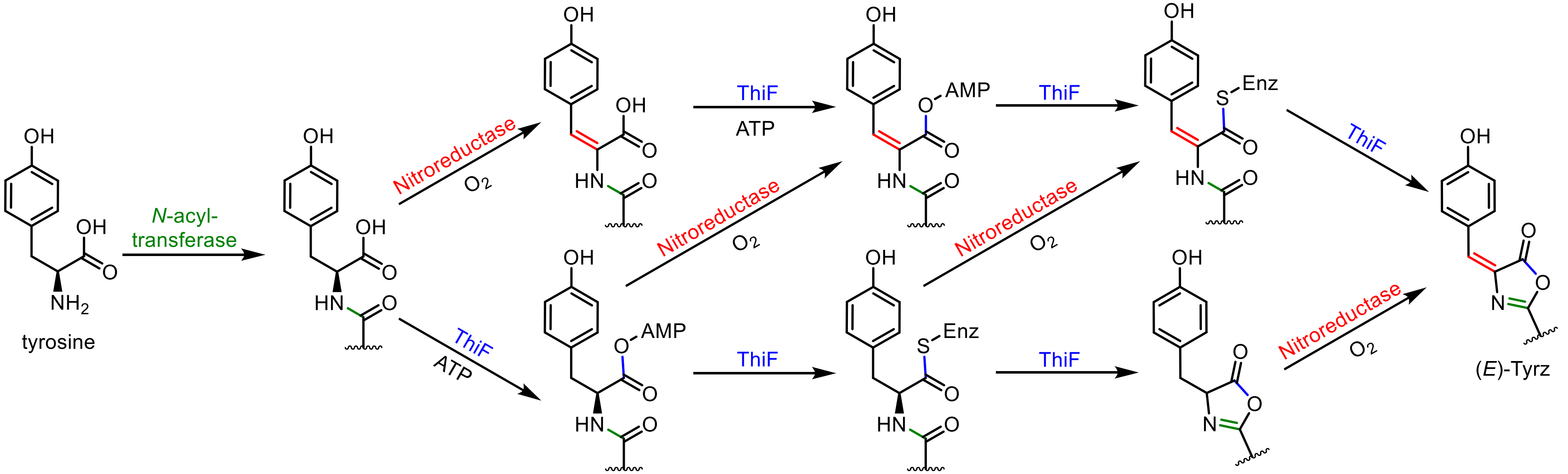

Possible orders of reactions in the OxzAB-catalyzed production of tyrazolone. Upon production, (*E*)-Tyrz will start isomerizing into (*Z*)-Tyrz, forming an equilibrium.

#### 1.12 Figure S12: (*E*)/(*Z*) ratio of OxzB products

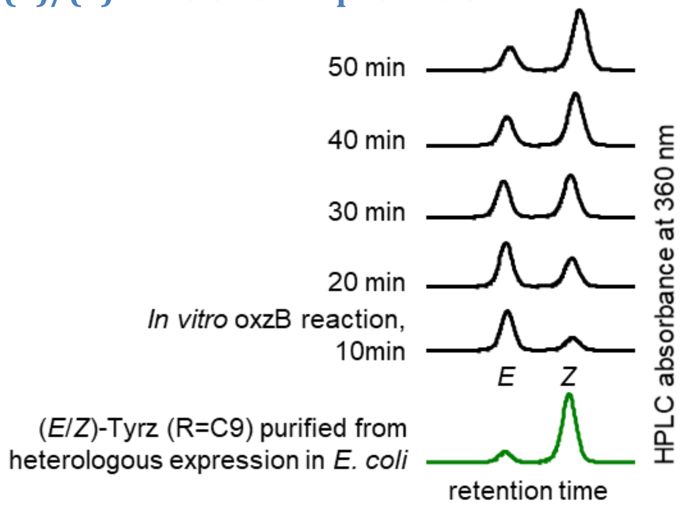

Analysis of the stereochemical ratio of nonyltyrazolone OxzB products. Since the extinction coefficients of the isomers are not known, the relative peak sizes are not necessarily indicative of true product ratios. The chromatograms shown here all derive from a single enzyme reaction, aliquots of which were quenched with acetonitrile at the timepoint indicated, and immediately analyzed by HPLC.

#### 1.13 Figure S13: HPLC analysis of oxazolone production in native hosts

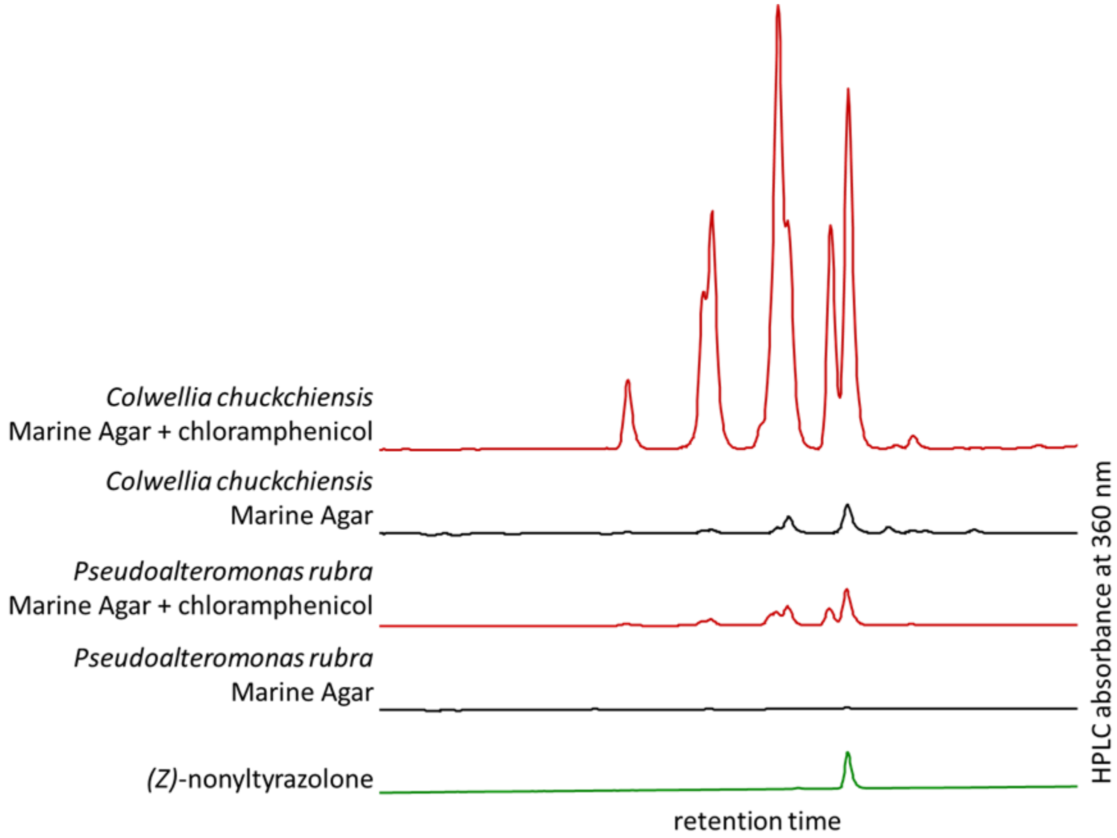

Native production of oxazolones. All Y-axes are on the same scale, indicating the dramatic differences in oxazolone production observed under these different conditions. It appears that by *P. rubra* and *C. chukchiensis* produce, in addition to oxazolones also produced when heterologously expressing *oxzAB* in *E. coli*, oxazolones that were not produced heterologously. Based on the masses and retention times, we suspect these oxazolones incorporate iso- or anteiso- branched fatty acyl chains, which are not natively found in *E. coli*.

### 1.14 Figure S14: CO-ED network for *S. coelicolor* A2(3)

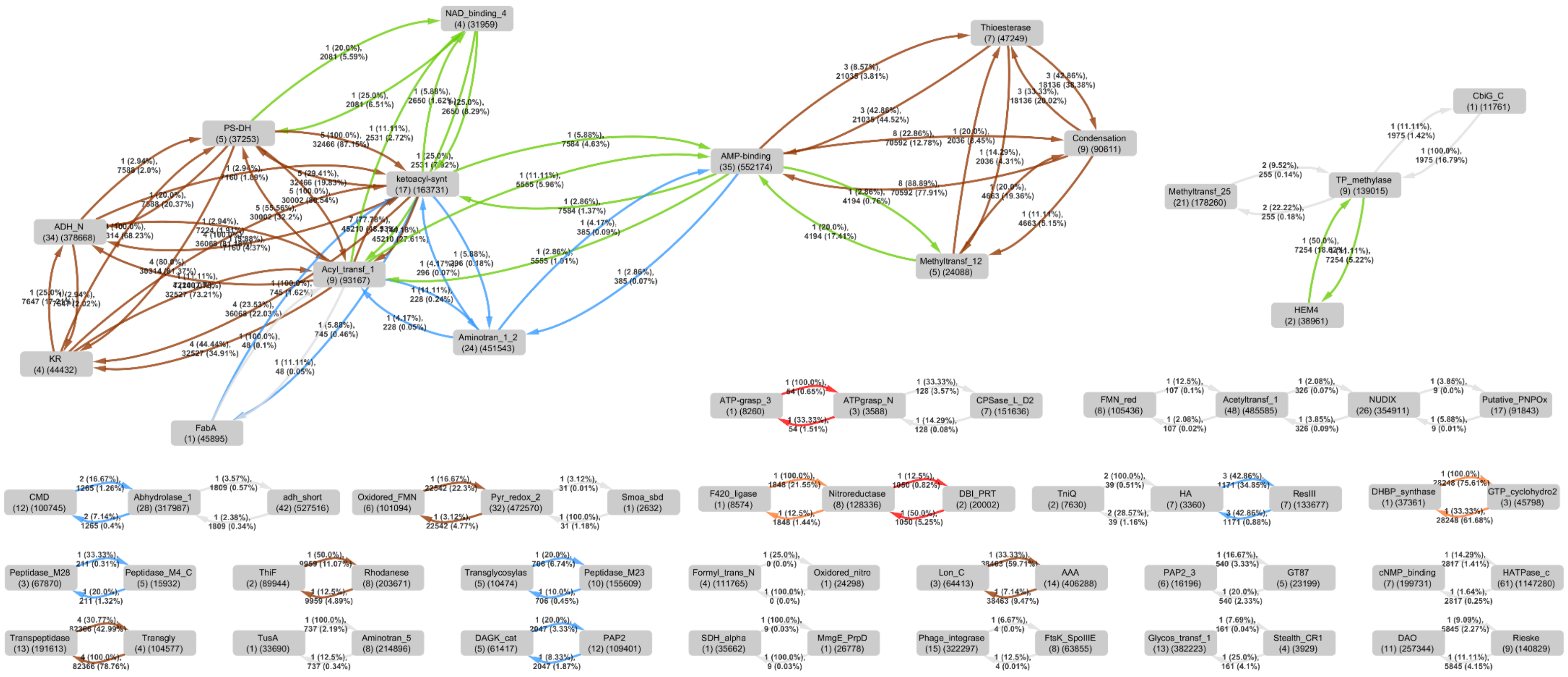

Based on the NCBI ASM20383v1[2] genome assembly. Please refer to Fig. S2 above for the legend.

### 1.15 Figure S15: CO-ED network for *S. tropica* CNB-440

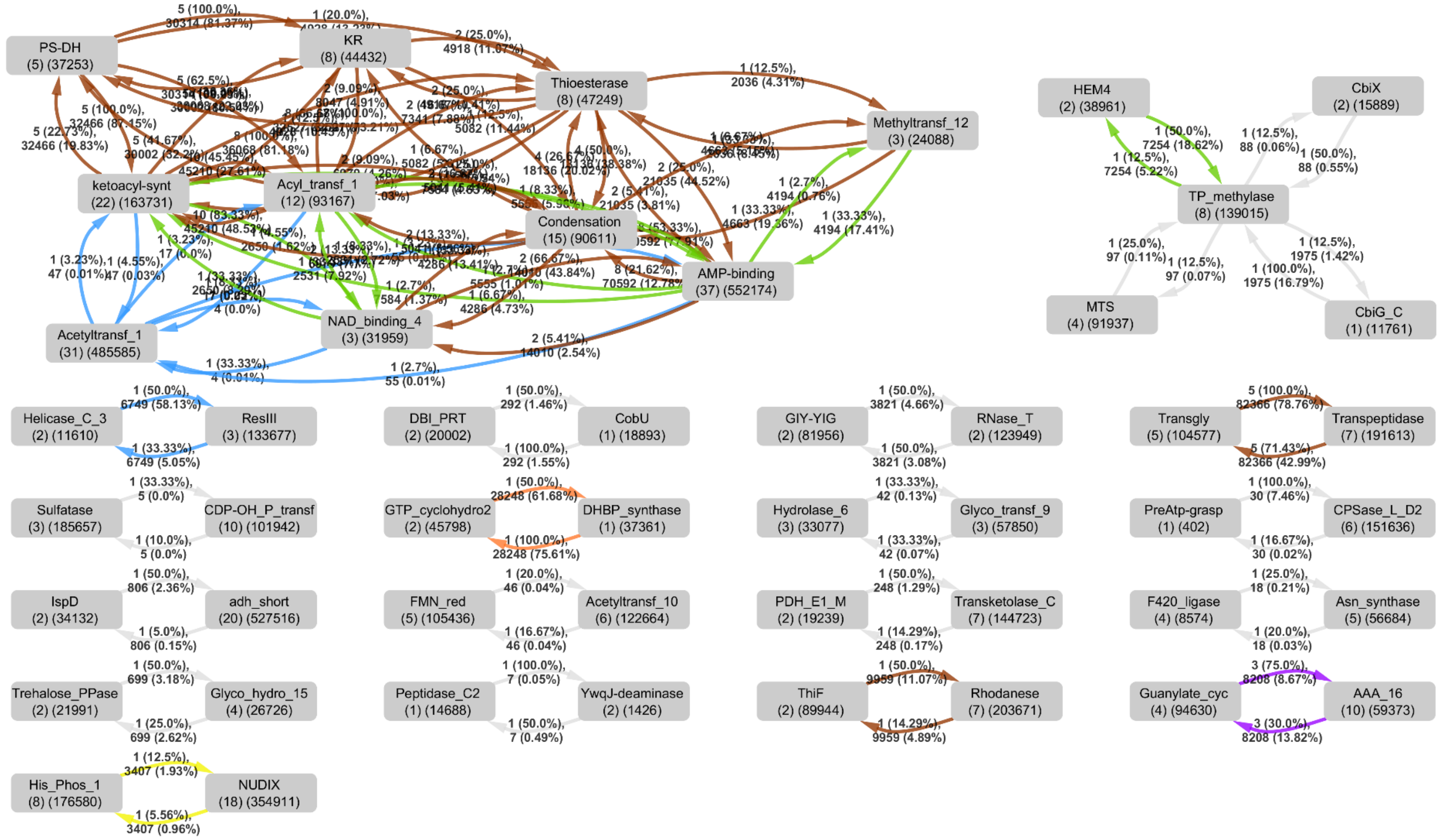

Based on the ASM1642v1[3] genome assembly. Please refer to Fig. S2 above for the legend.

#### 1.16 Figure S16: CO-ED network for *P. fluorescens* Pf-5

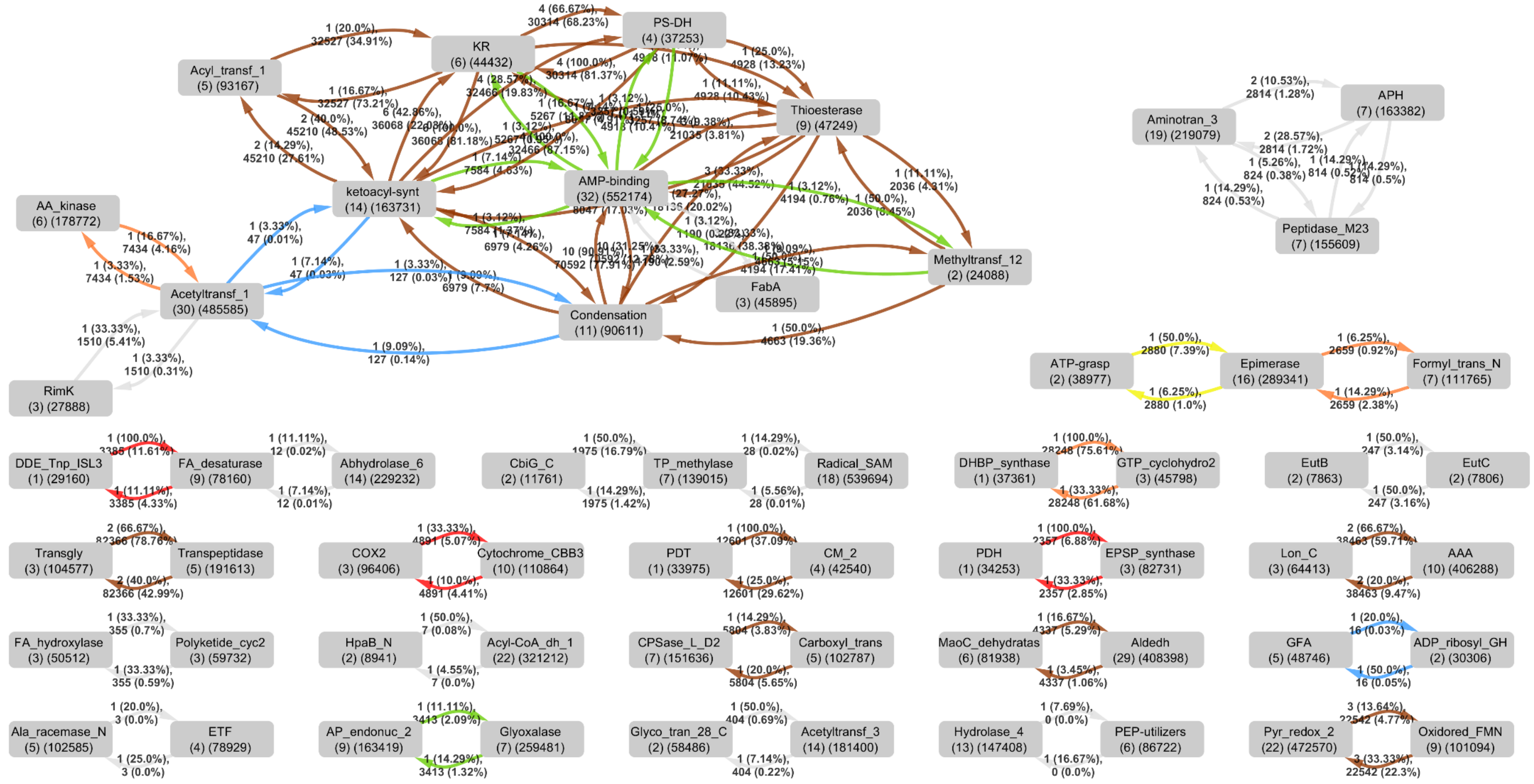

Based on the NCBI ASM1226v1[4] genome assembly. Please refer to Fig. S2 above for the legend.

### Figure S17: CO-ED analysis of all proteins in the Uniprot database

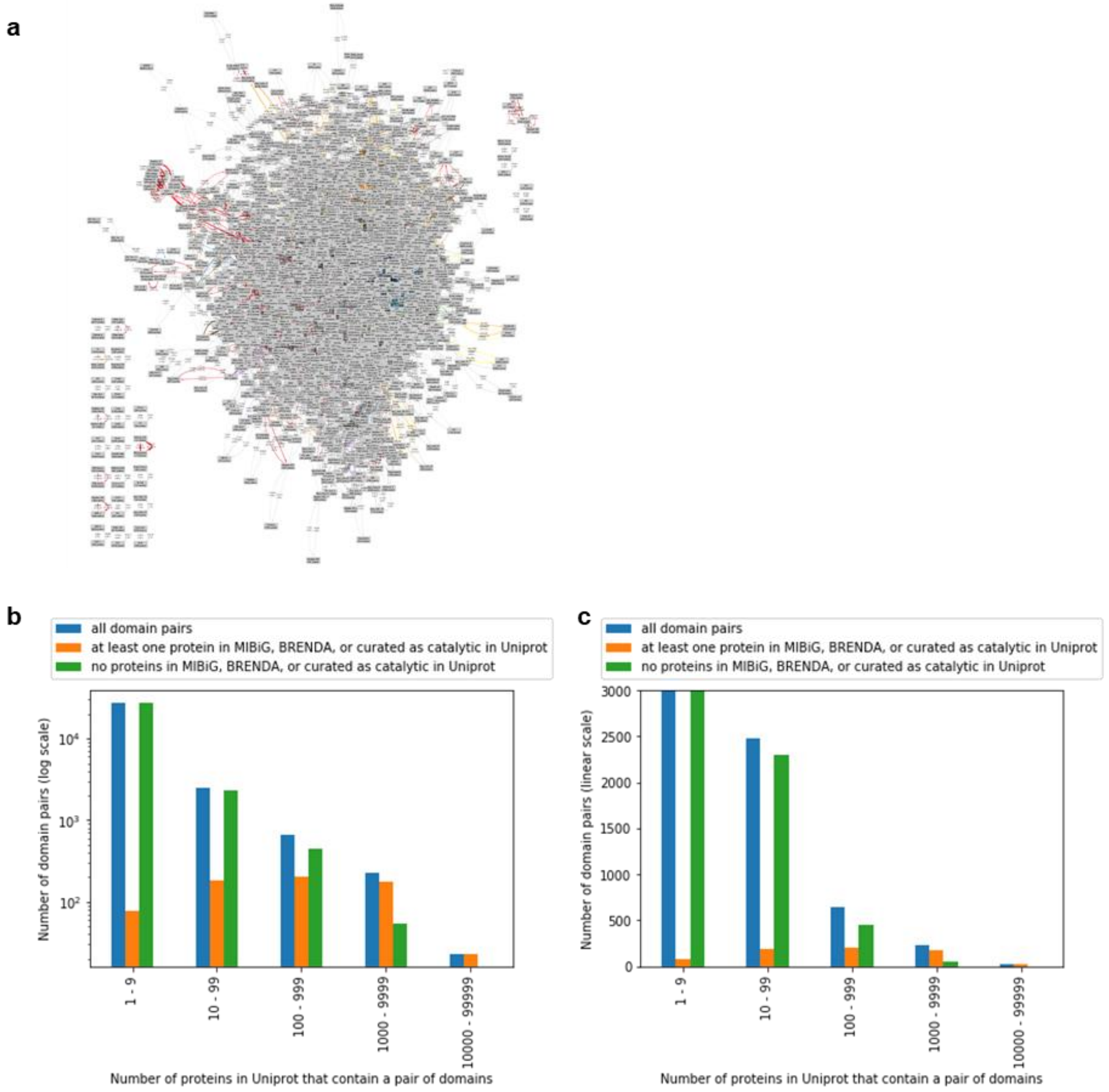

CO-ED analysis of all proteins in the Uniprot database.

**a**: CO-ED network generated from all proteins Uniprot, based on Pfam-A domains pre-annotated in Uniprot. Edges are only drawn for pairs of domains annotated in at least 10 uniprot entries. We strongly encourage the reader to explore these networks interactively as Dataset S1 in Cytoscape, which allows one to re-arrange the nodes, change formatting settings, and see the long-form descriptions annotated for each Pfam domain. Refer to Fig. S2 above for the legend.

**b** and **c** show the same data, but represented on log and linear y axes respectively. The bars show how many domain pairs are pre-annotated in Uniprot entries with a frequency shown on the x axis, and among those pairs, how many have an annotated member in at least one database. Only domains part of the “y”, “m”, “s” and “e” categories (see section 3.1) are considered for the analysis shown, however when the “u” category is included, while the absolute numbers change, the changes in the proportions of domains pairs that are unannotated is negligible.

The network and graphs can be re-generated using the Jupyter notebooks available at https://github.com/tderond/CO-ED

### 2 Supplementary Tables

#### 2.1 Table S1: OxzB-like proteins in Uniprot

"Type of genome sequence" category letters mean:

I Isolate
M Metagenome-Assembled Genome
C Unassigned metagenomic contigs

"Habitat" category letters mean:

M Marine
HT Hydrothermal vent
F Freshwater
W Wastewater treatment plant
HS Hypersaline aqueous
HA Hyperacidic aqueous
E Microbial Fuel Cell
T Terrestrial
A Air

"In cluster with *oxzA*?" letters mean:

"F" *oxzA* immediately upstream, same strand as *oxzB*
"R" *oxzA* immediately upstream, opposite strand as *oxzB*
"downstream F" *oxzA* immediately downstream, same strand as *oxzB*
"X genes back" *oxzA* not immediately upstream or downstream, but nearby

Proteins expressed heterologously in this study are bolded

| Uniprot Entry | Organism | Taxonomic lineage | type of genome sequence | habitat | habitat (details) | citation for sequence | OxzB length (AAs) | in cluster with *oxzA*? |
|---|---|---|---|---|---|---|---|---|
| A0A061QD22 | alpha proteobacterium Q-1 | Alphaproteobacteria | I | HS | natural gas fracking brine water | https://dx.doi.org/10.1128%2FgenomeA.00659-14 | 677 | F |
| **A0A0D2VND3** | **Skermanella aerolata KACC 11604** | **Alphaproteobacteria** | **I** | **A** | **air** | **https://doi.org/10.1099/ijs.0.64676-0** | **712** | **F** |
| A0A2A4V0E4 | Alphaproteobacteria bacterium | Alphaproteobacteria | M | M | subseafloor aquifer | https://dx.doi.org/10.1038/ismej.2017.187 | 671 | F 2 genes back |
| A0A2A5CHV2 | Alphaproteobacteria bacterium | Alphaproteobacteria | M | M | subseafloor aquifer | https://dx.doi.org/10.1038/ismej.2017.187 | 659 | F 2 genes back |
| A0A2E1CY12 | Kordiimonas sp. | Alphaproteobacteria | M | M | ocean | https://dx.doi.org/10.1038/sdata.2017.203 | 683 | F 2 genes back |
| A0A2S6NGC5 | Rhodopila globiformis (Rhodopseudomonas globiformis) | Alphaproteobacteria | I | HA | acidic sulfur sping | https://doi.org/10.1007/BF00446317 | 668 | unclear, on edge of contig |
| A0A3M1A419 | Alphaproteobacteria bacterium | Alphaproteobacteria | M | HT | geothermal spring | http://dx.doi.org/10.1264/jsme2.ME19017 | 685 | F |

| | | | | | | | | |
|---|---|---|---|---|---|---|---|---|
| A0A4R2PGL0 | Rhodothalassium salexigens DSM 2132 | Alphaproteobacteria | I | M | marine | https://doi.org/10.1007/BF00425949 | 682 | F |
| A0A501PH46 | Emcibacter nanhaiensis MCCC 1A06723 | Alphaproteobacteria | I | M | marine sediment | https://doi.org/10.1007/s10482-015-0381-y | 667 | F 2 genes back |
| A0A502GCF0 | Roseomonas nepalensis | Alphaproteobacteria | I | F | permafrost | https://doi.org/10.1111/1462-2920.14715 | 647 | downstream F |
| W9HBK6 | Skermanella stibiiresistens SB22 | Alphaproteobacteria | I | T | coal mine soil | https://doi.org/10.1099/ijs.0.033746-0 | 682 | F |
| A0A1Q4C5S3 | Nitrosospira sp. 56-18 | Betaproteobacteria | | W | Activated Sludge Tailings Effluent Remediation bioreactor | https://doi.org/10.1021/acs.est.6b04477 | 683 | downstream F |
| **A0A318J8Z4** | **Undibacterium pigrum** | **Betaproteobacteria** | **I** | **F** | **drinking water** | **https://doi.org/10.1099/ijs.0.64785-0** | **675** | **F** |
| Q2YC39 | Nitrosospira multiformis (strain ATCC 25196 / NCIMB 11849 / C 71) | Betaproteobacteria | I | T | soil | https://doi.org/10.1007/BF00409115 | 684 | F |
| A0A0S2TIG7 | Candidatus Tenderia electrophaga | Gammaproteobacteria | M | E | microbial fuel cell | https://doi.org/10.1099/ijsem.0.001006 | 682 | R |
| A0A0W1A4Y3 | Legionella waltersii | Gammaproteobacteria | I | F | drinking water | Int J Syst Bacteriol. 1996 Jul;46(3):631-4. | 653 | no |
| A0A0X3Y713 | Rheinheimera sp. EpRS3 | Gammaproteobacteria | I | T | rhizosphere | https://doi.org/10.3389/fmicb.2019.00510 | 655 | F |
| A0A148N8V8 | Methylothermaceae bacteria B42 | Gammaproteobacteria | M | HT | hydrothermal vent | https://dx.doi.org/10.3389/fmicb.2015.01425 | 656 | R |
| A0A166YVV4 | Pseudoalteromonas luteoviolacea DSM 6061 | Gammaproteobacteria | I | M | marine | INTERNATIONAL JOURNAL OF SYSTEMATIC BACTERIOLOGY, Jan. 1982, p. 82-86 | 655 | F |
| A0A1A9EXN0 | Marinobacterium aestuarii | Gammaproteobacteria | I | M | estuary sediment | https://doi.org/10.1099/ijsem.0.002561 | 665 | no |
| **A0A1H6KA59** | **Rheinheimera pacifica** | **Gammaproteobacteria** | **I** | **M** | **marine** | **https://doi.org/10.1099/ijs.0.02252-0** | **655** | **F** |
| **A0A1H7T5L9** | **Colwellia chukchiensis** | **Gammaproteobacteria** | **I** | **M** | **ocean** | **https://doi.org/10.1099/ijs.0.022111-0** | **650** | **F** |
| A0A1Y5FWE1 | Gammaproteobacteria bacterium 53_120_T64 | Gammaproteobacteria | M | M | marine oil plume simulation | https://doi.org/10.1073/pnas.1703424114 | 652 | F |
| A0A1Y5G2Z3 | Gammaproteobacteria bacterium 50_400_T64 | Gammaproteobacteria | M | M | marine oil plume simulation | https://doi.org/10.1073/pnas.1703424114 | 657 | F |
| A0A250KWS2 | Methylocaldum marinum | Gammaproteobacteria | I | M | marine sediment | https://doi.org/10.1099/ijs.0.063503-0 | 660 | R |
| A0A2A4LP23 | Cellvibrionales bacterium | Gammaproteobacteria | M | M | subseafloor aquifer | https://dx.doi.org/10.1038/ismej.2017.187 | 655 | no |
| A0A2A4VJE0 | Gammaproteobacteria bacterium | Gammaproteobacteria | M | M | subseafloor aquifer | https://dx.doi.org/10.1038/ismej.2017.187 | 651 | F |
| A0A2A5C283 | Cellvibrionales bacterium | Gammaproteobacteria | M | M | subseafloor aquifer | https://dx.doi.org/10.1038/ismej.2017.187 | 659 | F |
| A0A2D6RFY8 | Colwelliaceae bacterium | Gammaproteobacteria | M | M | ocean | https://dx.doi.org/10.1038/sdata.2017.203 | 651 | F |
| A0A2E0WMW7 | Gammaproteobacteria bacterium | Gammaproteobacteria | M | M | ocean | https://dx.doi.org/10.1038/sdata.2017.203 | 655 | R |
| A0A2E0YTU5 | Halioglobus sp. | Gammaproteobacteria | M | M | ocean | https://dx.doi.org/10.1038/sdata.2017.203 | 658 | F |
| A0A2G2IE75 | Colwellia sp. | Gammaproteobacteria | M | M | subseafloor aquifer | https://dx.doi.org/10.1038/ismej.2017.187 | 651 | F |

| | | | | | | | | |
|---|---|---|---|---|---|---|---|---|
| A0A2N1YEP3 | Gammaproteobacteria bacterium HGW-Gammaproteobacteria-15 | Gammaproteobacteria | M | T | terrestial sediment | https://dx.doi.org/10.1038/ismej.2017.39 | 655 | F |
| A0A317MTM7 | Plasticicumulans acidivorans | Gammaproteobacteria | I | W | wastewater | 10.1099/ijs.0.021410-0 | 663 | no |
| A0A317U3J5 | Legionella sp. Km488 (Legionella qingyii HEB18) | Gammaproteobacteria | I | F | Majiagou River | https://doi.org/10.1099/ijsem.0.003421 | 668 | no but NRPS |
| A0A349LX33 | Rheinheimera sp. | Gammaproteobacteria | M | ? | ? | http://dx.doi.org/.1038/nbt.4229 | 652 | F |
| A0A349Z0R5 | Alteromonas sp. | Gammaproteobacteria | M | M | ocean | http://dx.doi.org/.1038/nbt.4229 | 674 | F |
| A0A377GKY5 | Fluoribacter gormanii (Legionella gormanii) ATCC 33297 | Gammaproteobacteria | I | F | riverbank | J Clin Microbiol. 1980 Nov; 12(5): 718–721. | 652 | F |
| A0A3D0SPL4 | Gammaproteobacteria bacterium | Gammaproteobacteria | M | ? | ? | http://dx.doi.org/.1038/nbt.4229 | 662 | no |
| A0A3N1Y7J0 | Inmirania thermothiophila DSM 100275 | Gammaproteobacteria | I | HT | marine hydrothermal vent | https://doi.org/10.1099/ijsem.0.000767 | 657 | no |
| A0A426QM41 | Thiohalobacter thiocyanaticus | Gammaproteobacteria | I | HS | Hypersaline lake | https://dx.doi.org/10.1099/ijs.0.012880-0 | 656 | R |
| A0A431HTX1 | Xanthomonadales bacterium | Gammaproteobacteria | M | W | wastewater | https://doi.org/10.3389/fmicb.2019.00993 | 677 | no, MoaE-like |
| A0A432UXG8 | Thiothrix sp. | Gammaproteobacteria | M | HT | marine hydrothermal vent | https://doi.org/10.1038/s41396-019-0431-y | 655 | downstream F, edge of contig upstream |
| A0A486XSU6 | Rheinheimera sp. BAL341 | Gammaproteobacteria | ? | ? | ? | | 654 | F |
| A0A4E0QVS1 | Candidatus Thiomargarita nelsonii | Gammaproteobacteria | S | M | methane seep | https://dx.doi.org/10.3389/fmicb.2016.00603 | 658 | downstream NRPS |
| A0A4P7BWL2 | Nitrosococcus wardiae | Gammaproteobacteria | I | M | marine | https://dx.doi.org/10.3389%2Ffmicb.2016.00512 | 669 | R |
| A0A4R2LU62 | Plasticicumulans lactativorans | Gammaproteobacteria | I | W | wastewater | 10.1099/ijs.0.051045-0 | 664 | no, MoaE-like |
| A0A4V6PYV6 | Methylocaldum sp. 0917 | Gammaproteobacteria | I | T | desert soil | | 660 | R |
| A4BSM0 | Nitrococcus mobilis Nb-231 (ATCC 25380) | Gammaproteobacteria | | M | marine | https://www.atcc.org/products/all/25380.aspx | 672 | R 4 genes back |
| D5C057 | Nitrosococcus halophilus (strain Nc4) | Gammaproteobacteria | I | M | marine | https://doi.org/10.1111/j.1574-6941.2010.01027.x | 669 | R |
| Q3J9P0 | Nitrosococcus oceani (strain ATCC 19707 / BCRC 17464 / NCIMB 11848 / C-107) | Gammaproteobacteria | I | M | ocean | https://doi.org/10.1111/j.1574-6941.2010.01027.x | 672 | R |
| **U1KR87** | **Pseudoalteromonas rubra DSM 6842** | **Gammaproteobacteria** | **I** | **M** | **marine** | **INTERNATIONAL JOURNAL OF SYSTEMATIC BACTERIOLOGY, Oct. 1976, p. 459-466** | **664** | **F** |
| W6LT00 | Candidatus Contendobacter odensis Run_B_J11 | Gammaproteobacteria | M | W | wastewater | http://dx.doi.org/10.1038/ismej.2013.162 | 658 | unclear, on edge of contig |
| W8L8B1 | Ectothiorhodospira haloalkaliphila | Gammaproteobacteria | I | HS | Hypersaline lake | http://dx.doi.org/10.7150/jgen.9123 | 660 | F |
| A0A1J5Q684 | mine drainage metagenome | Unassigned metagenomic contigs | C | HA | acid mine drainage | https://www.ebi.ac.uk/ena/data/view/PRJNA343431 | 670 | no but methyltransferase |
| A0A3B0RW05 | hydrothermal vent metagenome | Unassigned metagenomic contigs | C | HT | hydrothermal vent | | 659 | F 2 genes back |

### 3 Methods

#### 3.1 Computational methods

Jupyter notebooks for CO-ED analysis are publicly available at https://github.com/tderond/CO-ED. Besides Python code, the Jupyter notebooks contain explanations of the workflow's steps and example command-line instructions. Here, we will briefly explain the steps of the CO-ED workflow, but for more details, please consult the Jupyter notebooks.

Briefly, the CO-ED workflow has the following stages:

- Annotate domains (See section 3.1.1) for a set of "query" protein sequences, and a set of "known enzymes" (See section 3.1.2).
- For every domain in the curated set of non-redundant enzymatic/catalytic domains (See section 3.1.3), find its occurrences in each of the proteins in the query set.
- Draw a node for each domain found in at least one protein in the query set, and label the number of proteins the domain was found in.
- For every **combination of two domains** in the curated set of non-redundant enzymatic/catalytic domains (See section 3.1.3), find the occurrences of each combination in the query set and in the "known enzymes" sets.
- Draw an edge for each combination of domains found in at least one protein in the query set, label it with the number of occurrences, and color the edge based on which set(s) of "known enzymes" contain at least one entry with this combination of domains.

##### 3.1.1 Domain annotation

Pfam-A domain annotations were either:

- Retrieved from Uniprot: this was done for the "Known" enzymes and for the all-of-uniprot analysis, or,
- Annotated using the PfamScan script (http://ftp.ebi.ac.uk/pub/databases/Pfam/Tools/): This was done for the single-genome queries. Annotating the domains this way gives the CO-ED workflow more flexibility, such as allowing for the analysis of genomes that are not in Uniprot, and the detection of domains that are not included in Pfam-A

##### 3.1.2 Selection of "known" enzymes

- Uniprot: All proteins in UniprotKB that **either** are manually annotated as having catalytic activity (i.e. match the query "annotation:(type:"catalytic activity" evidence:manual)") **or** are listed in Uniprot's pathway.txt file (https://www.uniprot.org/docs/pathway). This set is overlaps with, but is **not identical** to "Swissprot" (the subset of Uniprot marked "reviewed"): It contains some "unreviewed" entries, and Swissprot entries whose catalytic activity was automatically assigned by homology are not included.
- BRENDA: All entries in the BRENDA database that refer to Uniprot accessions
- MIBiG: For each entry in MIBiG, an NCBI nucleotide region is defined. NCBI protein identifiers for all proteins in this region were obtained and mapped to Uniprot accessions. A small number of proteins that could not be mapped to Uniprot were not included in the final set.

##### 3.1.3 Curation of enzyme Domains for CO-ED

The set of domains used to conduct CO-ED is intended to be enzymatic and non-redundant. With "enzymatic" we mean that only catalytic enzyme domains are included, and e.g. purely structural, regulatory, or docking domains are omitted. With "non-redundant" we mean that we wish to avoid "trivial" domain pairs where two crystallographic Pfam domains work together to catalyze a single reaction, such as "Terpene_synth" and "Terpene_synth_C", or "G6PD_N" and "G6PD_C". This section describes how this set of domains was curated.

A set of Pfam domains was compiled by taking all domains annotated for entries in Uniprot (www.uniprot.org) that are also annotated in MIBiG (mibig.secondarymetabolites.org, all proteins), BRENDA (www.brenda-enzymes.org, all proteins), or in Uniprot's pathway.txt. Non-catalytic domains were removed. For pseudo-catalytic domains that together catalyze one reaction (often detected by performing CO-ED analysis on all proteins in Uniprot and finding domains that co-occur with another a high percentage of the time), the more abundant domain was included. Overlapping domains with similar catalytic functions are often members of the same Pfam "Clan", causing only the best-matching domain to be annotated by PfamScan, but in some cases where both are annotated in a high proportion of proteins in Uniprot (e.g., because clan assignment has yet not been completed for those domains), only one of the proteins was included in our set.

Lastly, many enzymatic domains acting on macromolecules and domains with unknown functions were annotated as such, and the analysis can be run with or without their consideration. Annotation categories are as follows: "m": nucleases, topoisomerases, transposases, helicases, polymerases, proteases, protein kinases and phosphatases, ATP-dependent transporters; "s": glycosyltransferases, glycosylhydrolases (cellulases, amylases, etc.); "e": enzymes in electron transport chains (oxidative phosphorylation, photosynthesis, etc.); "u": domains with unknown function; "y": all other enzymes, but only those transporters that couple transport to a reaction besides ATP hydrolysis; "n": determined to either not be catalytic or to comprise a catalytic domain together with a domain annotated in one of the above categories. For the analyses shown in this manuscript, annotation categories "y", "m", "s" and "e" were considered, totaling 1745 domains. We realize the domain curation process is somewhat subjective, and hence the CO-ED analysis Jupyter notebook allows for facile re-analysis with different sets of domains.

We realize the domain curation process is somewhat subjective, and hence the CO-ED analysis Jupyter notebook allows for facile re-analysis with different sets of domains by editing the `pfamID_to_name_desc_longdesc.tsv` file.

#### 3.2 Instrumentation

##### 3.2.1 HPLC-UV-MS

Samples were analyzed on an Agilent 1290 infinity liquid chromatography system with a diode array detector and Agilent 6530 Q-TOF mass spectrometer with a Dual Spray Electrospray Ionization source in positive ionization mode. Chromatography conditions were as follows: Poroshell 120 Phenyl-Hexyl column (100 × 4.6 mm, 2.7 µm particle size). Mobile phase: A: 0.1 % formic acid in water, B: 0.1 % formic acid in acetonitrile. Gradient: 40% B for 2 minutes, ramp to 100% B in 10 minutes, 100% B for 4 minutes, ramp to 40% B in 0.5 minutes, 40% B for 4.5 minutes. Flow rate: 0.5 mL/min. Source parameters: Drying gas: 11 liters per minute, 300 °C; Nebulizer: 35 psig; Capillary: 3000 V; Fragmentor: 100 V; Skimmer: 65 V; OCT 1 RF Vpp: 750. Tandem MS (Collision Induced Dissociation) parameters: Isolation width: Narrow; Collision energy: 20 V.

##### 3.2.2 NMR

Spectra were recorded on a JEOL spectrometer (500 MHz) or a Bruker Avance III spectrometer (600 MHz) with a 1.7 mm inverse-detection triple-resonance (H-C/N/D) cryoprobe. Chemical shifts are referenced to the residual solvent signal.

#### 3.3 Bacteriological culture

Bacterial stocks were obtained from DSMZ (Deutsche Sammlung von Mikroorganismen und Zellkulturen GmbH), and cultured at 30 °C in media indicated in the table below.

| **Species Name** | **DSM Number** | **Media Type** |
|---|---|---|
| *Pseudoalteromonas rubra* | DSM 6842 | Marine agar/broth 2216 |
| *Colwellia chukchiensis* | DSM 22576 | Marine agar/broth 2216 |
| *Rheinheimera pacifica* | DSM 17616 | Marine agar/broth 2216 |
| *Undibacterium pigrum* | DSM 19792 | Reasoner's 2A (R2A) agar/broth |
| *Skermanella aerolata* | DSM 18479 | Reasoner's 2A (R2A) agar/broth |

#### 3.4 Molecular cloning

Genomic DNA was isolated from overnight liquid cultures using the Zymo Research Quick-DNA Miniprep Kit. *oxz* genes were PCR amplified from this genomic DNA using NEB Q5 High-Fidelity polymerase. Cycling conditions were as follows: initial denaturation temperature of 98 °C for 30 seconds followed by 35 cycles of a denaturing temperature of 98 °C for 10 seconds, an annealing temperature of 68 °C for 30 seconds, and a 72 °C extension for 5 minutes. After cycling, the temperature was set to 72 °C for 2 minutes before going down to 10 °C. All amplicons were purified by gel extraction. *oxzAB* gene pairs were cloned using the Thermo Fischer ZERO Blunt TOPO kit, and Pr*oxzA* and Pr*oxzB* were cloned into pET28a without and with MBP respectively using the NEB Hi-Fi Assembly master mix. All plasmids were verified by Sanger sequencing.

All plasmid and primer sequences used in this study are available at the following URL: https://benchling.com/tderond/f_/TUemteIN-de-rond-et-al-2020/

#### 3.5 Metabolite analysis of heterologously expressed *oxzAB*

pTOPO-*oxzAB* plasmids were transformed into *E. coli* BLR(DE3) and grown up in 50 mL of terrific broth (TB), 10 mL/L glycerol, and 50ug/mL Kan at 30 °C, induced with 0.1 mM IPTG at an OD of 0.8, and left to shake overnight at 30 °C. The cells were pelleted at 10,000 × g, and the pellets were frozen, lyophilized, extracted for two hours in ethyl acetate. The extracts were filtered through glass pipette cotton filters, evaporated under a stream of nitrogen gas, redissolved in 200 µL 35:65 acetonitrile:water, filtered through a 0.2 µm filter, and analyzed by HPLC-UV-MS as described in section 3.2.1

#### 3.6 Bulk heterologous production and isolation of oxazolones

*E. coli* BLR(DE3) containing pTOPO-Pr_oxzAB or pTOPO-Sa_oxzAB was grown up in 3x 1.5L Terrific Broth with 50 µg/mL Kan and 10 mL/L of glycerol for 3 days at 30 °C. The cultures were centrifuged at 10,000g for 20 minutes, and the pellets were frozen and lyophilized for two days. The dried biomass was crushed, extracted overnight in 100 mL ethyl acetate, filtered, dried under reduced pressure. The residue was subjected to flash chromatography using hexanes:ethyl acetate on a CombiFlash EZPrep system using a 24g ReadySep gold silica gel column. UV active peaks were then further subjected to preparative reverse phase HPLC (Phenomenex Luna c18 column (100mm x 21.2 mm, 5 µm particle size) using isocratic conditions between 80:20 and 90:10 acetonitrile:water with 0.1% formic acid, and evaporated by rotory evaporation followed by lyophilization, yielding between 1mg and 9mg of purified oxazolone.

##### 3.6.1 Nonyltyrazolone (Tyrz, R = C9)

IUPAC name: (Z)-4-(4-hydroxybenzylidene)-2-nonyloxazol-5-one

Yellow powder

$^1$H NMR (500 MHz, Chloroform-*d*) δ 8.03 (d, *J* = 8.7 Hz, 2H), 7.10 (s, 1H), 6.90 (d, *J* = 8.7 Hz, 2H), 5.56 (s, 1H), 2.64 (t, *J* = 7.6 Hz, 2H), 1.79 (p, *J* = 7.6 Hz, 2H), 1.46 – 1.38 (m, 2H), 1.38 – 1.22 (m, 12H), 0.88 (t, *J* = 6.8 Hz, 3H).

$^{13}$C NMR (126 MHz, CHLOROFORM-*D*) δ 168.58, 168.36, 158.51, 134.71, 131.55, 130.61, 126.52, 116.16, 31.98, 29.57, 29.51, 29.38, 29.32, 29.25, 25.48, 22.80, 14.25.

HRMS(ESI): observed 316.1905, expected 316.1907 for $[M+H]^+$

##### 3.6.2 Decyltyrazolone (Tyrz, R = C10)

IUPAC name: (Z)-4-(4-hydroxybenzylidene)-2-decyloxazol-5-one

Yellow powder

$^1$H NMR (500 MHz, Chloroform-*d*) δ 8.04 (d, *J* = 8.7 Hz, 2H), 7.09 (s, 1H), 6.90 (d, *J* = 8.5 Hz, 2H), 2.64 (t, *J* = 7.6 Hz, 2H), 1.79 (p, *J* = 7.7 Hz, 2H), 1.42 (p, *J* = 7.2 Hz, 2H), 1.38 – 1.20 (m, 14H), 0.88 (t, *J* = 6.8 Hz, 3H).

HRMS(ESI): observed 330.207, expected 330.206 for $[M+H]^+$

##### 3.6.3 Undecyltyrazolone (Tyrz, R = C11)

IUPAC name: (Z)-4-(4-hydroxybenzylidene)-2-undecyloxazol-5-one

Yellow powder

$^1$H NMR (599 MHz, DMSO-$d_6$) δ 8.03 (d, *J* = 8.4 Hz, 2H), 7.08 (s, 1H), 6.85 (d, *J* = 8.3 Hz, 2H), 2.57 (t, *J* = 7.7 Hz, 2H), 1.64 (p, *J* = 7.6 Hz, 2H), 1.30 (q, *J* = 7.7 Hz, 2H), 1.18 (d, *J* = 16.2 Hz, 14H), 0.80 (t, *J* = 7.0 Hz, 3H).

$^1$H NMR (500 MHz, Chloroform-*d*) δ 8.03 (d, *J* = 8.5 Hz, 2H), 7.09 (s, 1H), 6.90 (d, *J* = 8.5 Hz, 2H), 5.26 (s, 1H), 2.64 (t, *J* = 7.6 Hz, 2H), 1.79 (p, *J* = 7.6 Hz, 2H), 1.42 (p, *J* = 7.0 Hz, 2H), 1.27 (d, *J* = 10.6 Hz, 14H), 0.88 (t, *J* = 6.7 Hz, 3H).

$^{13}$C NMR (151 MHz, DMSO) δ 167.71, 167.25, 161.20, 134.69, 131.27, 128.97, 124.37, 116.13, 31.52, 29.24, 29.10, 28.97, 28.86, 28.68, 28.62, 24.82, 22.31, 14.01.

HRMS(ESI): observed 344.2217, expected 344.2220 for $[M+H]^+$

##### 3.6.4 ω-6-undecenyltyrazolone (Tyrz, R = C11:1 ω-6)

IUPAC name: 4-((Z)-4-hydroxybenzylidene)-2-((Z)-undec-4-en-1-yl)oxazol-5-one

Yellow powder

$^1$H NMR (500 MHz, Chloroform-*d*) δ 8.04 (d, *J* = 8.8 Hz, 2H), 7.09 (s, 1H), 6.89 (d, *J* = 8.6 Hz, 2H), 5.45 (dt, *J* = 10.7, 7.2 Hz, 1H), 5.36 (dt, *J* = 11.0, 7.2 Hz, 1H), 2.65 (t, *J* = 7.5 Hz, 2H), 2.19 (q, *J* = 7.3 Hz, 2H), 2.03 (q, *J* = 7.1 Hz, 2H), 1.86 (p, *J* = 7.4 Hz, 2H), 1.38 – 1.20 (m, 8H), 0.87 (t, *J* = 6.7 Hz, 4H).

HRMS(ESI): observed 342.207 , expected 342.206 for $[M+H]^+$

##### 3.6.5 Heptylphenazolone (Phez, R = C7)

IUPAC name: (Z)-4-benzylidene-2-heptyloxazol-5-one

White powder

$^1$H NMR (500 MHz, Chloroform-*d*) δ 8.11 – 8.07 (m, 2H), 7.48 – 7.41 (m, 3H), 7.14 (s, 1H), 2.66 (t, *J* = 7.6 Hz, 2H), 1.80 (p, *J* = 7.6 Hz, 2H), 1.43 (td, *J* = 9.0, 8.4, 4.9 Hz, 2H), 1.39 – 1.26 (m, 6H), 0.89 (t, *J* = 6.7 Hz, 3H).

$^{13}$C NMR (126 MHz, CHLOROFORM-*D*) δ 169.45, 168.15, 133.38, 132.76, 132.36, 131.52, 131.21, 129.01, 31.73, 29.60, 29.19, 28.97, 25.39, 22.72, 14.20.

HRMS(ESI): observed 272.1647, expected 272.1645 for $[M+H]^+$

#### 3.7 Methanol adduct formation

2 mg Nonyltyrazolone or Heptylphenazolone was dissolved in 1 mL of methanol, 10 mg of $K_2CO_3$ was added, upon which the Nonyltyrazolone reaction turned bright yellow (See image in section 1.6c). The reactions were left to react for 5 minutes at room temperature, filtered, and purified by preparative reverse phase HPLC as described in section 3.6 under isocratic conditions of 78:22 acetonitrile:water with 0.1% formic acid.

##### 3.7.1 *N*-decanoyldehydrotyrosine methyl ester

IUPAC name: methyl (Z)-2-decanamido-3-(4-hydroxyphenyl)acrylate

White powder

$^1$H NMR (500 MHz, Methanol-*d*$_4$) δ 7.49 (d, *J* = 8.7 Hz, 2H), 7.41 (s, 1H), 6.79 (d, *J* = 8.7 Hz, 2H), 3.78 (s, 3H), 2.38 (t, *J* = 7.4 Hz, 2H), 1.70 (p, *J* = 7.4 Hz, 2H), 1.44 – 1.25 (m, 12H), 0.94 – 0.88 (m, 3H).

$^1$H NMR (599 MHz, DMSO-*d*$_6$) δ 9.39 (s, 1H), 7.48 (d, *J* = 8.3 Hz, 2H), 7.15 (s, 1H), 6.75 (d, *J* = 8.3 Hz, 2H), 3.65 (s, 3H), 2.25 (t, *J* = 7.2 Hz, 2H), 1.55 (p, *J* = 7.1 Hz, 2H), 1.26 (t, *J* = 10.1 Hz, 12H), 0.85 (t, *J* = 6.6 Hz, 3H).

$^{13}$C NMR (126 MHz, METHANOL-*D*$_4$) δ 176.38, 167.63, 161.00, 136.55, 133.24, 125.88, 123.39, 116.68, 52.75, 36.82, 33.06, 30.67, 30.54, 30.45, 30.36, 26.73, 23.75, 14.44.

HRMS(ESI): observed 348.2174, expected 348.2169 for $[M+H]^+$

##### 3.7.2 *N*-octanoyldehydrophenylalanine methyl ester

IUPAC name: methyl (Z)-2-octanamido-3-phenylacrylate

White powder

$^1$H NMR (500 MHz, DMSO-*d*$_6$) δ 9.60 (s, 1H), 7.62 (d, *J* = 7.2 Hz, 2H), 7.44 – 7.34 (m, 3H), 7.18 (s, 1H), 3.69 (s, 3H), 2.26 (t, *J* = 7.2 Hz, 2H), 1.55 (p, *J* = 7.2 Hz, 2H), 1.34 – 1.18 (m, 6H), 0.87 (t, *J* = 6.9 Hz, 3H).

$^{13}$C NMR (126 MHz, DMSO-*D*$_6$) δ 172.52, 165.67, 133.49, 131.16, 129.86, 129.40, 128.63, 126.78, 52.18, 34.98, 31.28, 28.56, 28.50, 25.01, 22.12, 14.03.

#### 3.8 Chemical synthesis of OxzB substrate

10 mmol (1 equivalent) of decanoyl chloride and 4 equivalents of 4N NaOH were added to 10mmol Tyrosine. An additional 4 equivalents of decanoyl chloride was added over the course of 2 hours while stirring at room temperature.

Four more equivalents of NaOH and 10mL of methanol were added and allowed to stir at room temperature for half an hour. The reaction mixture was evaporated by rotory evaporation followed by lyophilization. The dried reaction mixture was chromatographed over silica gel to afford *N*-decanoyltyrosine

##### 3.8.1 *N*-decanoyl-L-tyrosine

White powder

$^{1}$H NMR (500 MHz, DMSO-$d_6$) δ 8.01 (d, *J* = 8.2 Hz, 1H), 6.99 (d, *J* = 8.4 Hz, 2H), 6.99 (d, *J* = 8.4 Hz, 2H), 6.63 (d, *J* = 8.5 Hz, 2H), 4.32 (ddd, *J* = 9.7, 8.1, 4.8 Hz, 1H), 3.36 (bs, 1H), 2.91 (dd, *J* = 13.8, 4.8 Hz, 1H), 2.71 (dd, *J* = 13.9, 9.6 Hz, 1H), 2.03 (t, *J* = 7.3 Hz, 2H), 1.39 (p, *J* = 7.5 Hz, 2H), 1.31 – 1.08 (m, 12H), 0.85 (t, *J* = 6.9 Hz, 3H).

$^{13}$C NMR (126 MHz, DMSO-$D_6$) δ 173.44, 172.19, 155.90, 129.99, 127.79, 114.91, 53.70, 36.05, 35.10, 31.34, 28.94, 28.86, 28.75, 28.54, 25.24, 22.16, 14.01.

#### 3.9 Protein purification and enzyme assays

Overnight cultures of *E. coli* BLR(DE3) harboring pET28a-His6-PrOxzA or pET28a-MBP-PrOxzB respectively were diluted into 1.5 L Terrific Broth with 50 µg/mL Kan and 10 mL/L of glycerol at 30 °C. When an OD of 0.8 was reached, the flasks were cooled to 18 °C and induced with 50 µM IPTG and left to shake at 18 °C overnight. The cells were pelleted at 4 °C for 15-20 minutes at 15,000 × g and resuspended in 40mL cold 50mM Tris, 200 mM NaC,l and 10% glycerol at pH 7 ("lysis buffer"), and sonicated at 50% amplitude, 15 seconds on, 15 seconds off for 5 minutes. The proteins were purified in "batch" format using loose resin in 50 mL falcon tubes. For His6-PrOxzA, 2.5 mL Nickel-IDA resin was used, the resin washed twice with 40mL lysis buffer + 50 mM imidazole, eluted with 5 mL lysis buffer + 500 mM imidazole, and concentrated using a 15 kDa MWCO filter. For MBP-PrOxzB, 2.5 mL NEB amylose resin was used, the resin was washed twice with 40 mL lysis buffer, eluted with 5mL, and concentrated using a 50 kDa MWCO filter. MBP-PrOxzB is visibly yellow, suggesting it binds a flavin cofactor, as expected from nitroreductase-family enzymes.

In vitro assays of His6-OxzA and MBP-OxzB were conducted in 200uL 50 mM potassium phosphate pH 7, 200 mM NaCl, 10% DMSO, with reactants at the following final concentrations: 150 µM ATP, and 150 µM decanoyl-CoA, 100 µM L-Tyrosine, and 100 µM N-acyltyrosine. After 1 hour, the reactions were quenched with 100 uL acetonitrile, filtered through 0.2 µM filters, and analyzed as described in section 3.2.1

#### 3.10 Analysis of oxazolone production in *P. rubra* and *C. chukchiensis*

Cells were grown as a lawn on Marine Agar 2216 with or without a drop of antibiotic stock (see table below) in the middle of the plate. After 2 days of growth, biomass adjacent to the zone of inhibition was harvested by scraping the plate. An effort was made to harvest a roughly equivalent amount of biomass from each plate. The biomass was lyophilized, extracted with ethyl acetate for 2 hours, filtered through glass pipette cotton filters, evaporated under a stream of nitrogen gas, redissolved in 200 uL 35:65 acetonitrile:water, filtered through a 0.2 µm filter, and analyzed by HPLC-UV-MS as described in section 3.2.1.

| Antibiotic | Stock Concentration (mg/mL) | Droplet volume (µL) |
|---|---|---|
| Bacitracin | 50 | 10 |
| Chloramphenicol | 30 | 10 |
| Erythromycin | 5 | 30 |
| Fosfomycin | 25 | 30 |
| Glyphosate | 100 | 10 |
| Kanamycin | 50 | 10 |
| Nalidixic Acid | 25 | 20 |
| Rifampicin | 10 | 10 |
| Tetracycline | 5 | 30 |

### 4 NMR Spectra

#### 4.1 $^1$HNMR

##### 4.1.1 Nonyltyrazolone

###### *4.1.1.1 Nonyltyrazolone 1HNMR CDCl3 500 MHz*

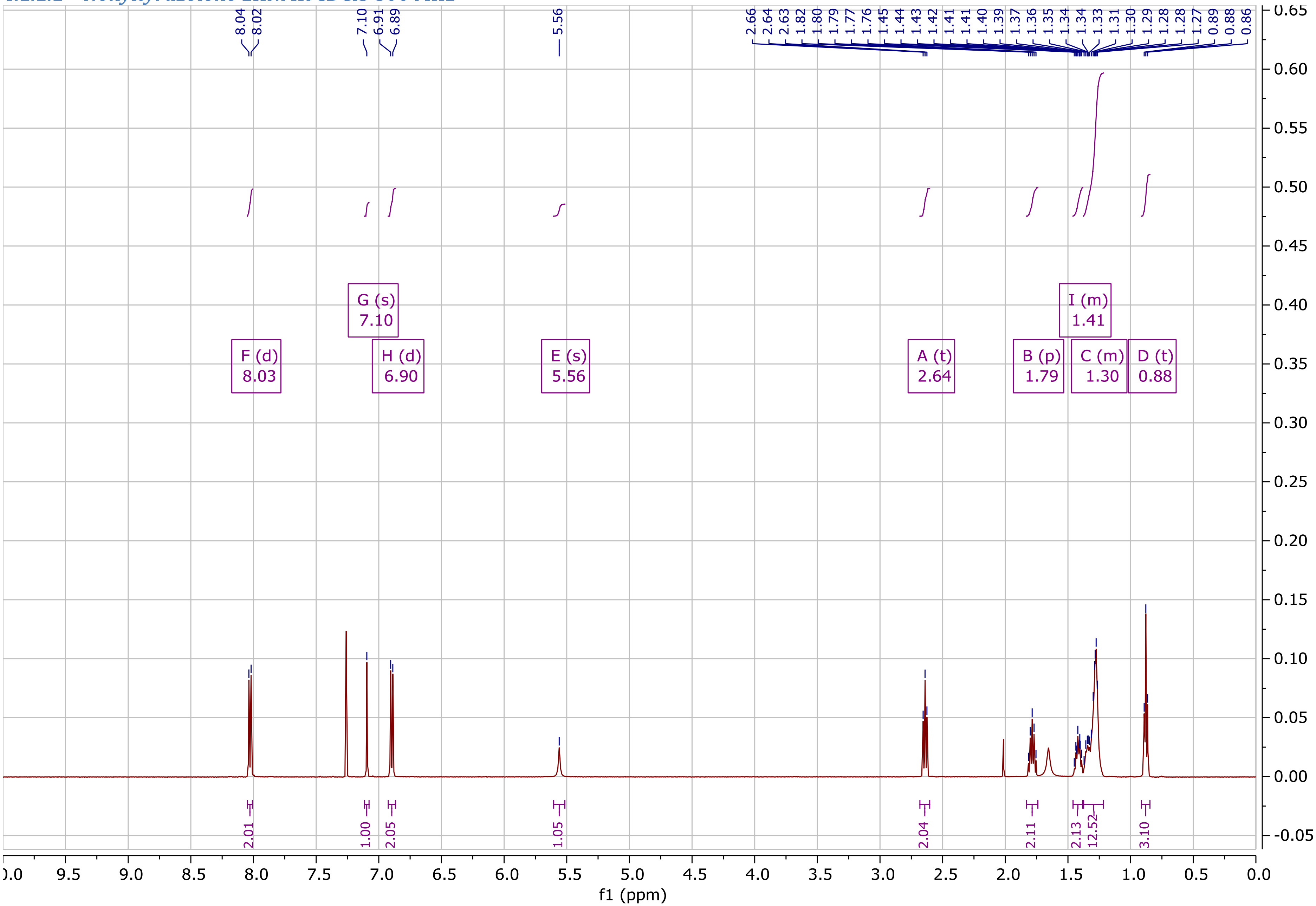

##### 4.1.2 Decyltyrazolone

###### *4.1.2.1 Decyltyrazolone 1HNMR CDCl3 500MHz*

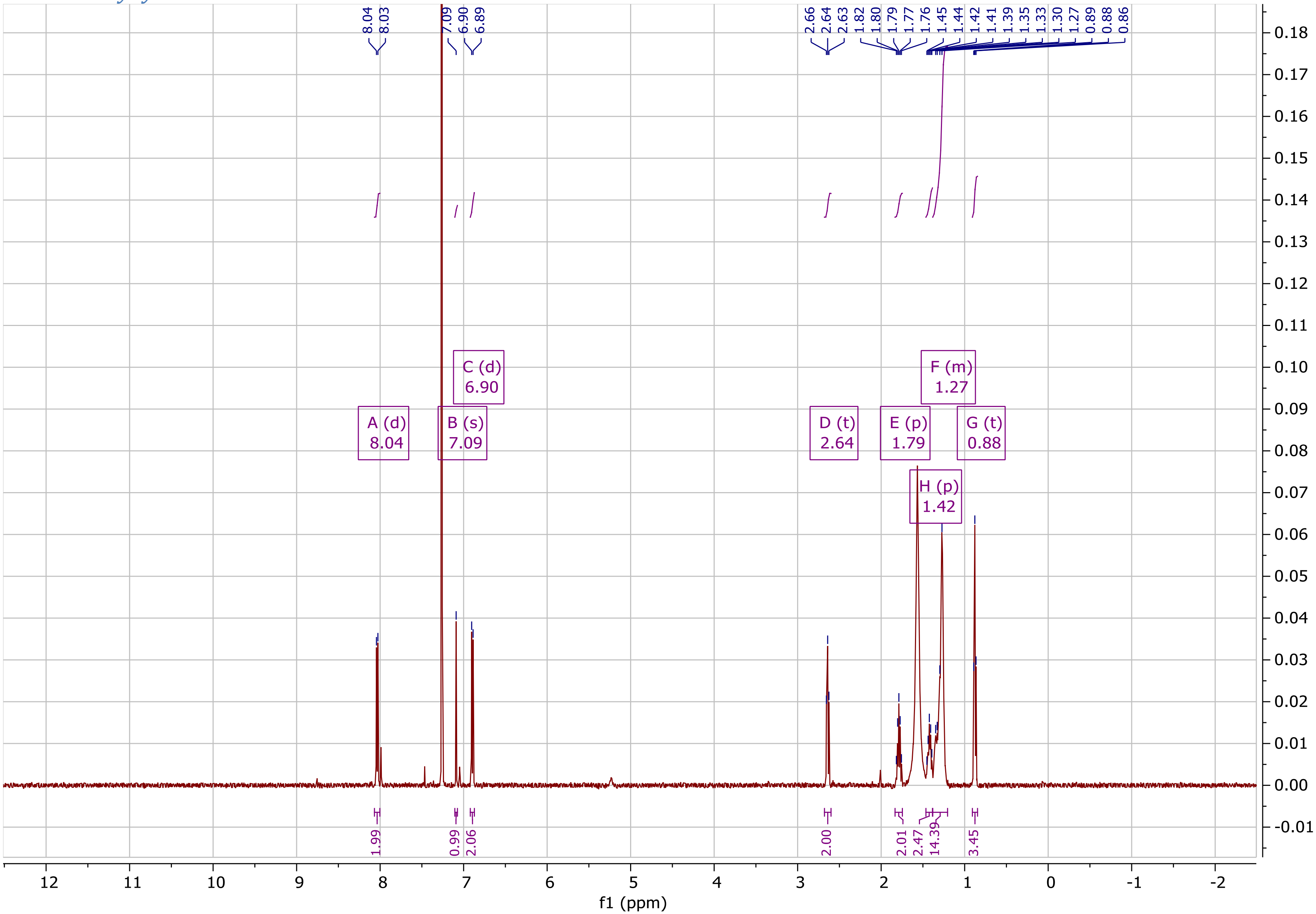

##### 4.1.3 Undecyltyrazolone

###### *4.1.3.1 Undecyltyrazolone 1HNMR DMSO-d6 600MHz*

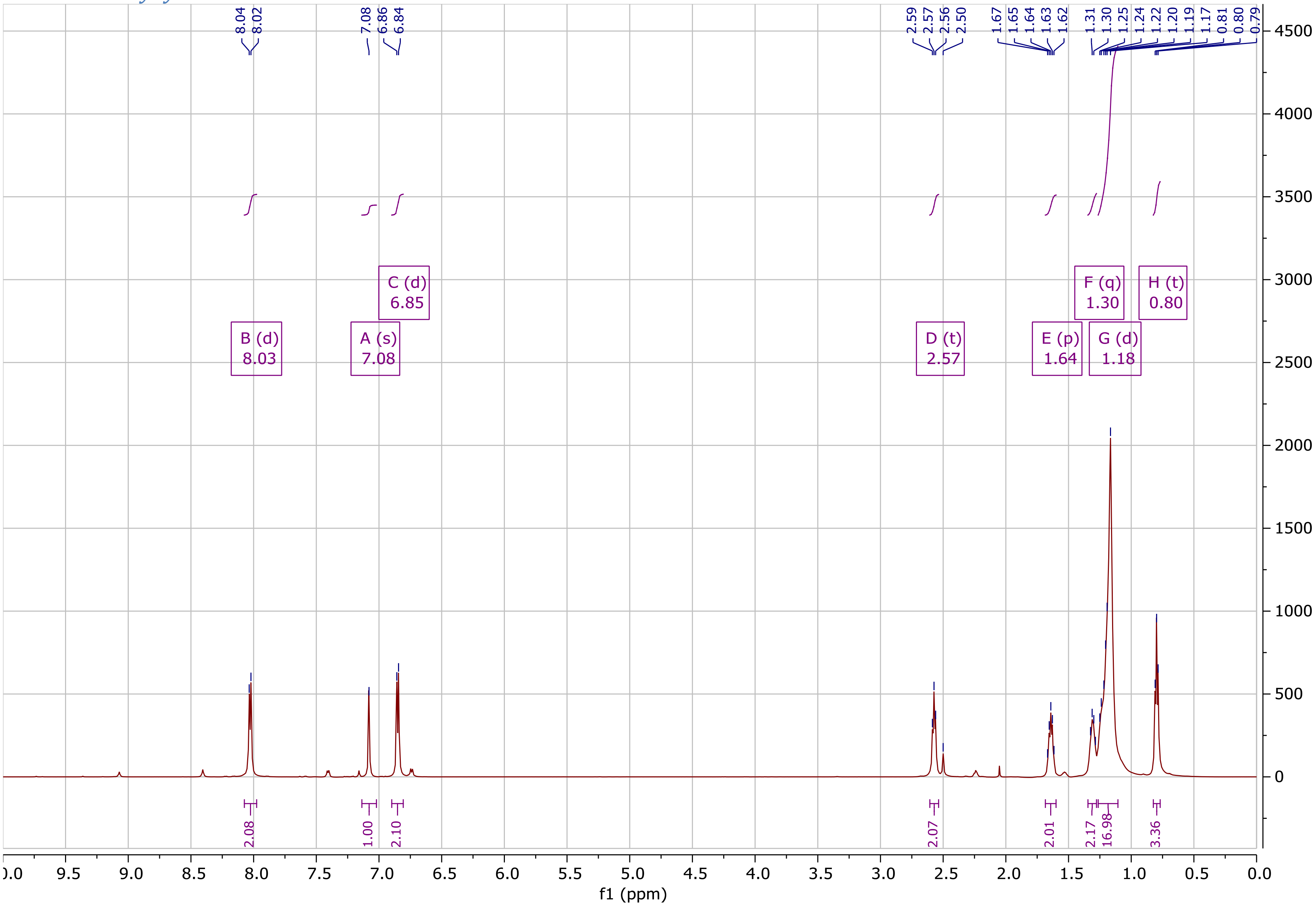

##### *4.1.3.2 Undecyltyrazolone 1HNMR CDCl3 500MHz*

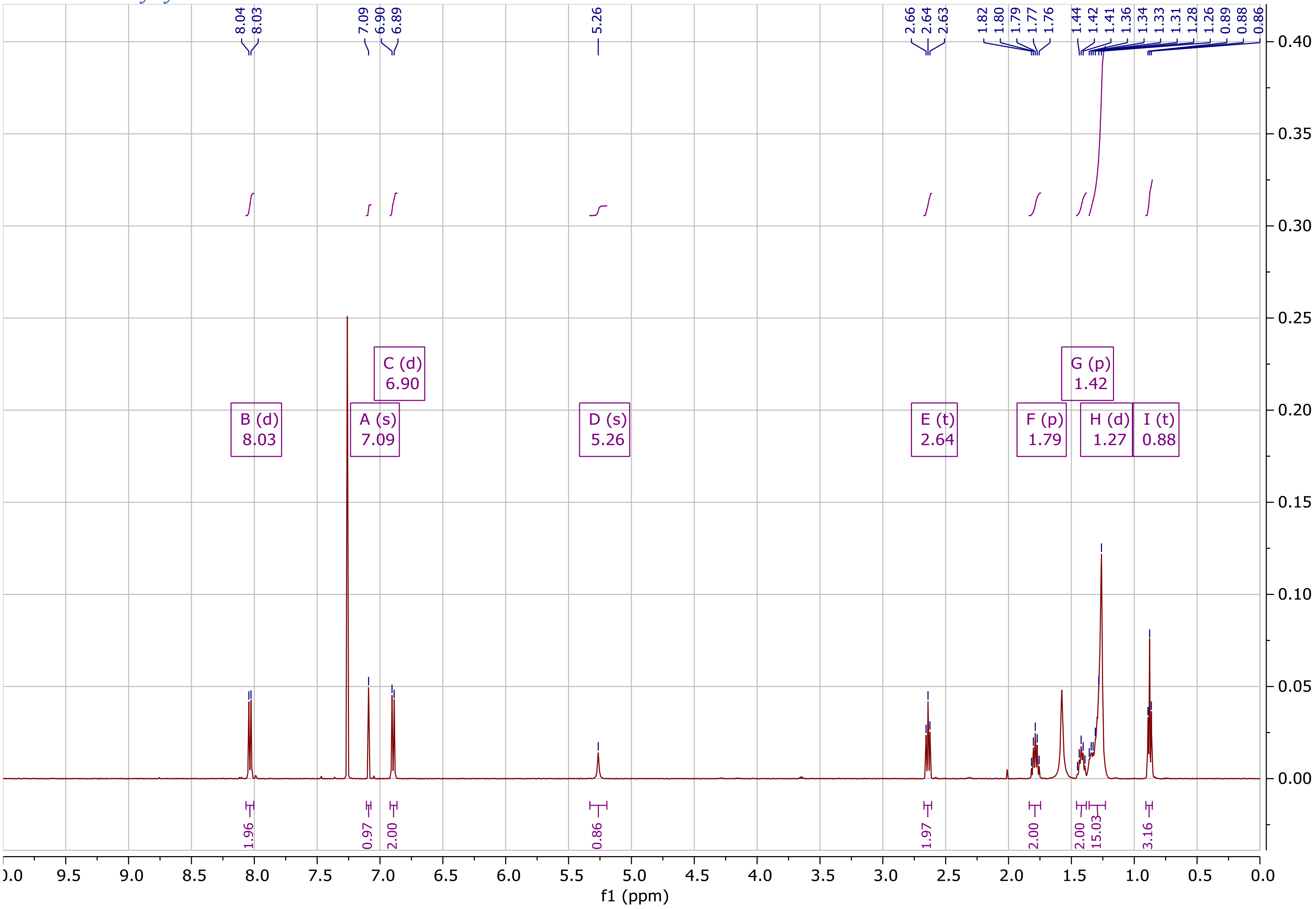

##### 4.1.4 ω-6-undecenyltyrazolone

###### *4.1.4.1 ω-6-undecenyltyrazolone 1HNMR CDCl3 500 MHz*

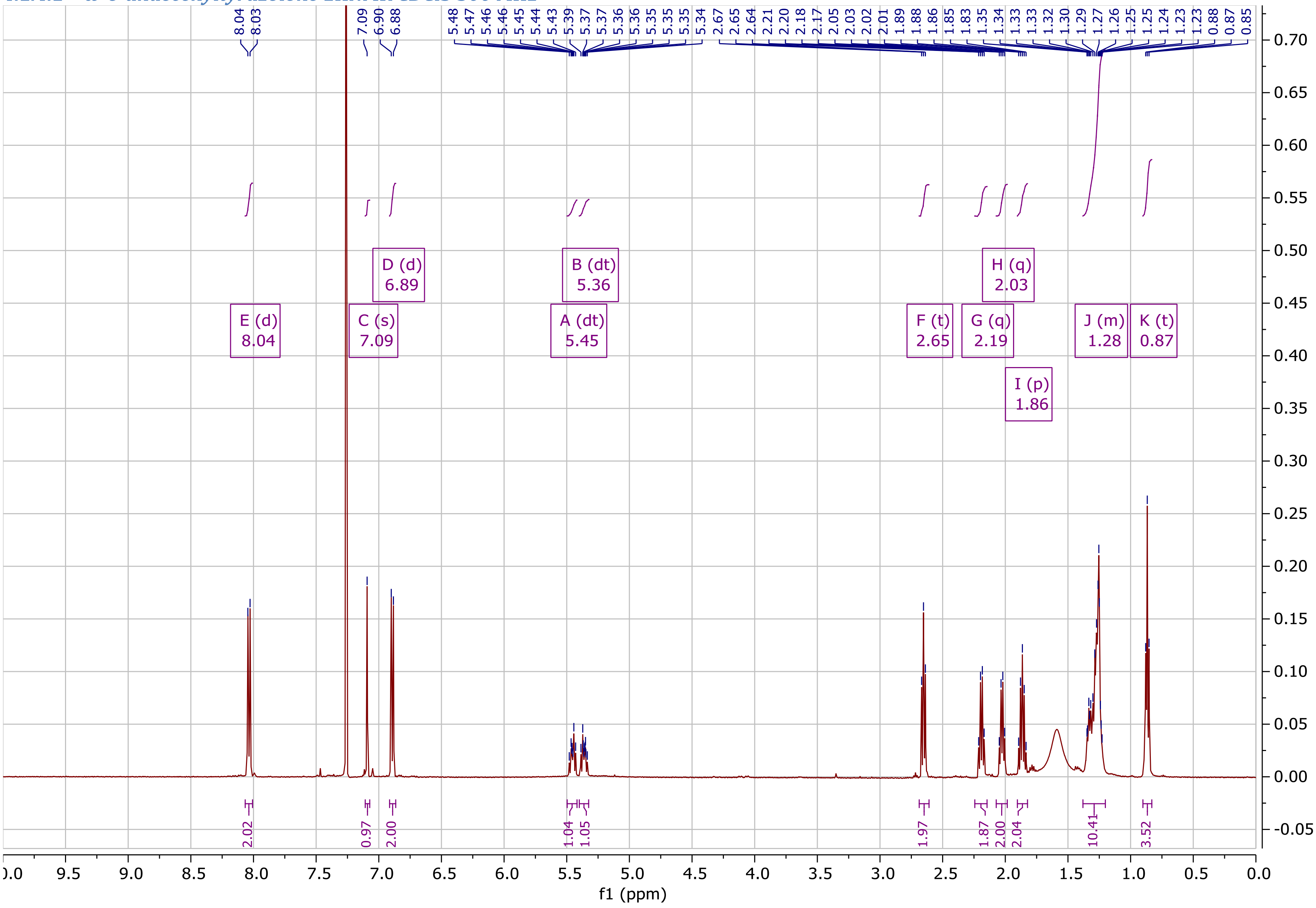

#### 4.1.5 Heptylphenazolone

##### *4.1.5.1 Heptylphenazolone 1HNMR CDCl3 500MHz*

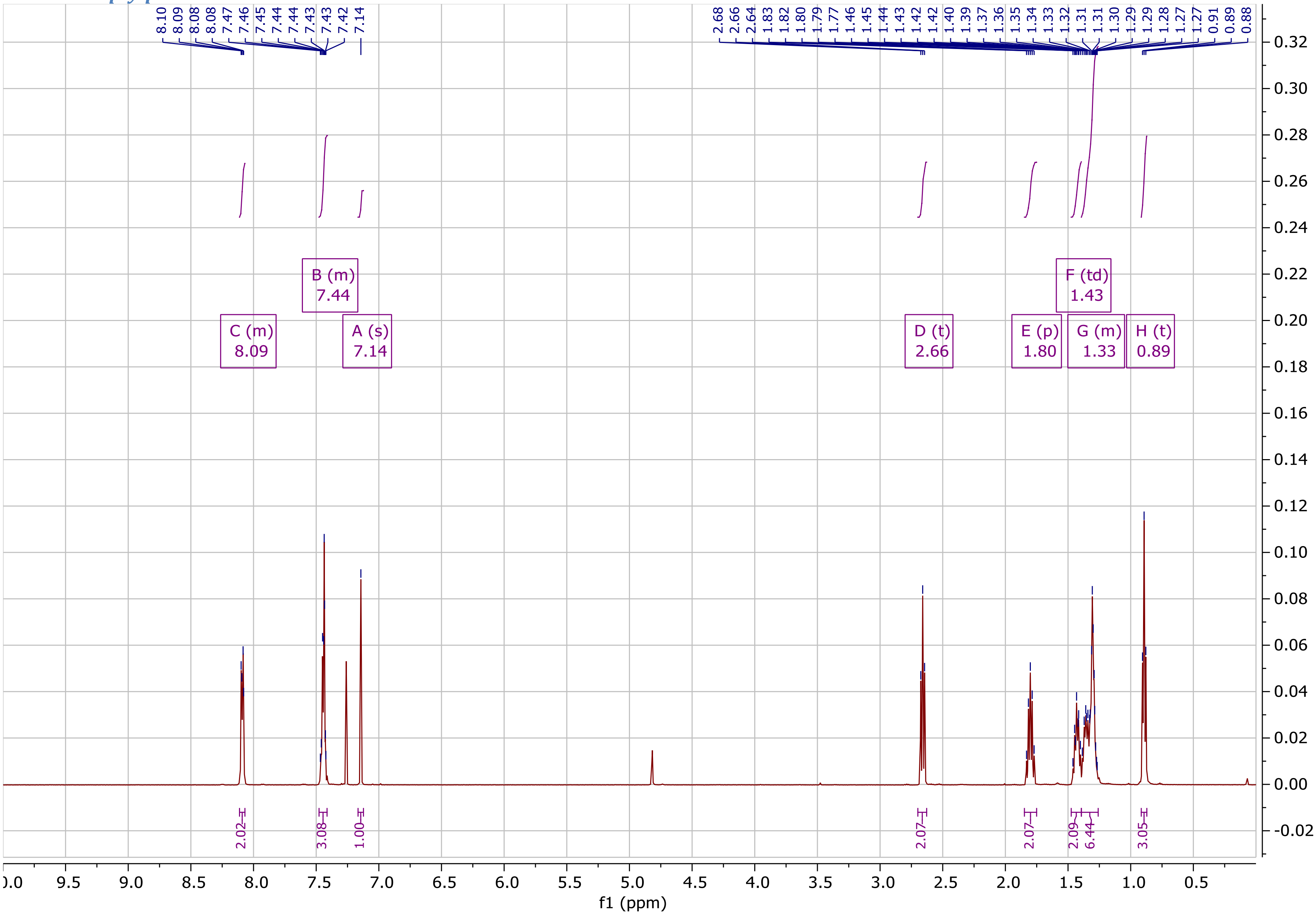

##### 4.1.6 N-decanoyldehydrotyrosine

###### *4.1.6.1 N-decanoyldehydrotyrosine methyl ester 1HNMR MeOH-d4 500MHz*

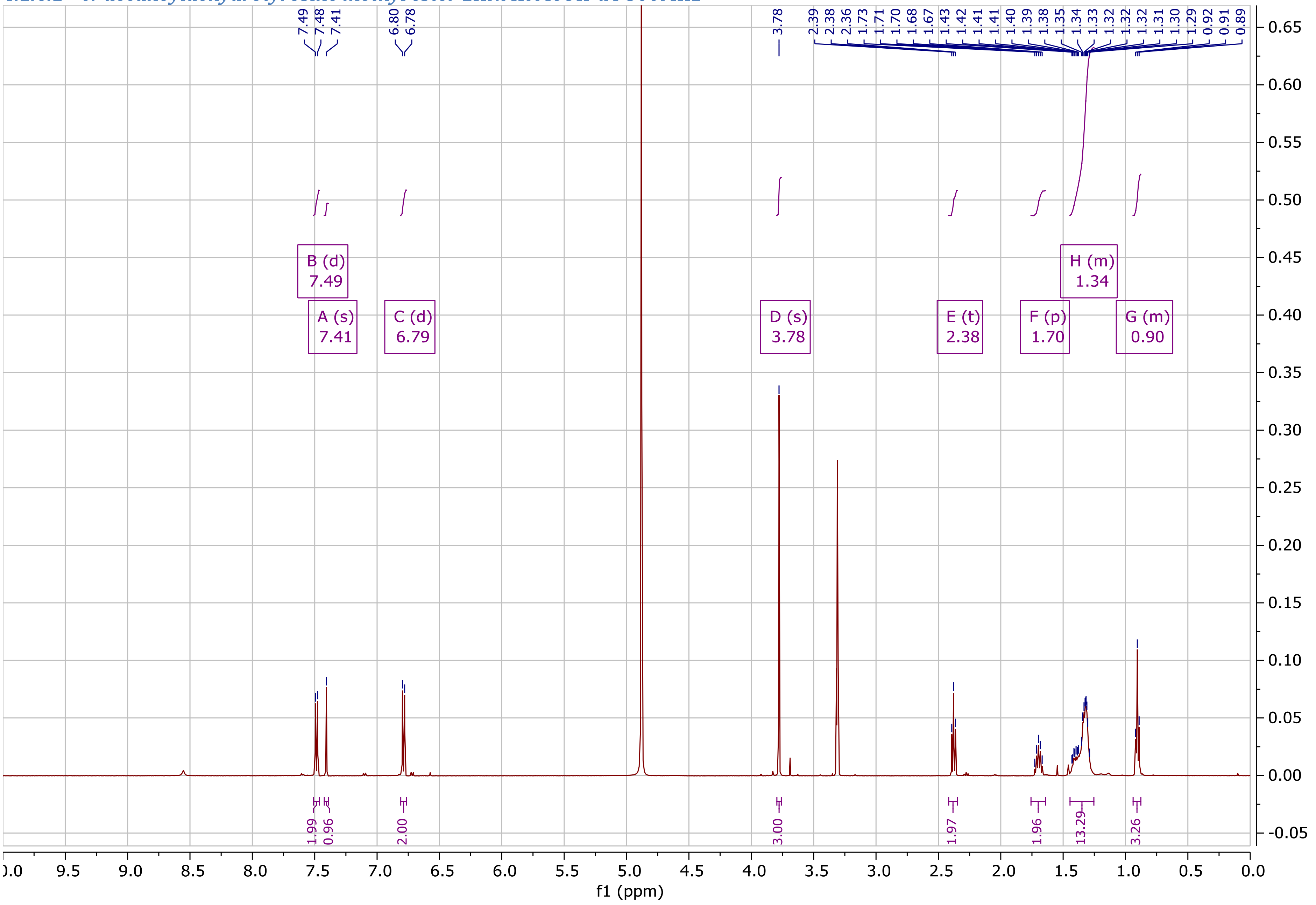

##### *4.1.6.2 N-decanoyldehydrotyrosine methyl ester 1HNMR DMSO 600MHz*

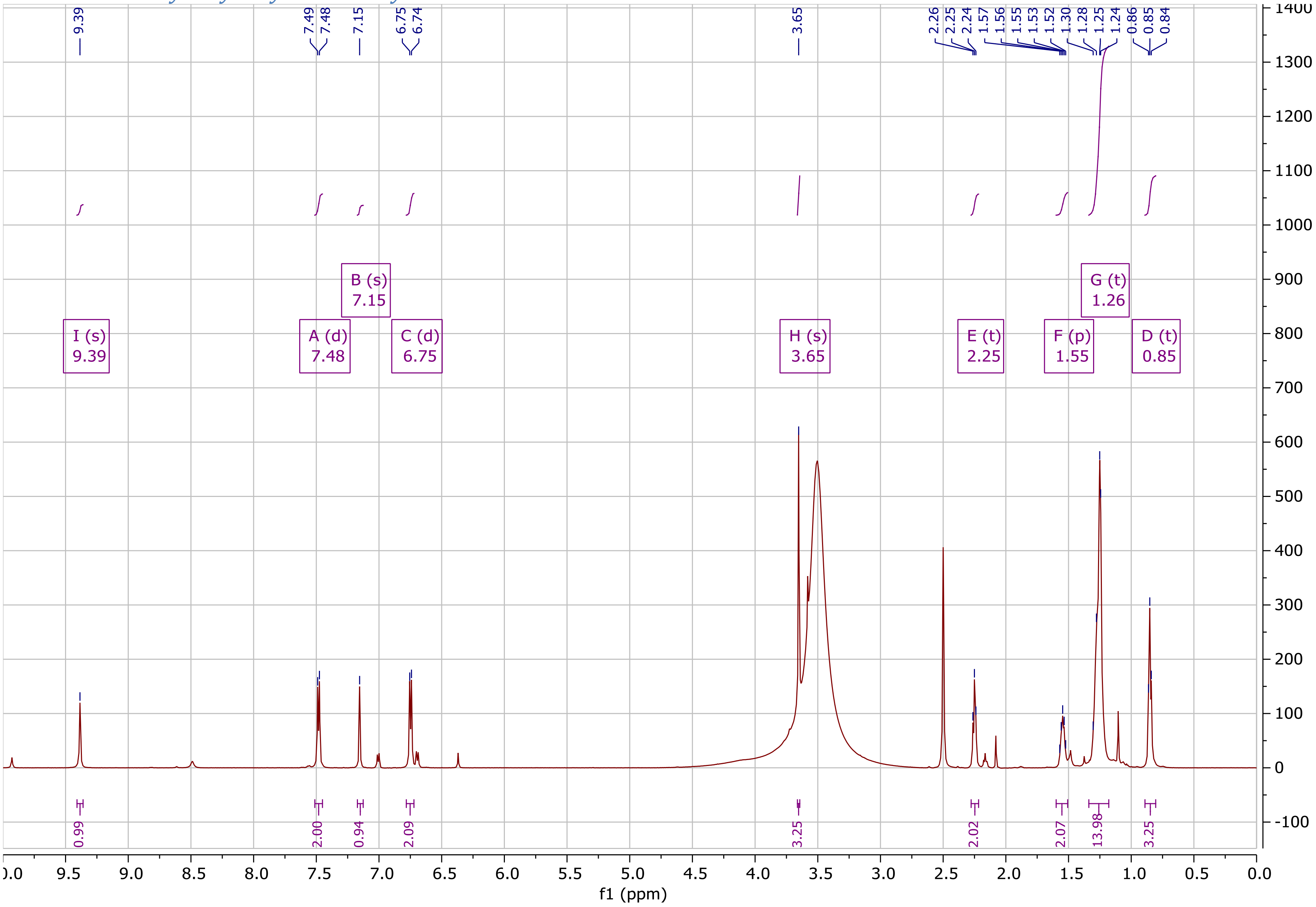

##### 4.1.7 N-octanoyldehydrophenylalanine methyl ester

###### *4.1.7.1 N-octanoyldehydrophenylalanine methyl ester 1HNMR DMSO 500MHz*

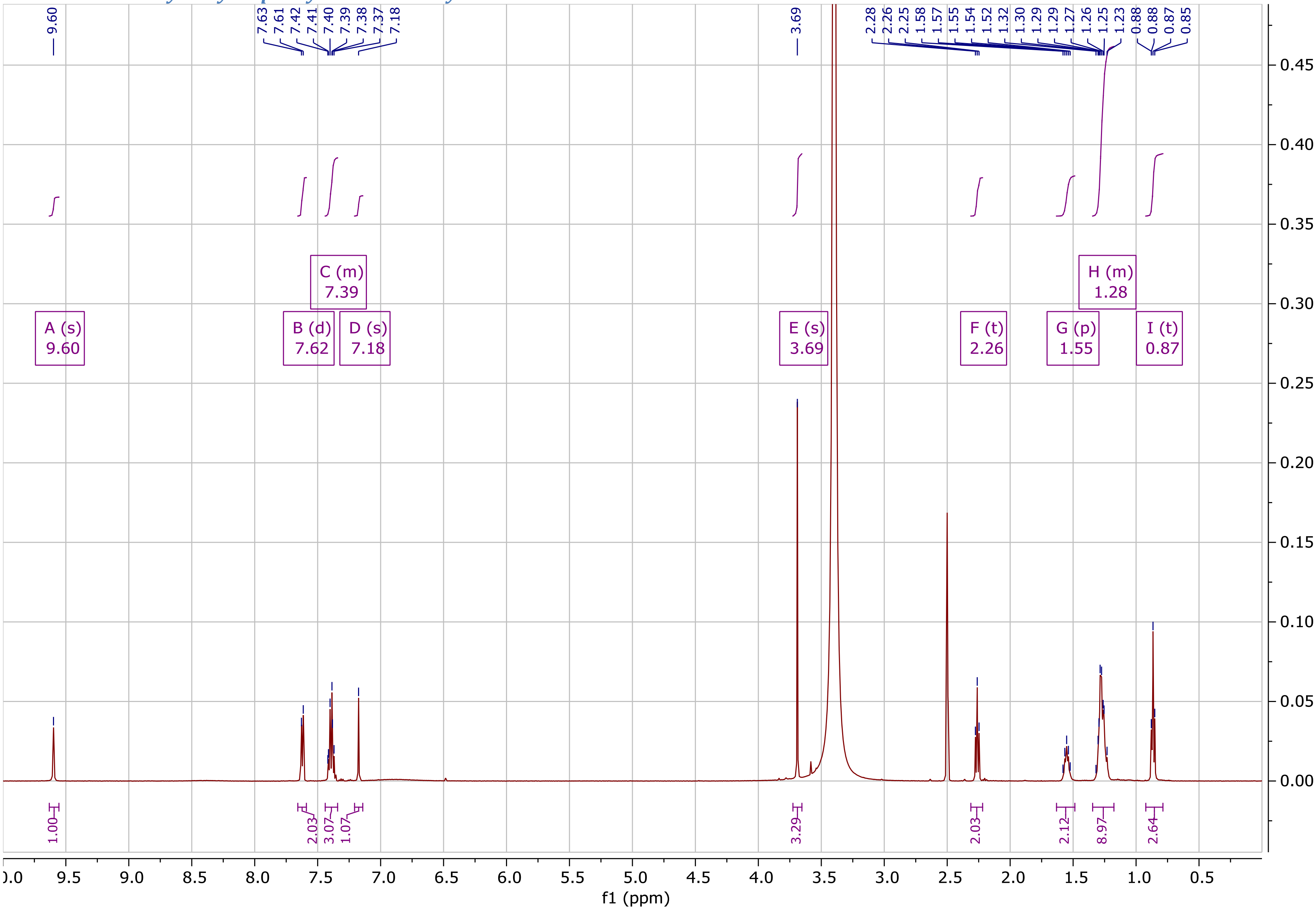

##### 4.1.8 N-decanoyltyrosine

###### *4.1.8.1 N-decanoyltyrosine 1HNMR DMSO-d6 500MHz*

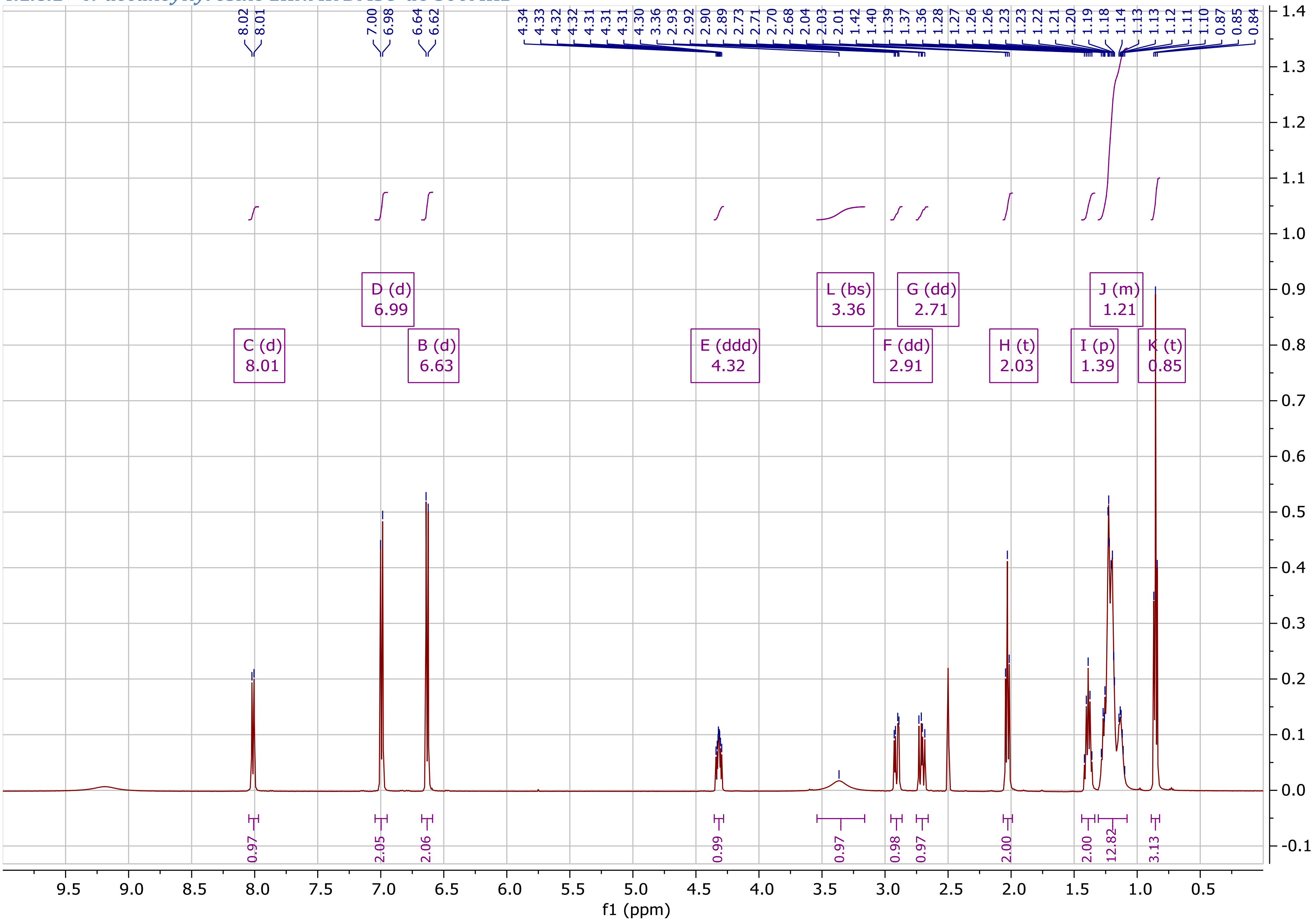

#### 4.2 $^{13}$C NMR

##### 4.2.1 Nonyltyrazolone

###### *4.2.1.1 Nonyltyrazolone 13CNMR CDCl3 500MHz*

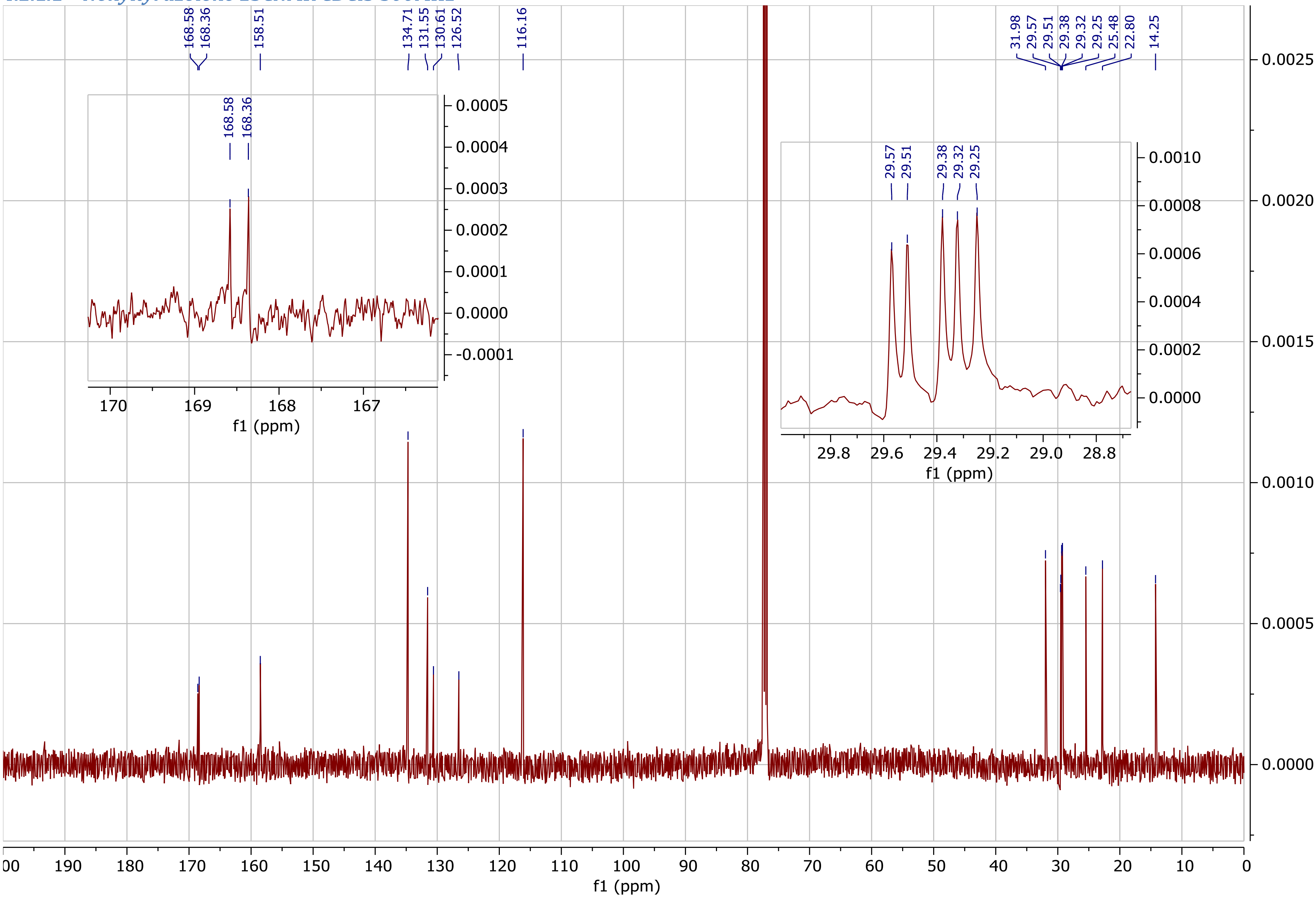

##### 4.2.2 Undecyltyrazolone

###### *4.2.2.1 Undecyltyrazolone 13CNMR DMSO-d6 600MHz*

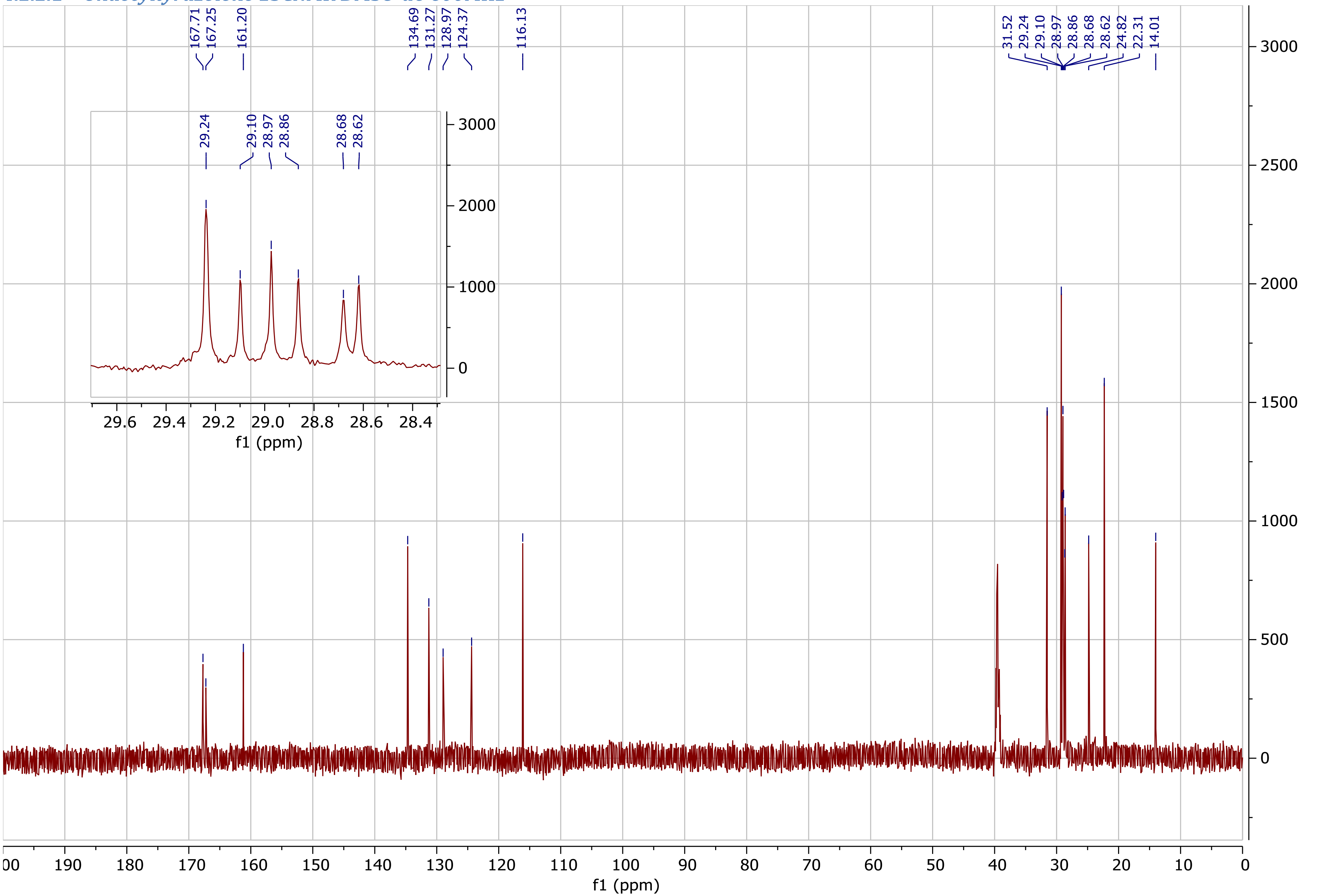

##### 4.2.3 Heptylphenazolone

###### *4.2.3.1 Heptylphenazolone 13CNMR CDCl3 500MHz*

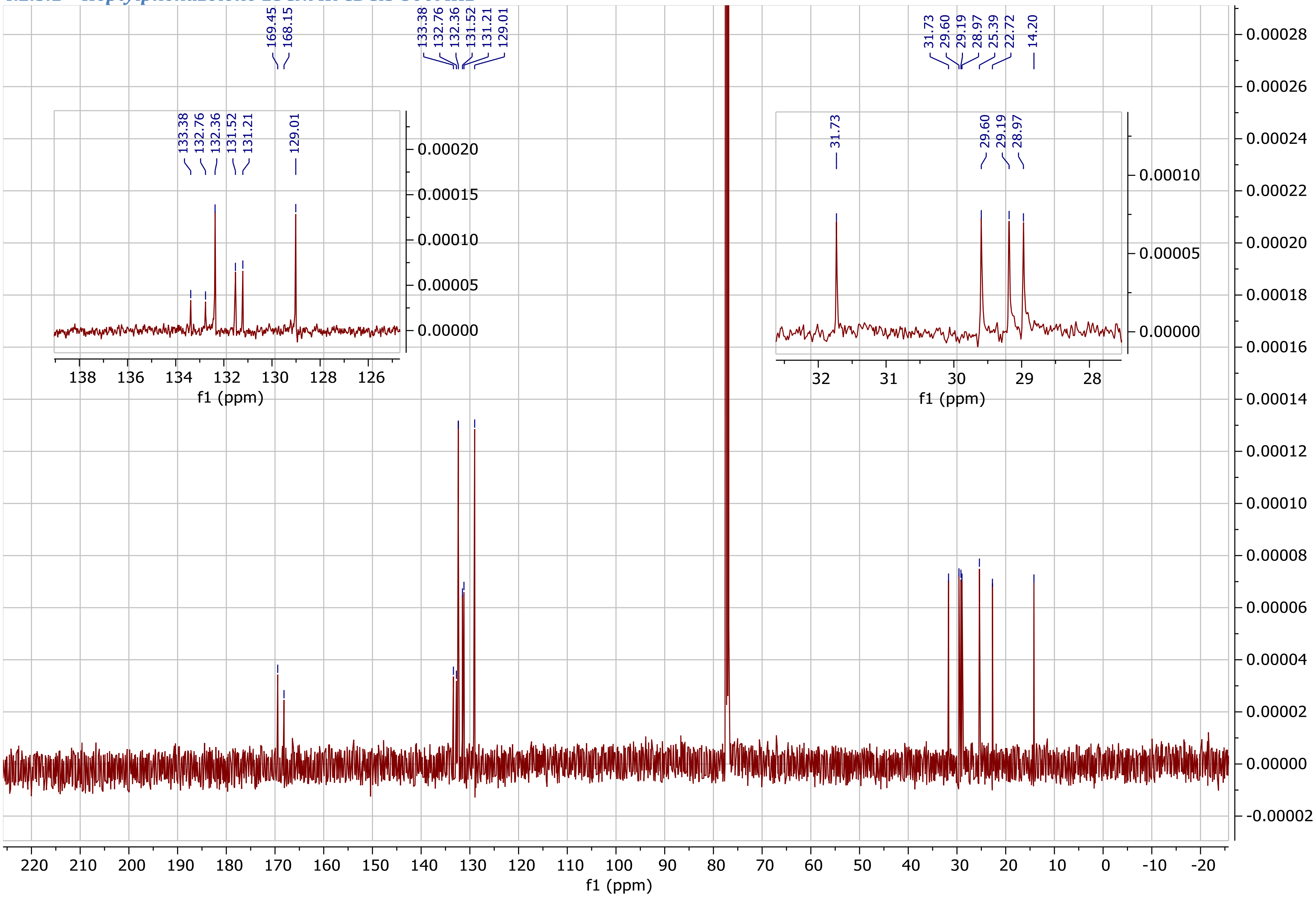

##### 4.2.4 N-decanoyldehydrotyrosine methyl ester

###### *4.2.4.1 N-decanoyldehydrotyrosine methyl ester 13CNMR MeOH-d4 500MHz*

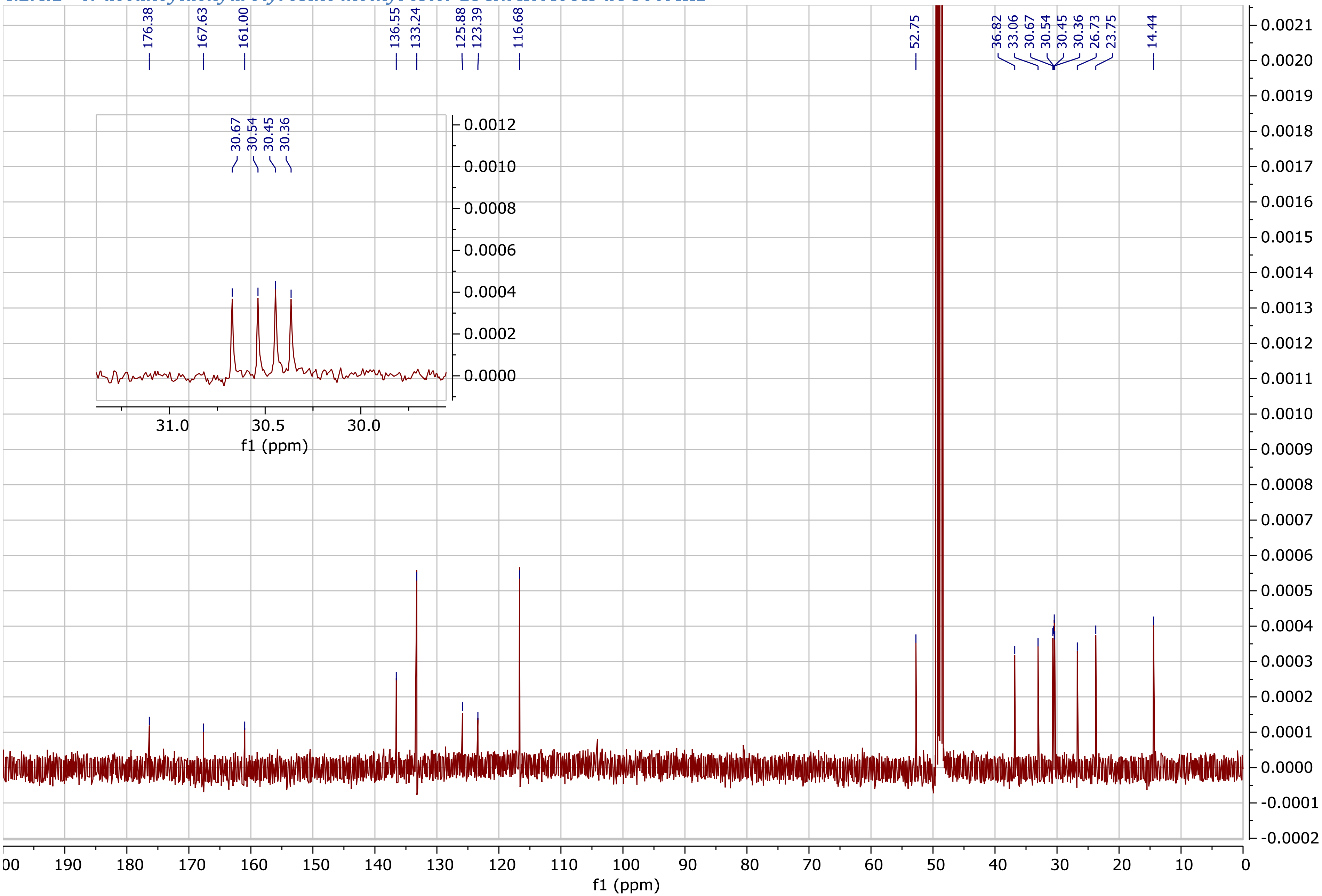

##### 4.2.5 N-octanoyldehydrophenylalanine methyl ester

##### 4.2.6 N-octanoyldehydrophenylalanine methyl ester 13CNMR DMSO 500MHz

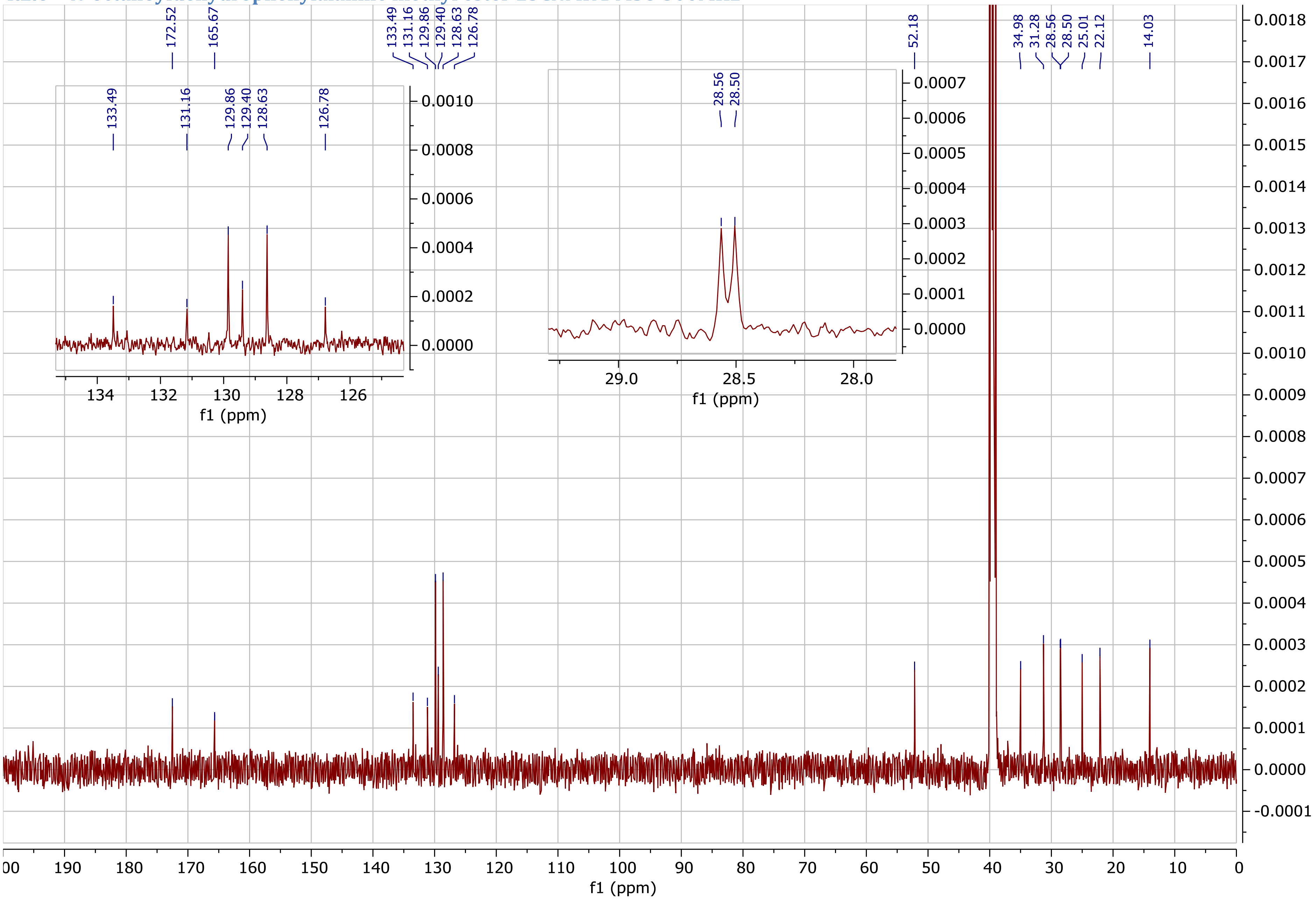

##### 4.2.7 N-decanoyltyrosine

###### *4.2.7.1 N-decanoyltyrosine 13CNMR DMSO 500MHz*

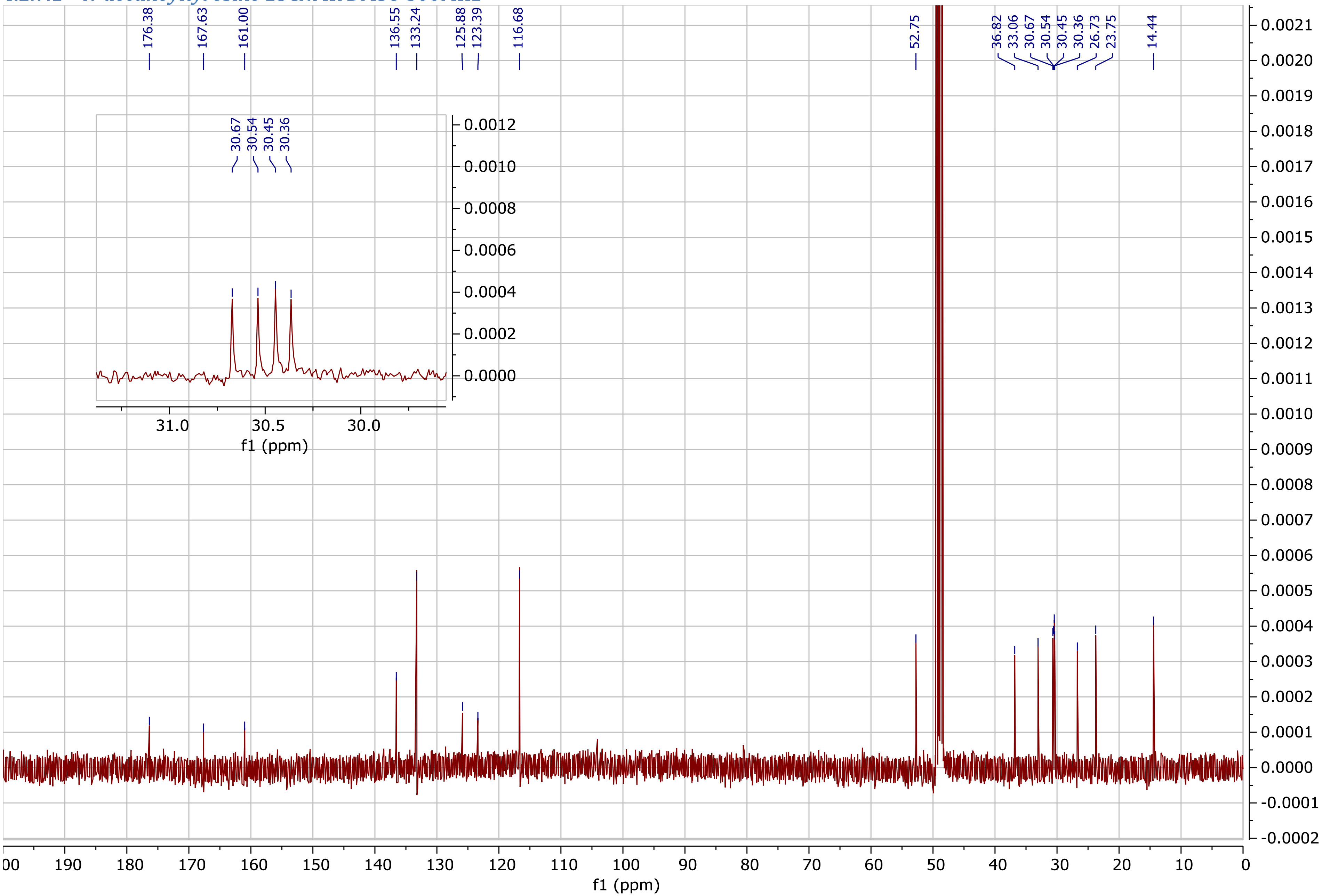

##### 4.3 $^{1}H$-$^{13}C$ HSQC and HMQC NMR

##### 4.3.1 Nonyltyrazolone

###### *4.3.1.1 Nonyltyrazolone 1H-13C HSQC CDCl3 500MHz*

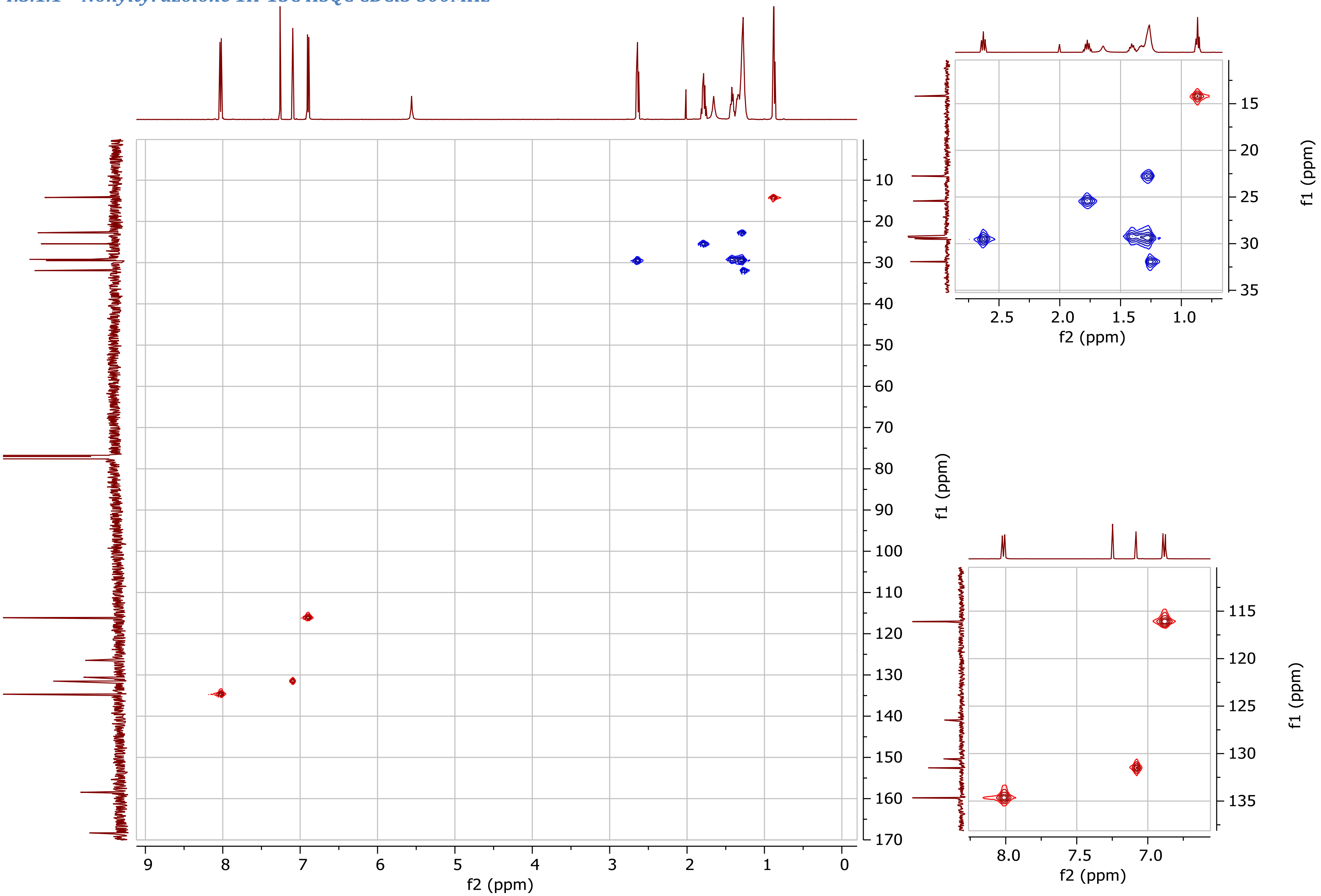

##### 4.3.2 Undecyltyrazolone

###### *4.3.2.1 Undecyltyrazolone 1H-13C HSQC DMSO 600MHz*

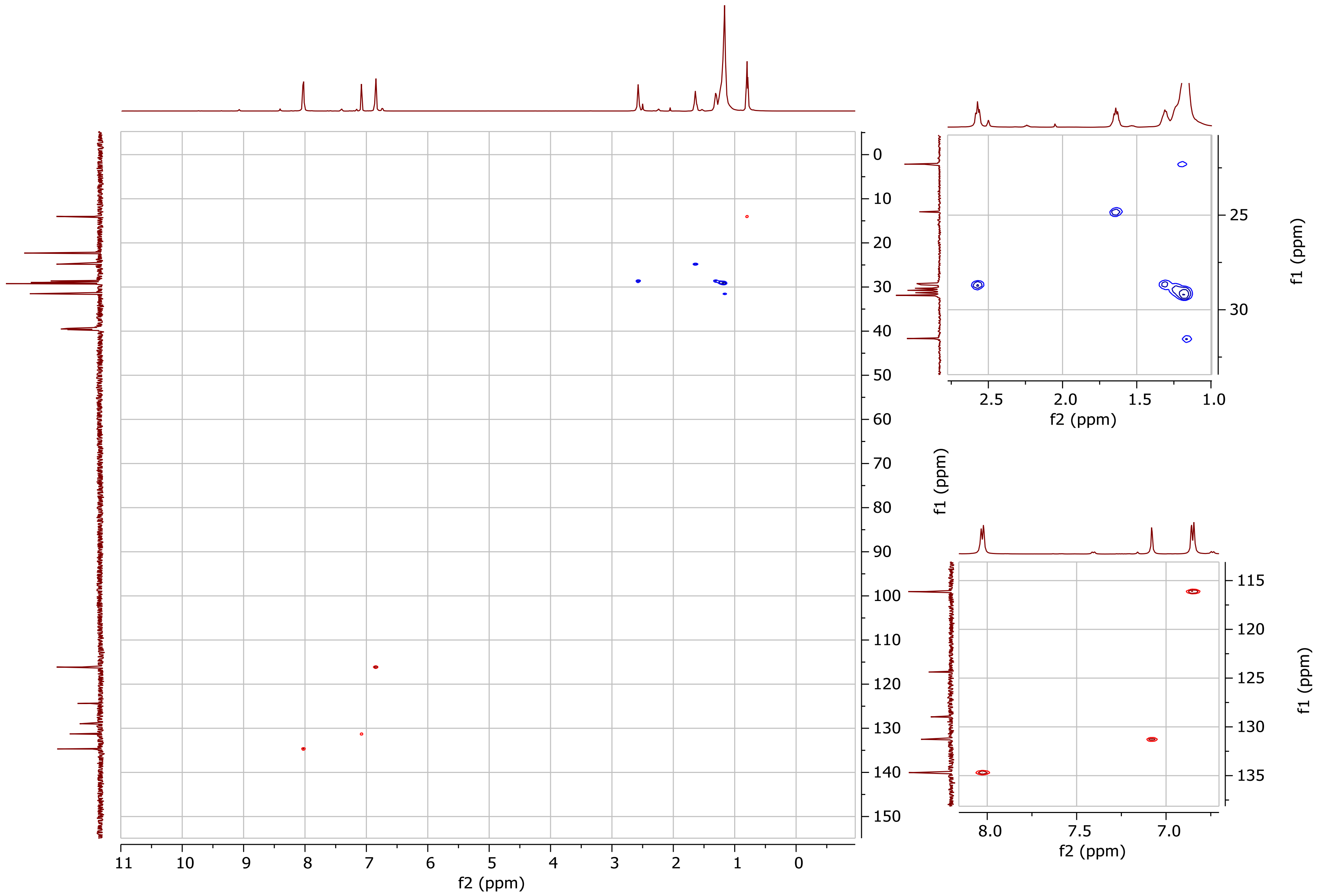

##### *4.3.2.2 Undecyltyrazolone 1H-13C HSQC CDCl3 600MHz*

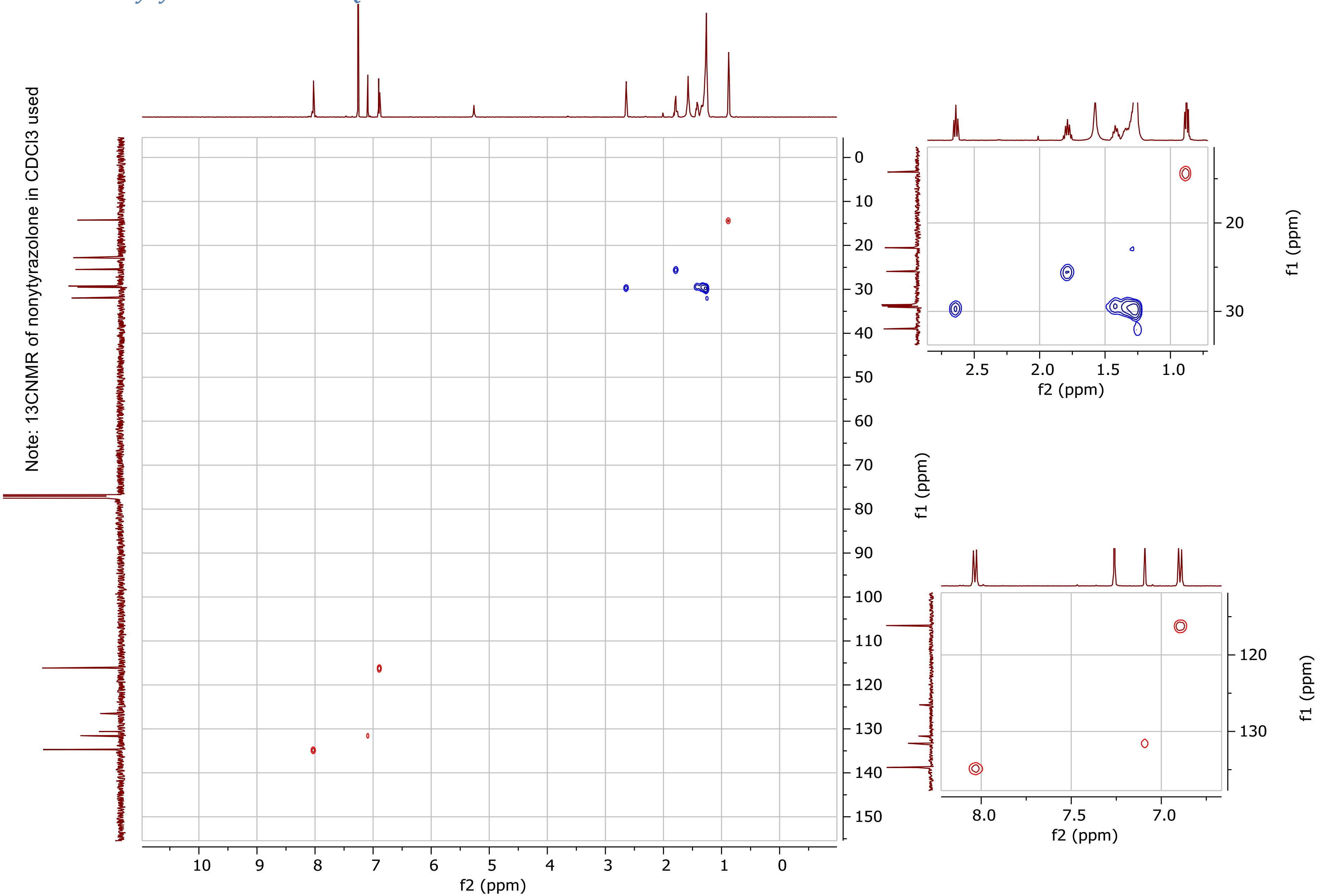

##### 4.3.3 Heptylphenazolone

###### *4.3.3.1 Heptylphenazolone 1H-13C HSQC CDCl3 500MHz*

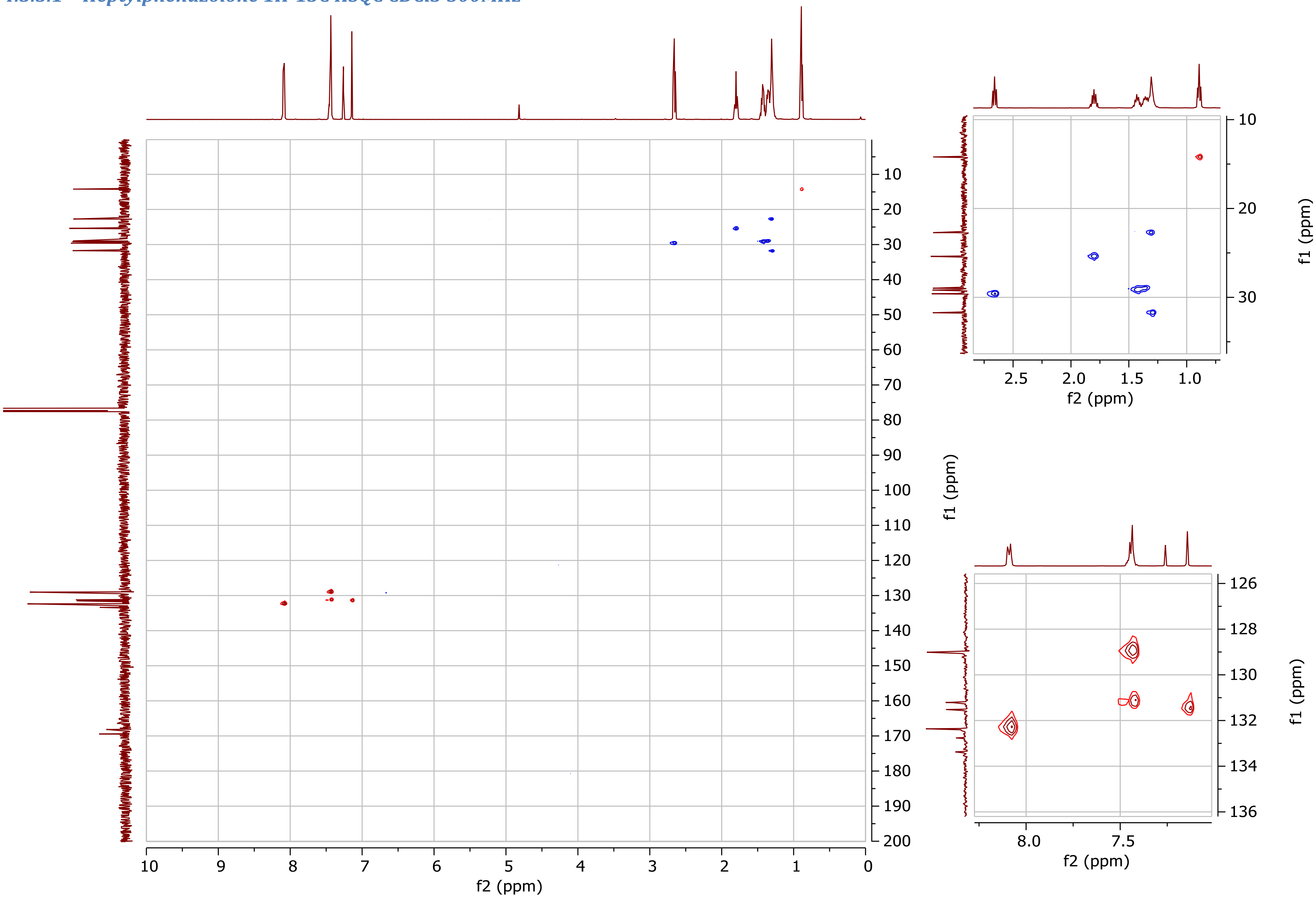

##### 4.3.4 N-decanoyldehydrotyrosine methyl ester

###### *4.3.4.1 N-decanoyldehydrotyrosine methyl ester 1H-13C HSQC DMSO 600MHz*

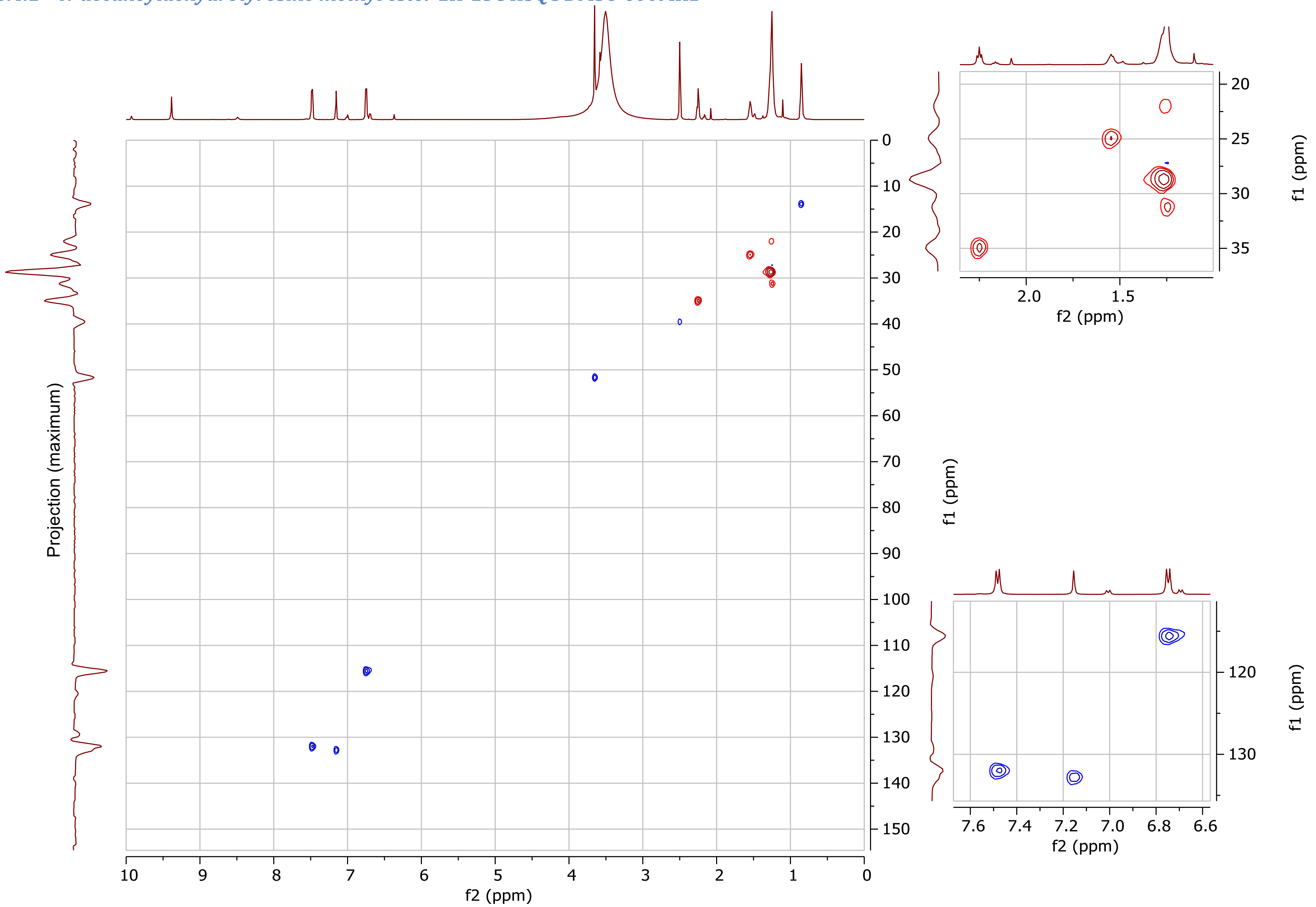

###### 4.3.4.2 N-decanoyldehydrotyrosine methyl ester 1H-13C HSQC MeOH-d4 500MHz (contaminated with ethyl acetate)

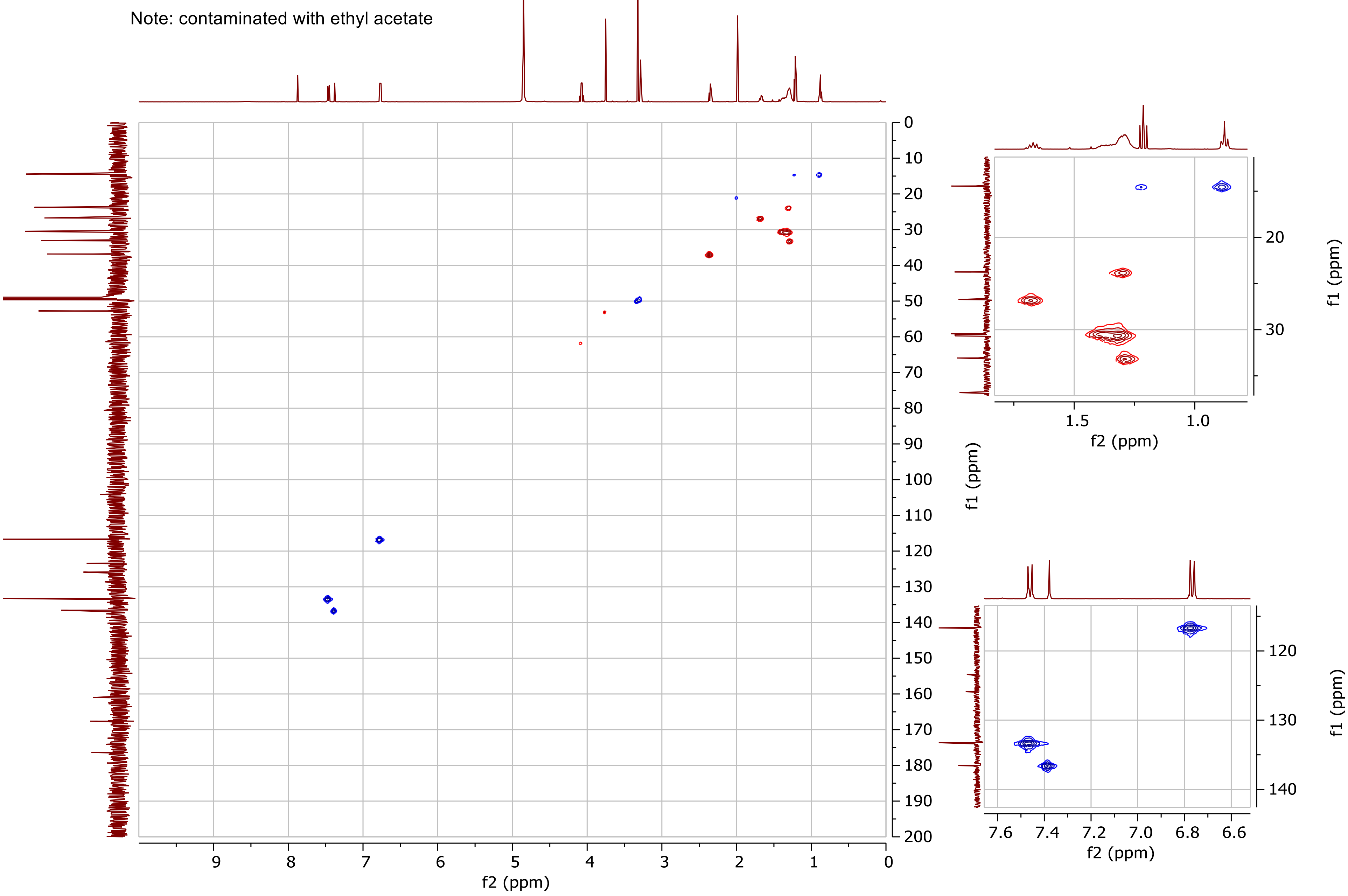

##### 4.3.5 N-octanoyldehydrophenylalanine methyl ester

###### *4.3.5.1 N-octanoyldehydrophenylalanine methyl ester 1H-13C HMQC DMSO-d6 500MHz*

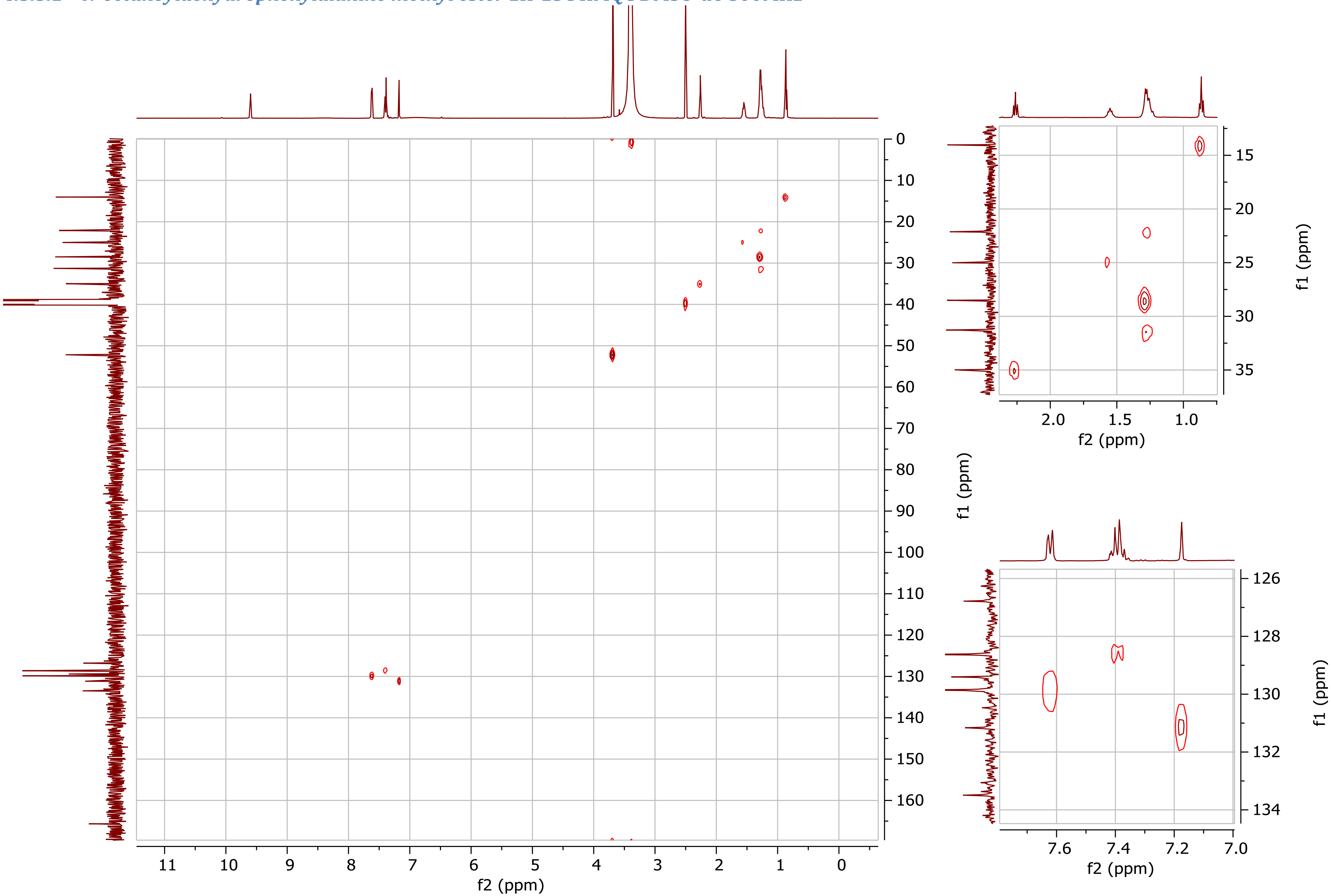

#### 4.4 $^{1}H$-$^{13}C$ HMBC NMR

#### 4.4.1 Nonyltyrazolone

##### *4.4.1.1 Nonyltyrazolone 1H-13C HMBC CDCl3 500MHz*

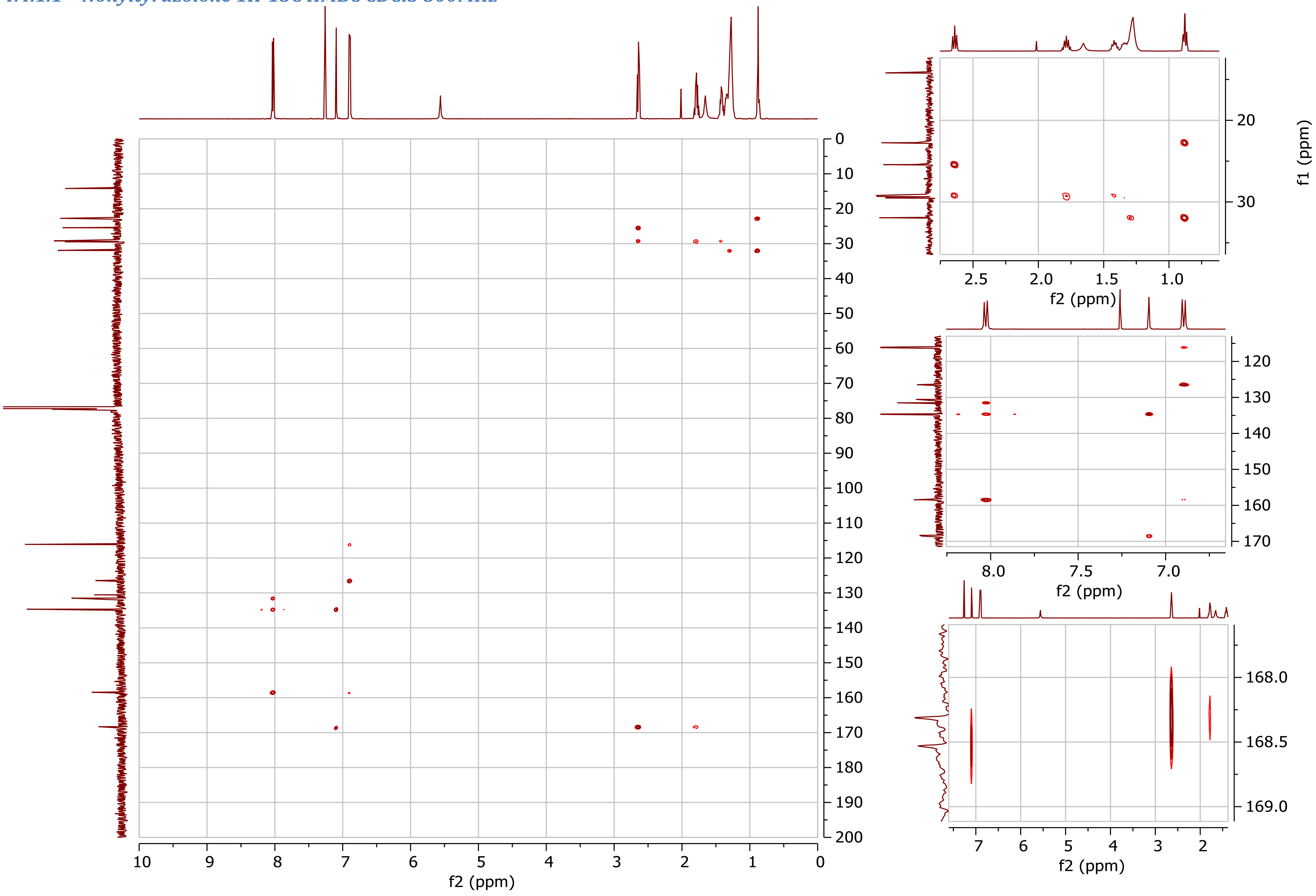

#### 4.4.2 Undecyltyrazolone

##### *4.4.2.1 Undecyltyrazolone 1H-13C HMBC DMSO-d6 600MHz*

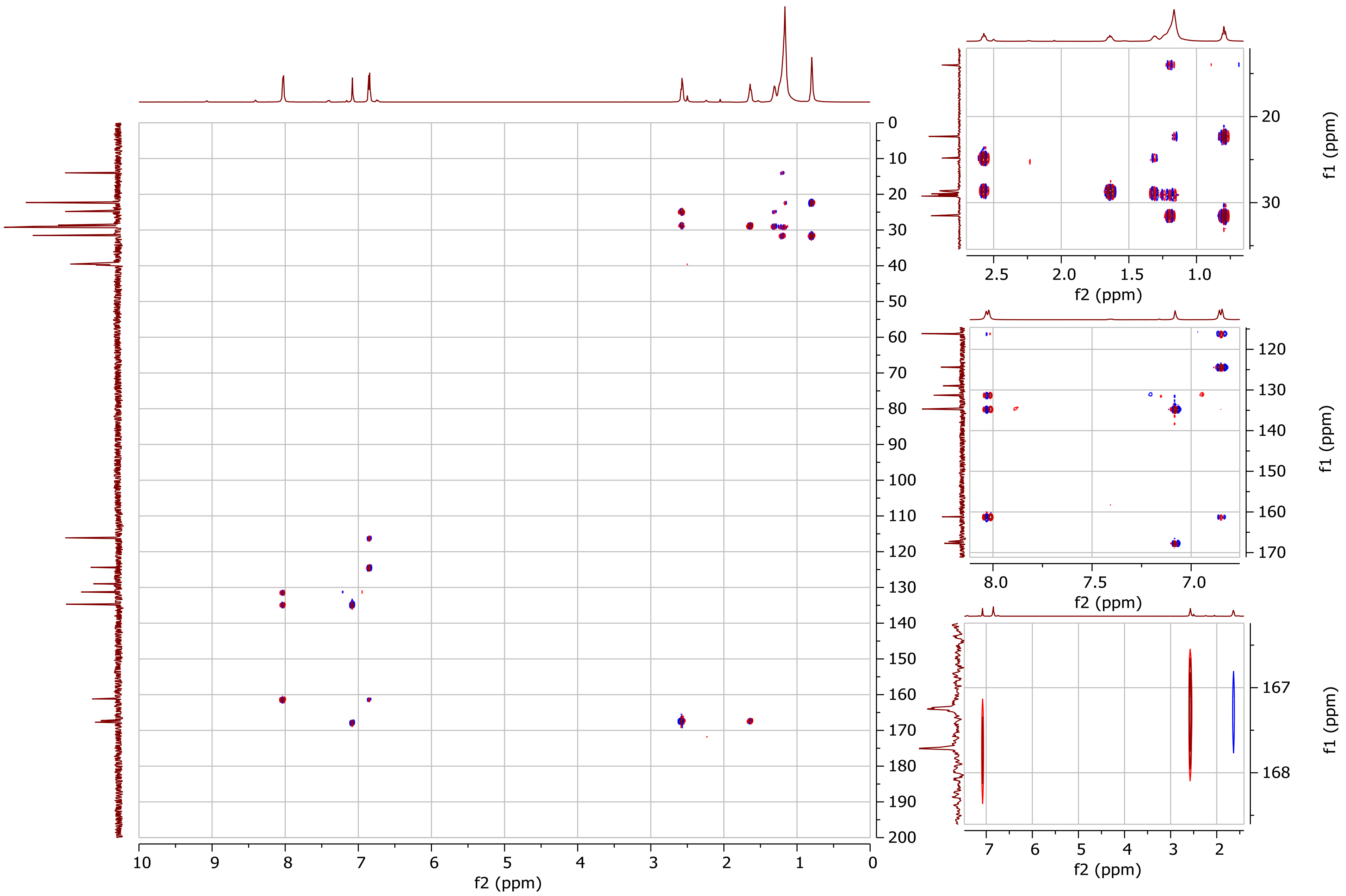

##### *4.4.2.2 Undecyltyrazolone 1H-13C HMBC CDCl3 600MHz*

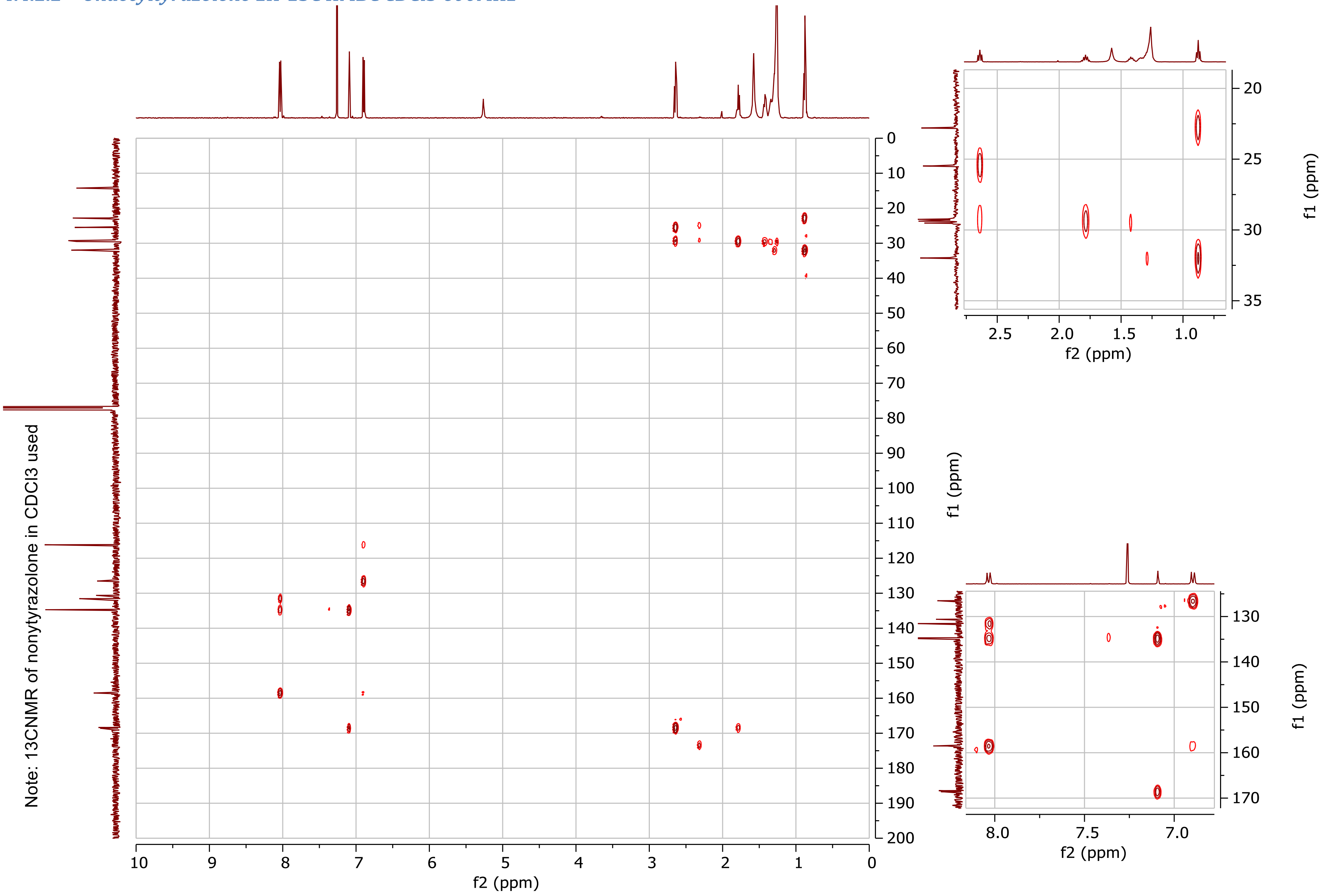

##### 4.4.3 ω-6-undecenyltyrazolone

###### *4.4.3.1 ω-6-undecenyltyrazolone 1H-13C HMBC CDCl 500MHz*

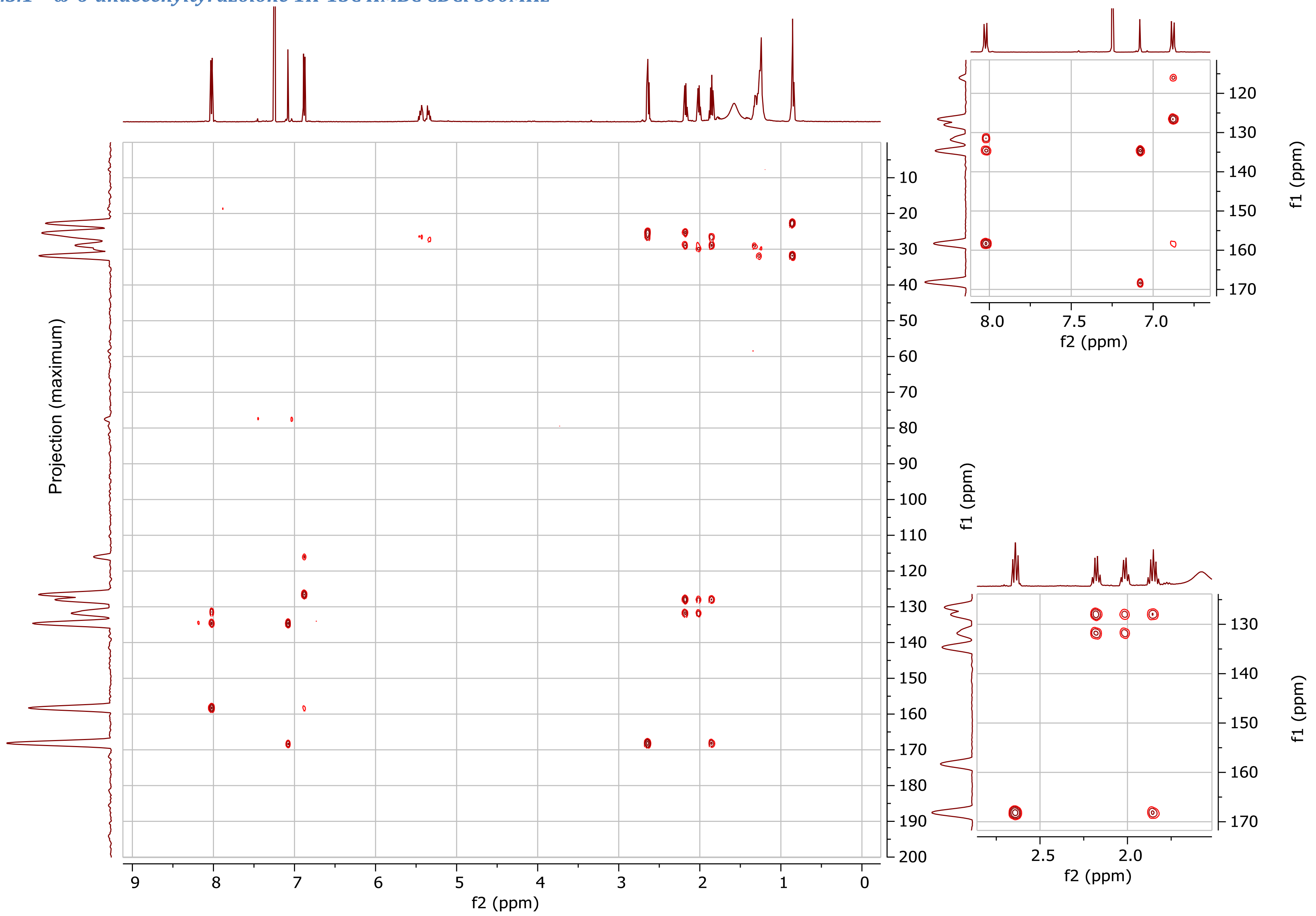

##### 4.4.4 Heptylphenazolone

###### *4.4.4.1 Heptylphenazolone 1H-13C HMBC CDCl3 500MHz*

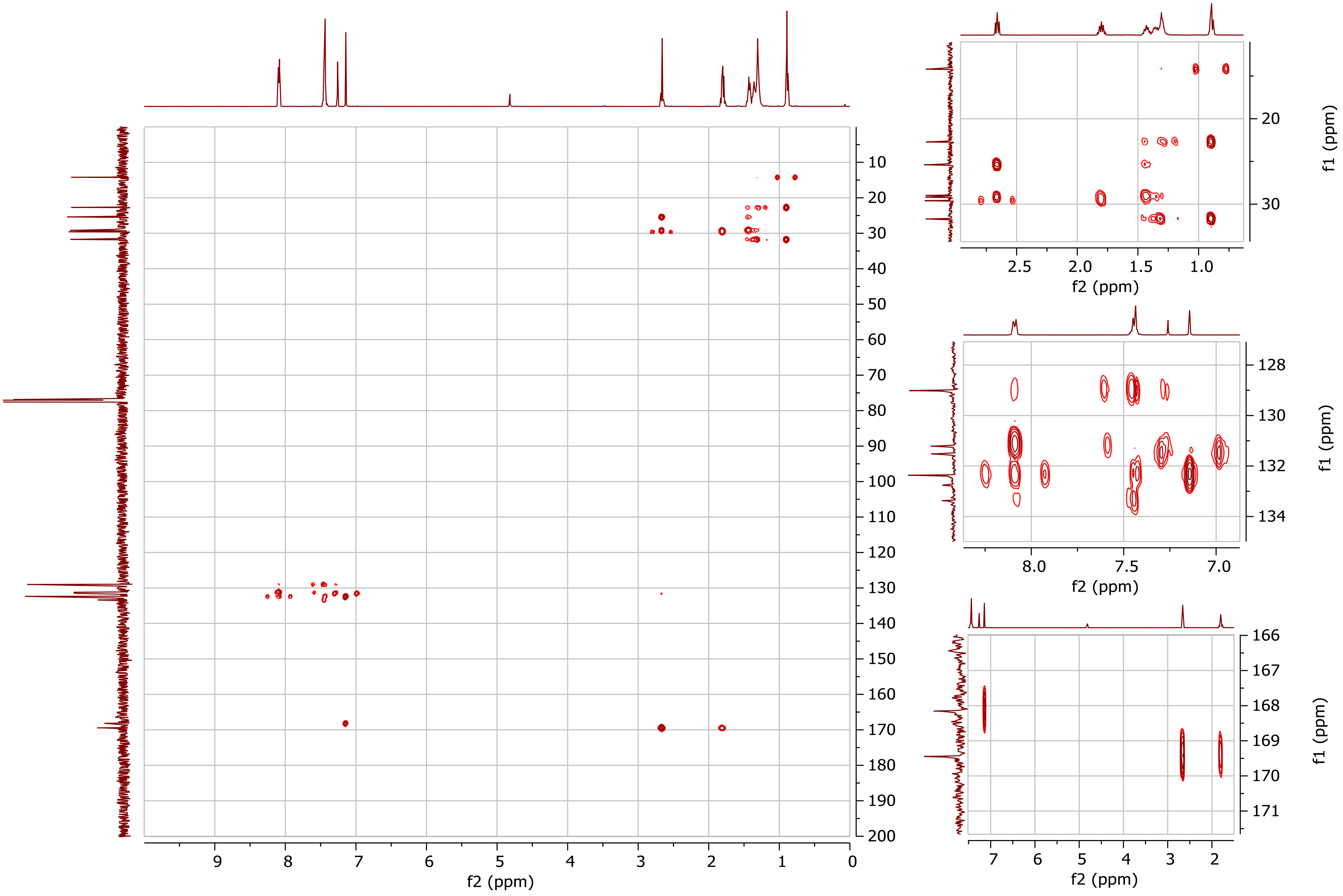

##### 4.4.5 N-decanoyldehydrotyrosine methyl ester

###### *4.4.5.1 N-decanoyldehydrotyrosine methyl ester 1H-13C HMBC MeOH-d4 500MHz*

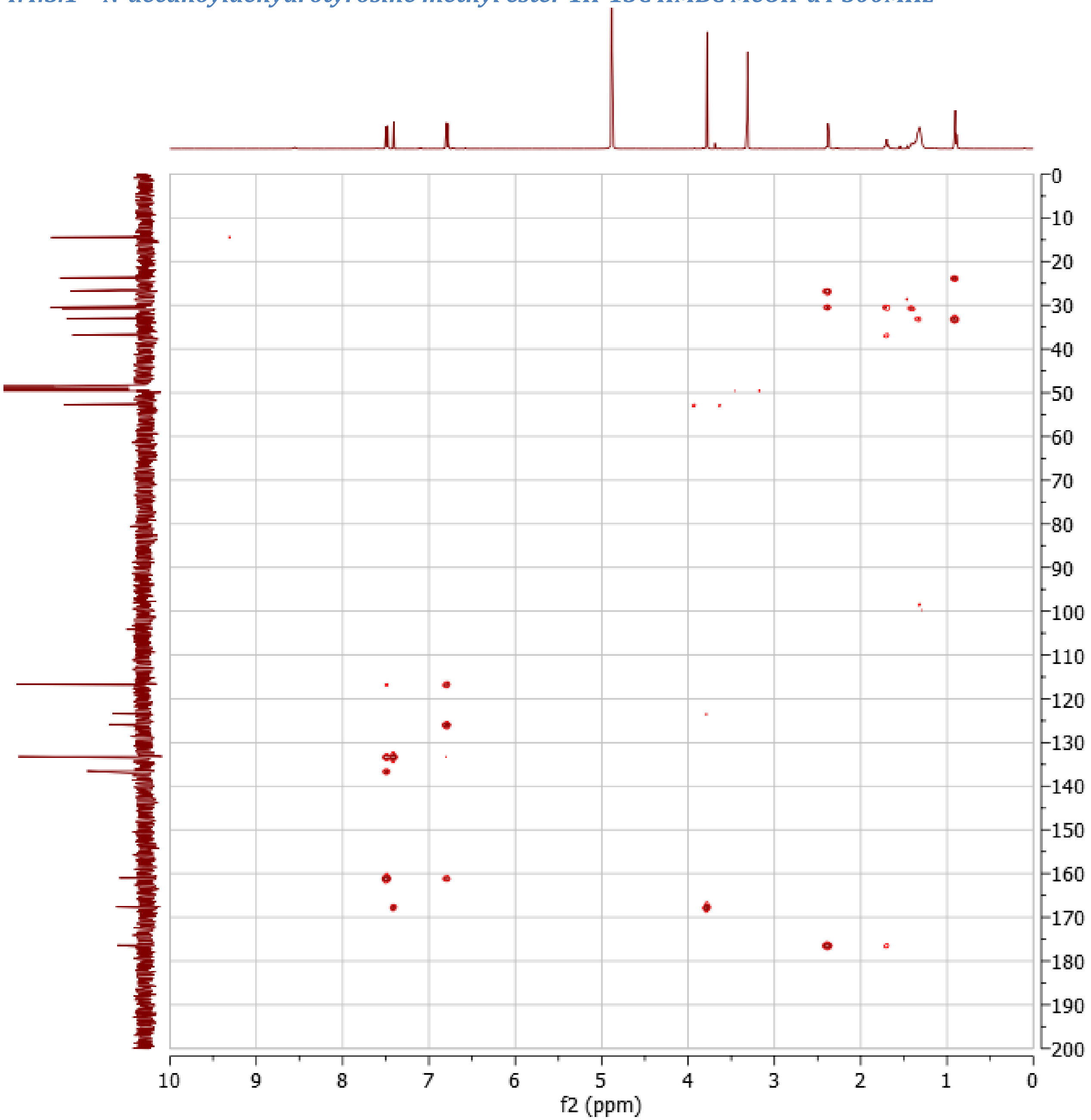

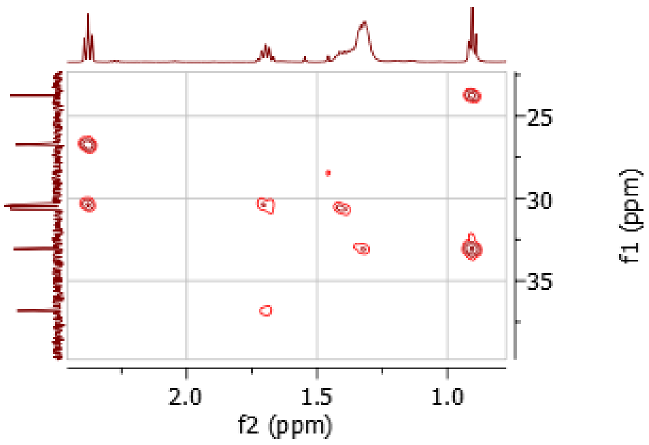

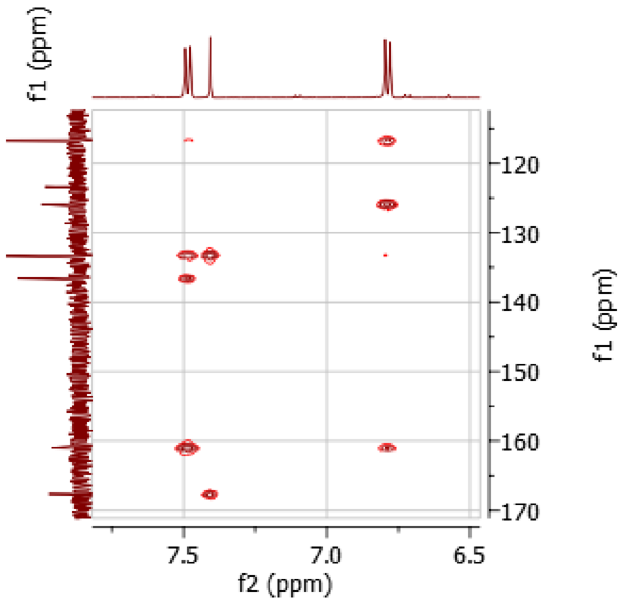

##### 4.4.5.2 N-decanoyldehydrotyrosine methyl ester 1H-13C HMBC DMSO 600MHz

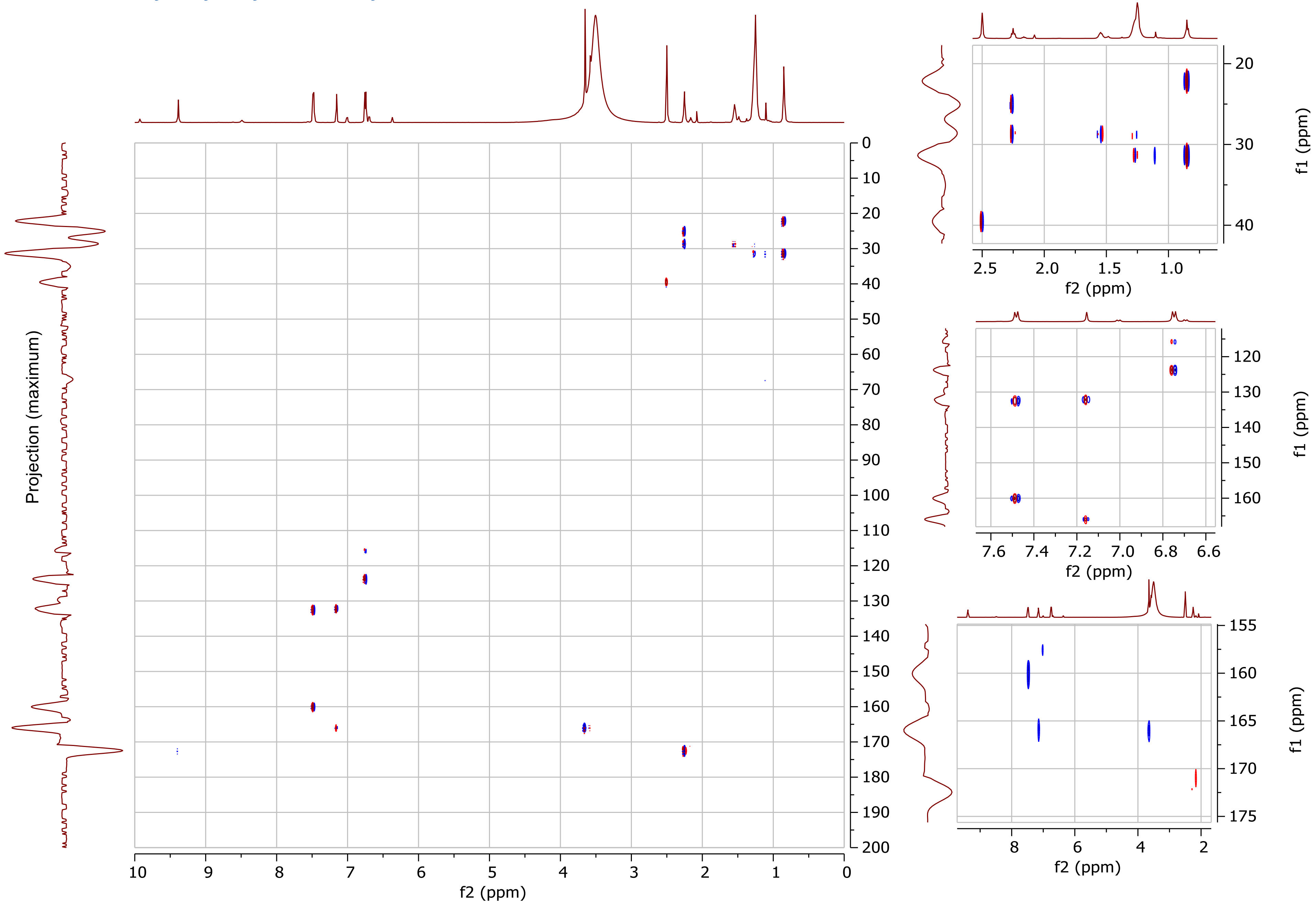

##### 4.4.6 N-octanoyldehydrophenylalanine methyl ester

###### *4.4.6.1 N-octanoyldehydrophenylalanine methyl ester 1H-13C HMBC DMSO-d6 500MHz*

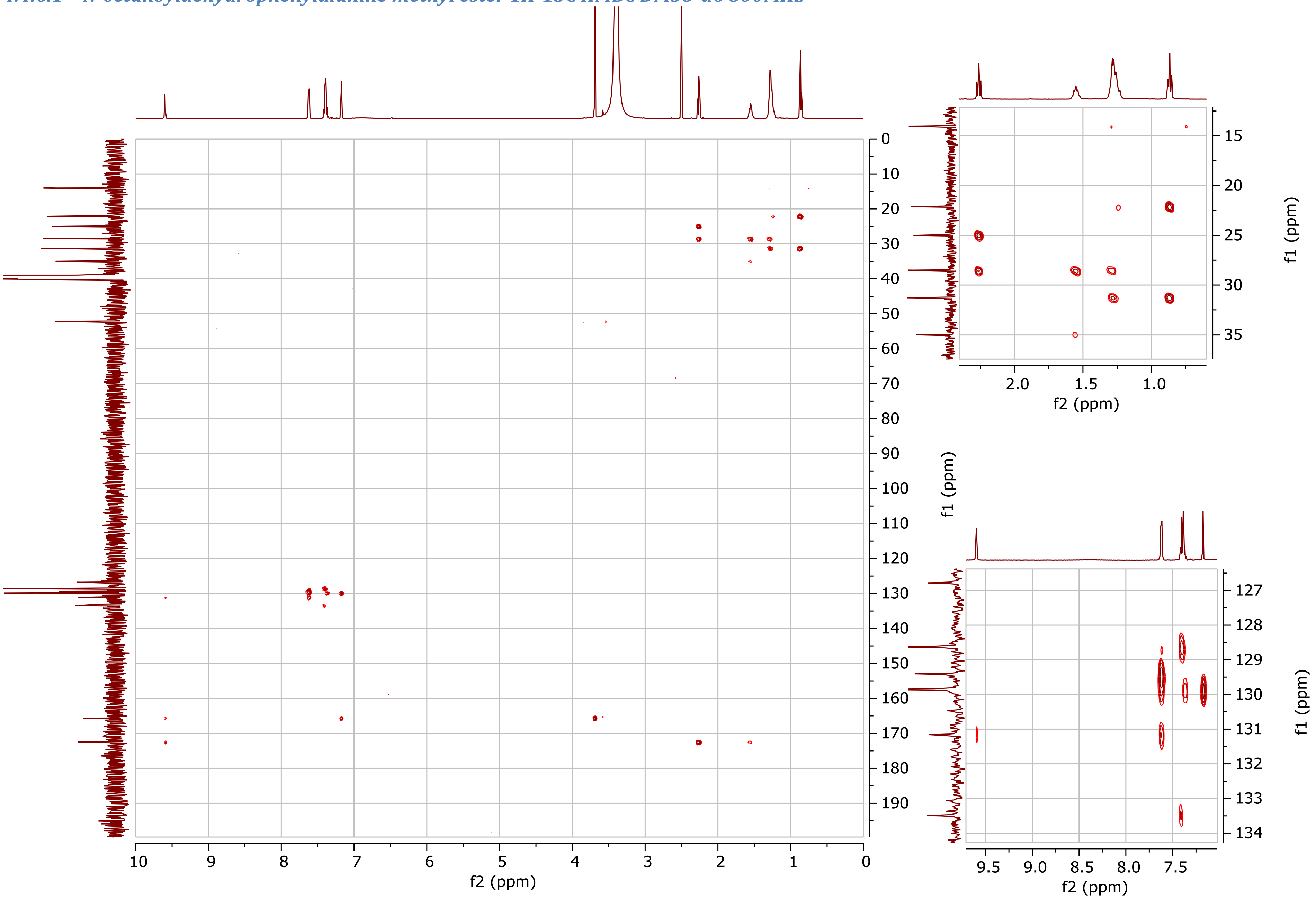

#### 4.5 $^{1}H$-$^{13}C$ ADEQUATE NMR

##### 4.5.1 Undecyltyrazolone

###### *4.5.1.1 Undecyltyrazolone 1H-13C ADEQUATE J=45Hz DMSO-d6 600MHz*

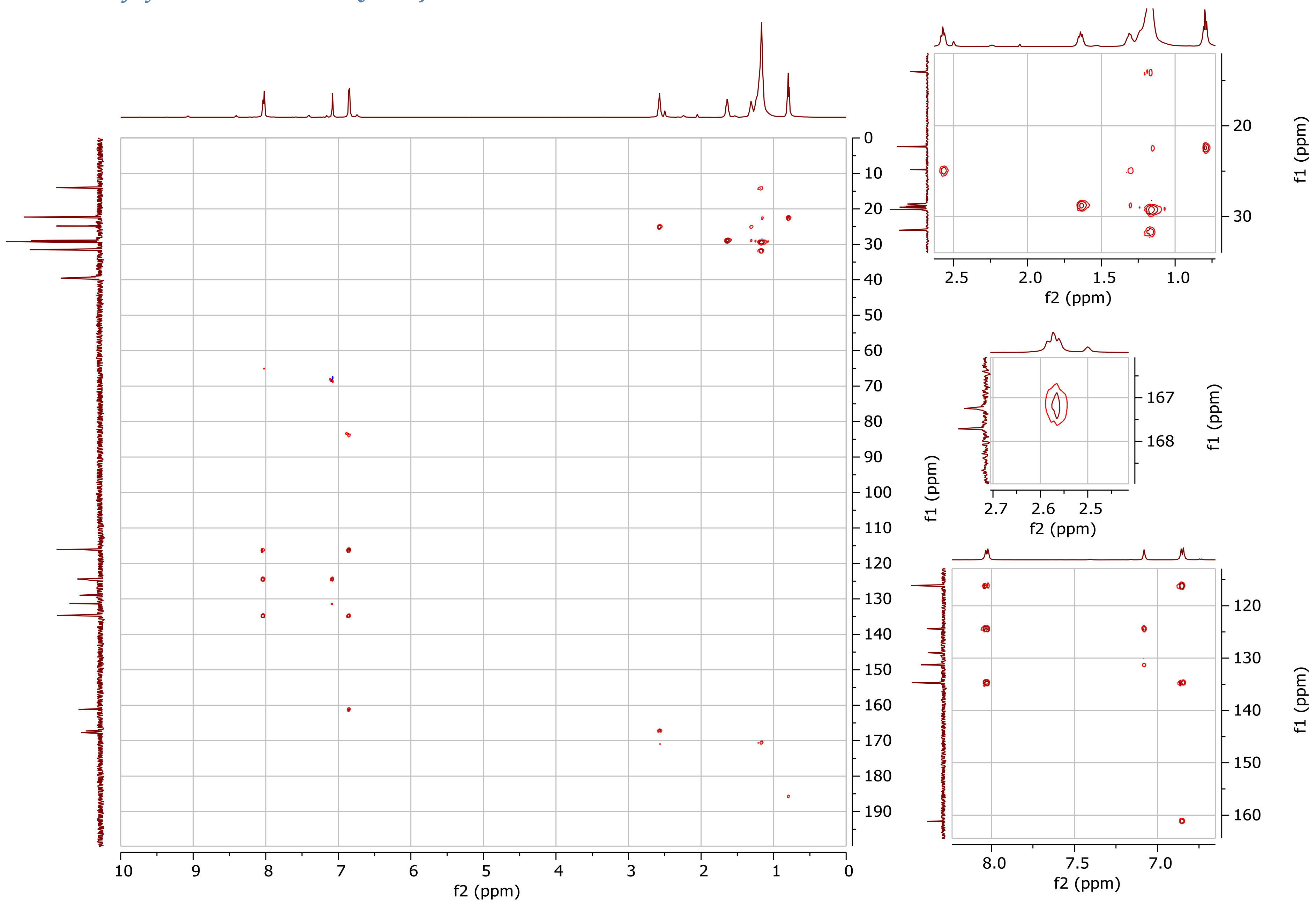

##### *4.5.1.2 Undecyltyrazolone 1H-13C ADEQUATE J=70Hz DMSO-d6 600MHz*

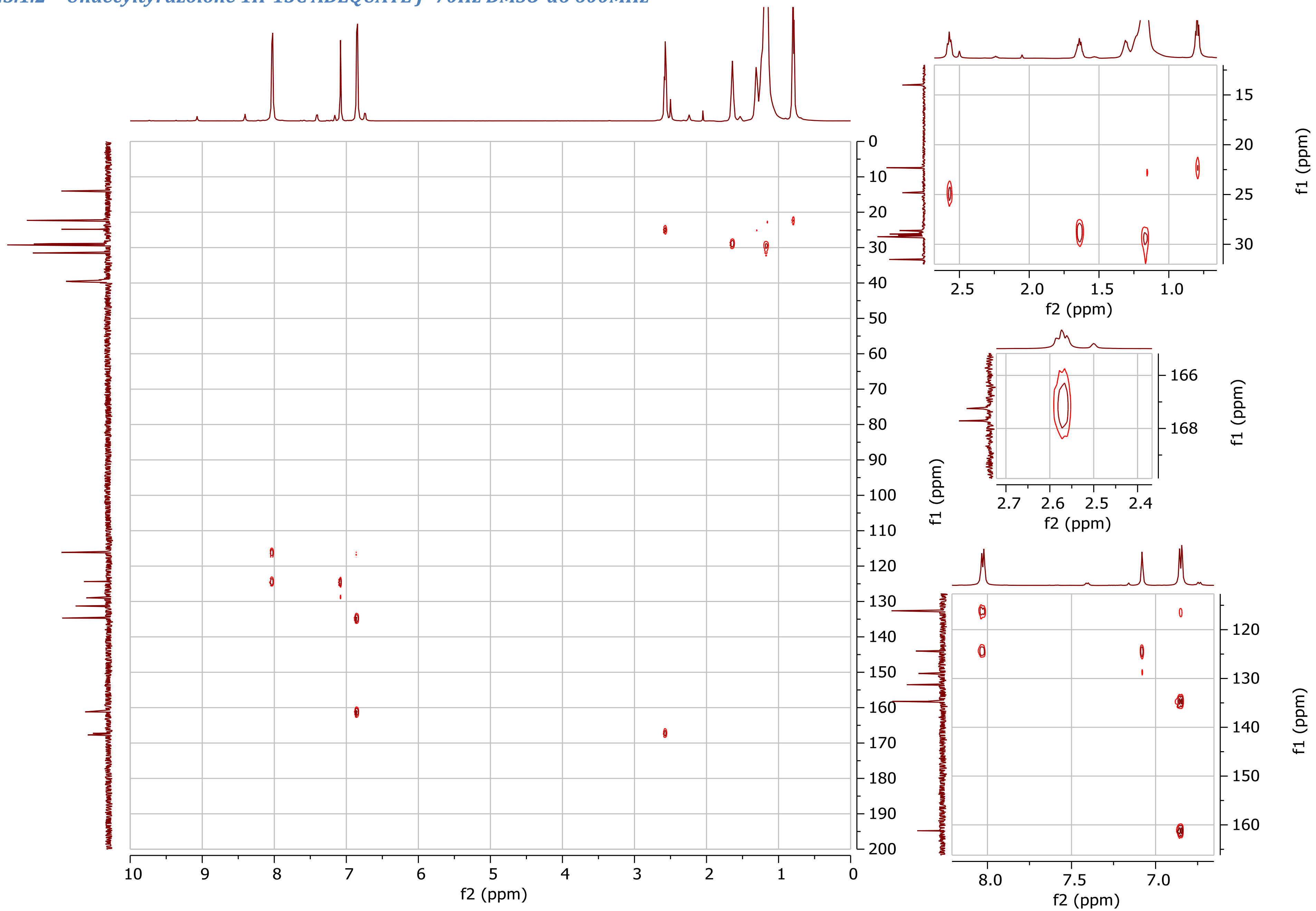

#### 4.6 1H-15N HMBC NMR

#### 4.6.1 Undecyltyrazolone

##### *4.6.1.1 Undecyltyrazolone 1H-15N HMBC DMSO-d6 600MHz*

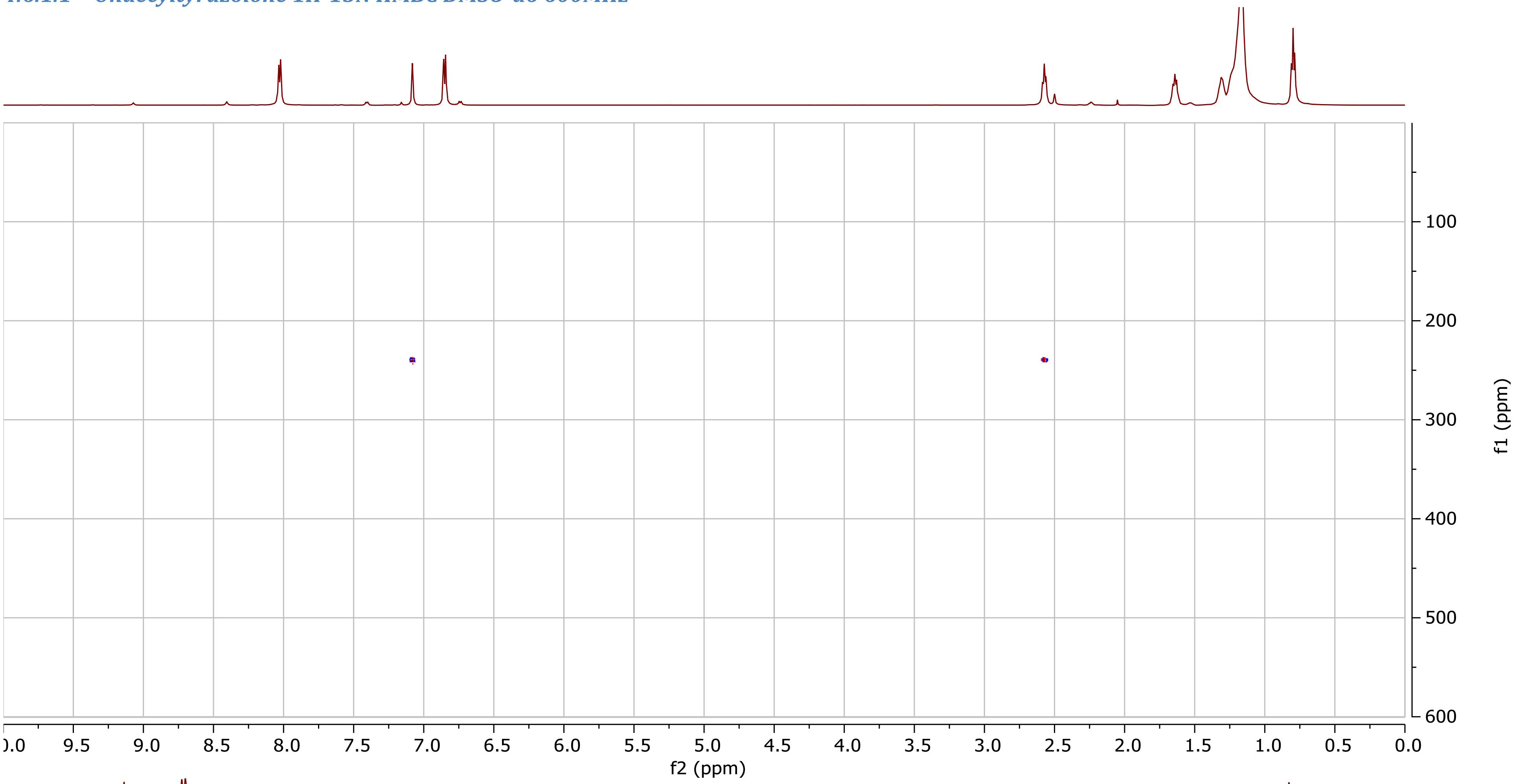

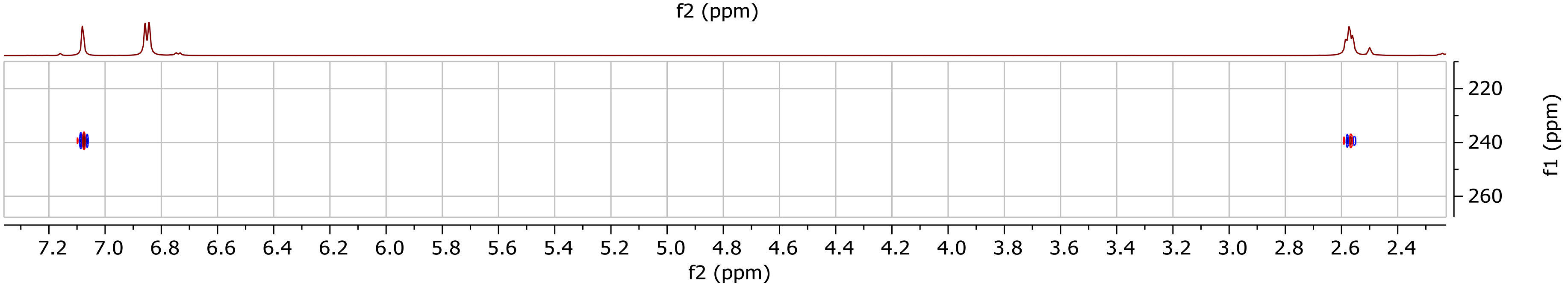

#### 4.7 $^{1}H$ COSY NMR

##### 4.7.1 Undecyltyrazolone

###### *4.7.1.1 Undecyltyrazolone 1H COSY DMSO-d6 600MHz*

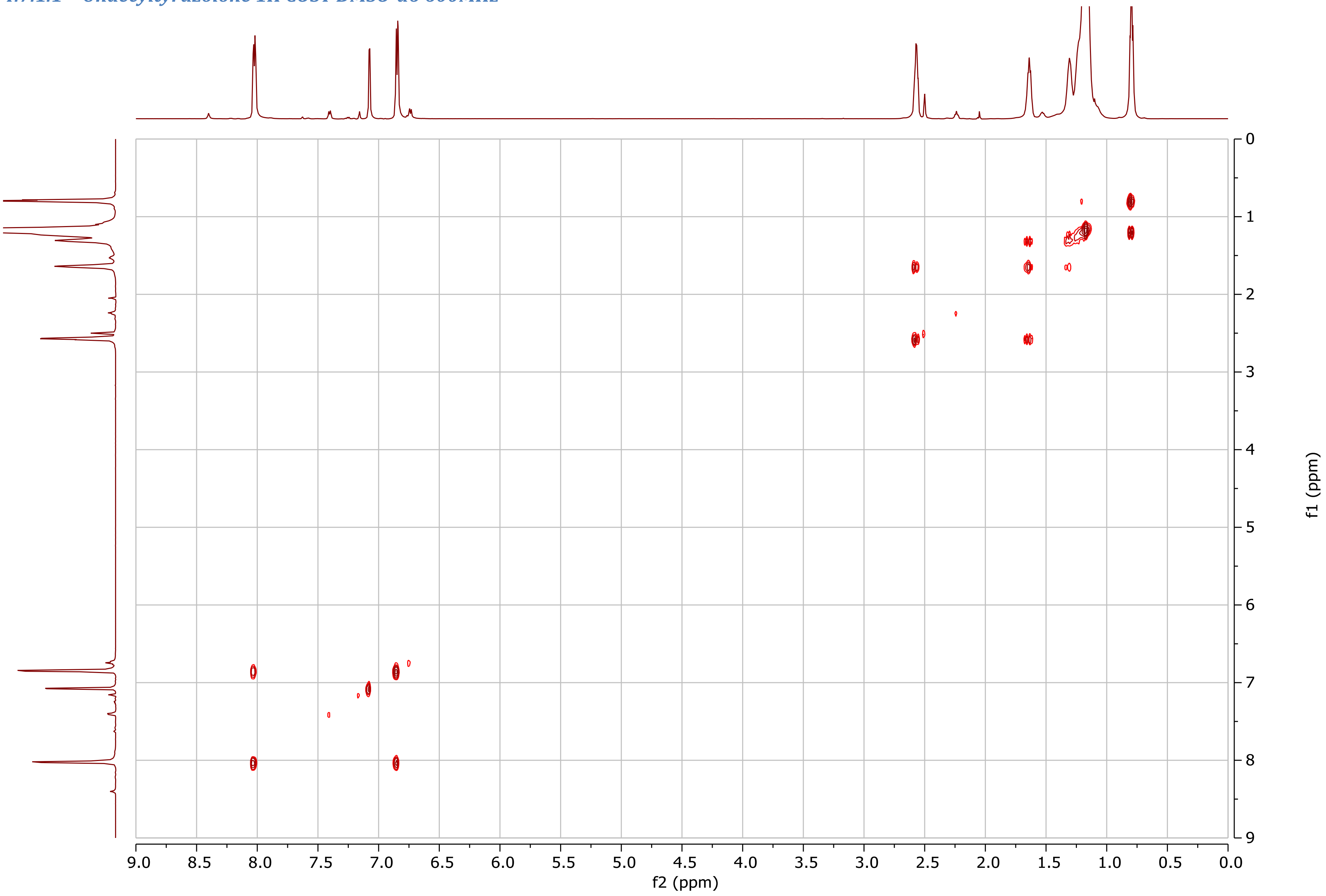

##### 4.7.2 ω-6-undecenyltyrazolone

###### *4.7.2.1 ω-6-undecenyltyrazolone 1H COSY CDCl3 500MHz*

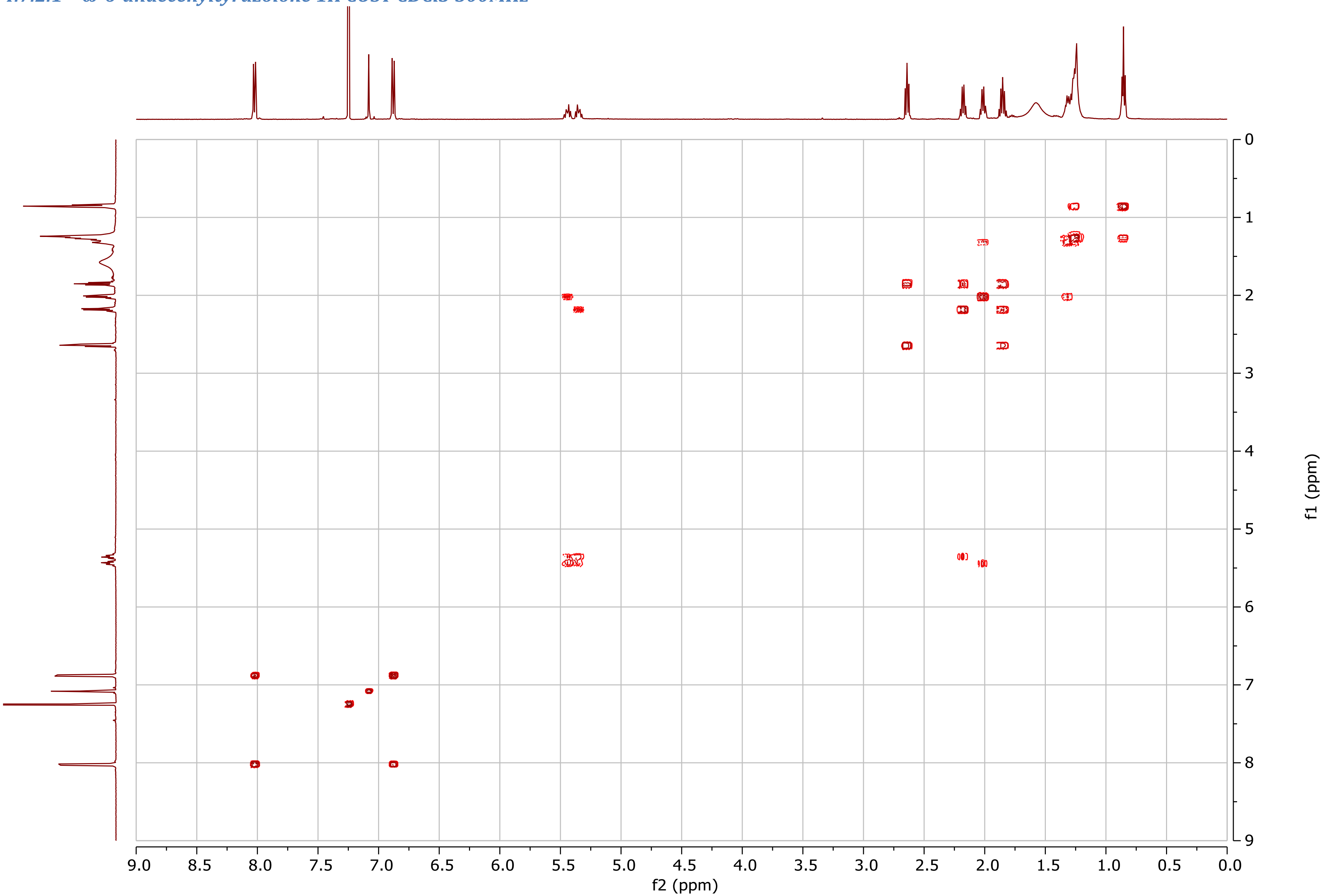

##### 4.7.3 N-decanoyldehydrotyrosine methyl ester

###### *4.7.3.1 N-decanoyldehydrotyrosine methyl ester 1H COSY MeOH-d4 500MHz*

Note: contaminated with ethyl acetate

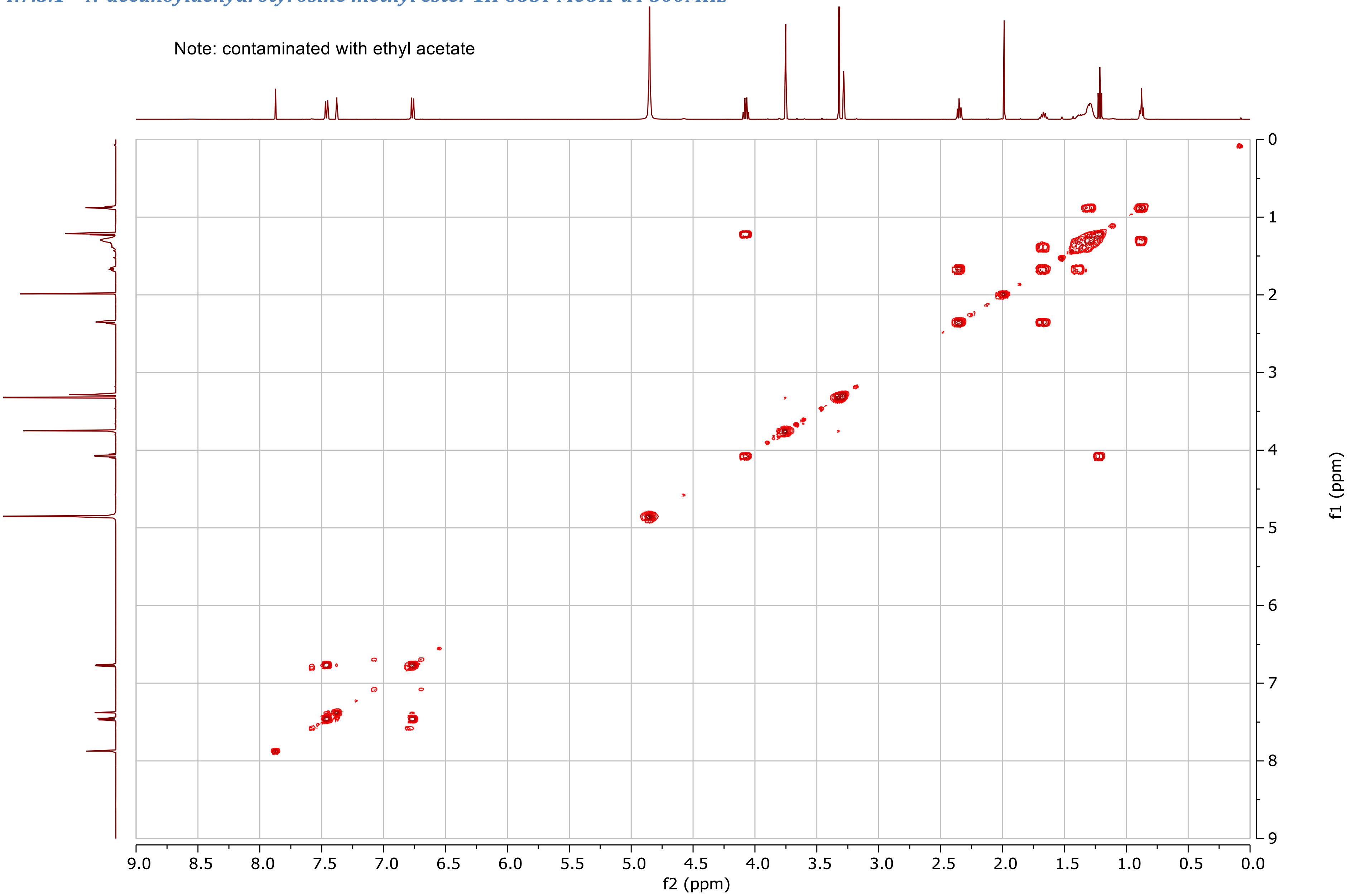

##### 4.7.4 N-octanoyldehydrophenylalanine methyl ester

###### *4.7.4.1 N-octanoyldehydrophenylalanine methyl ester 1H COSY DMSO-d6 500MHzv*

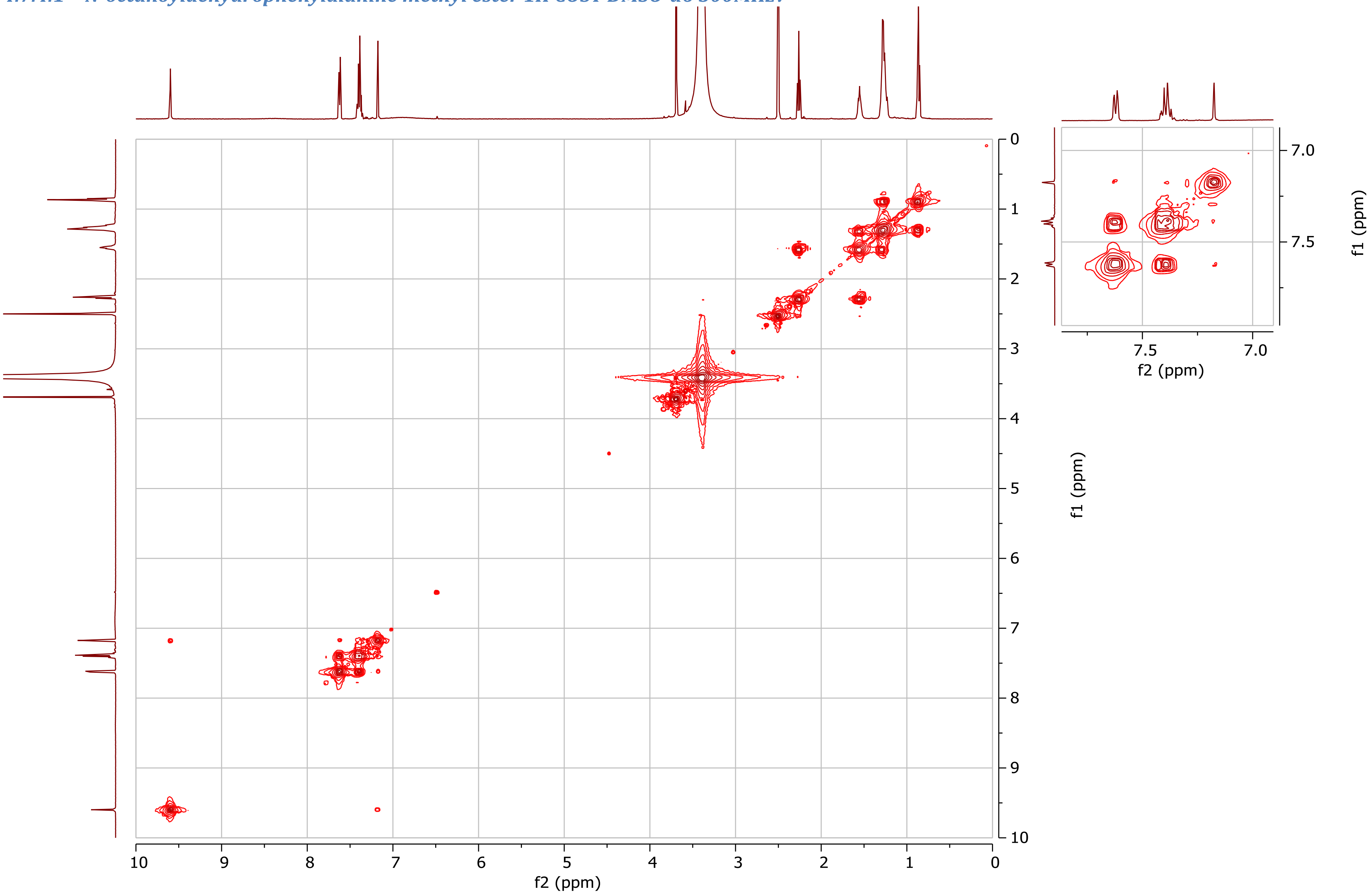

#### 4.8 $^{1}H$ NOESY NMR

##### 4.8.1 Undecyltyrazolone

###### *4.8.1.1 Undecyltyrazolone 1H NOESY CDCl3 500MHz*

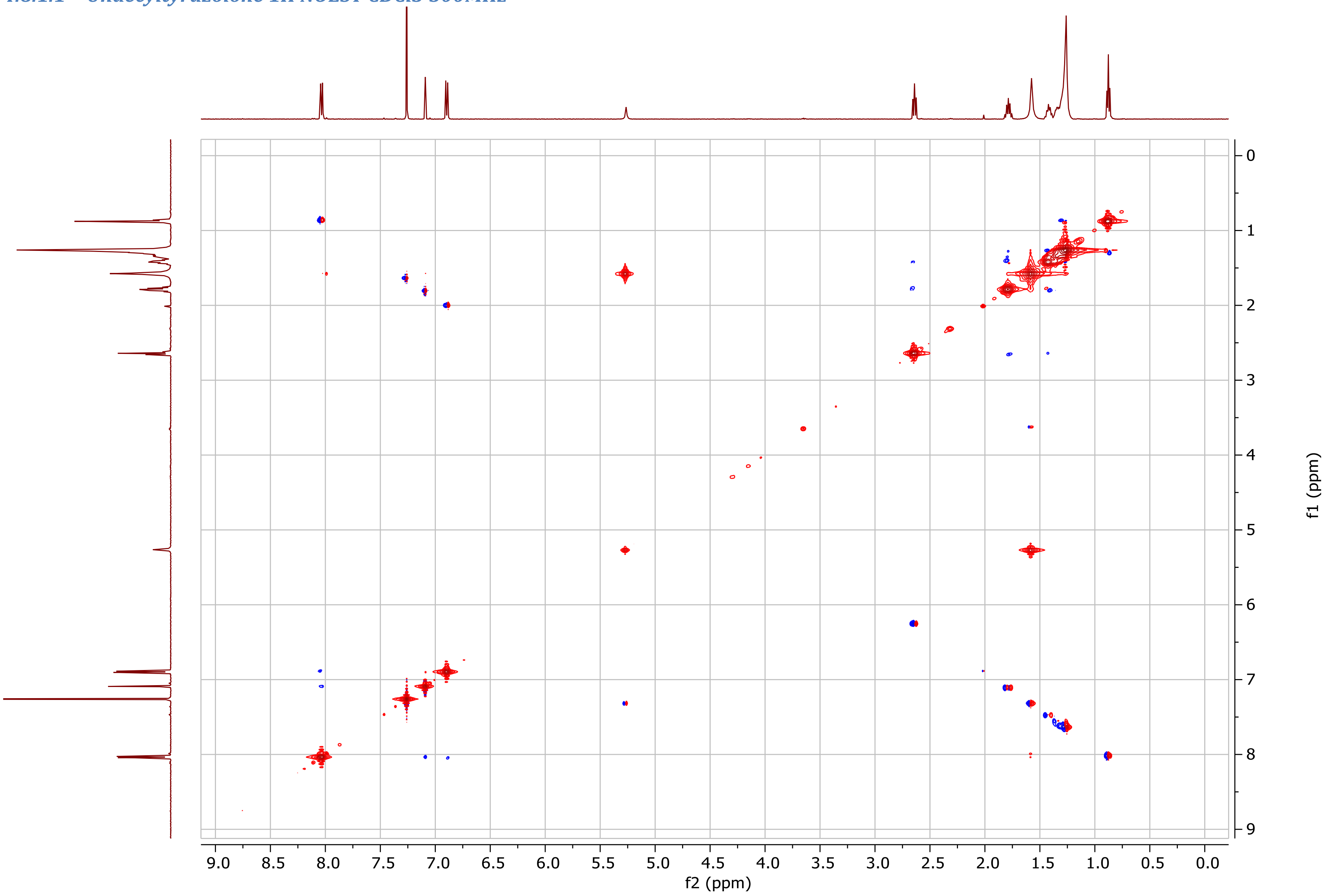

##### 4.8.2 N-decanoyldehydrotyrosine methyl ester

###### *4.8.2.1 N-decanoyldehydrotyrosine methyl ester 1H NOESY MeOH-d4 500MHz*

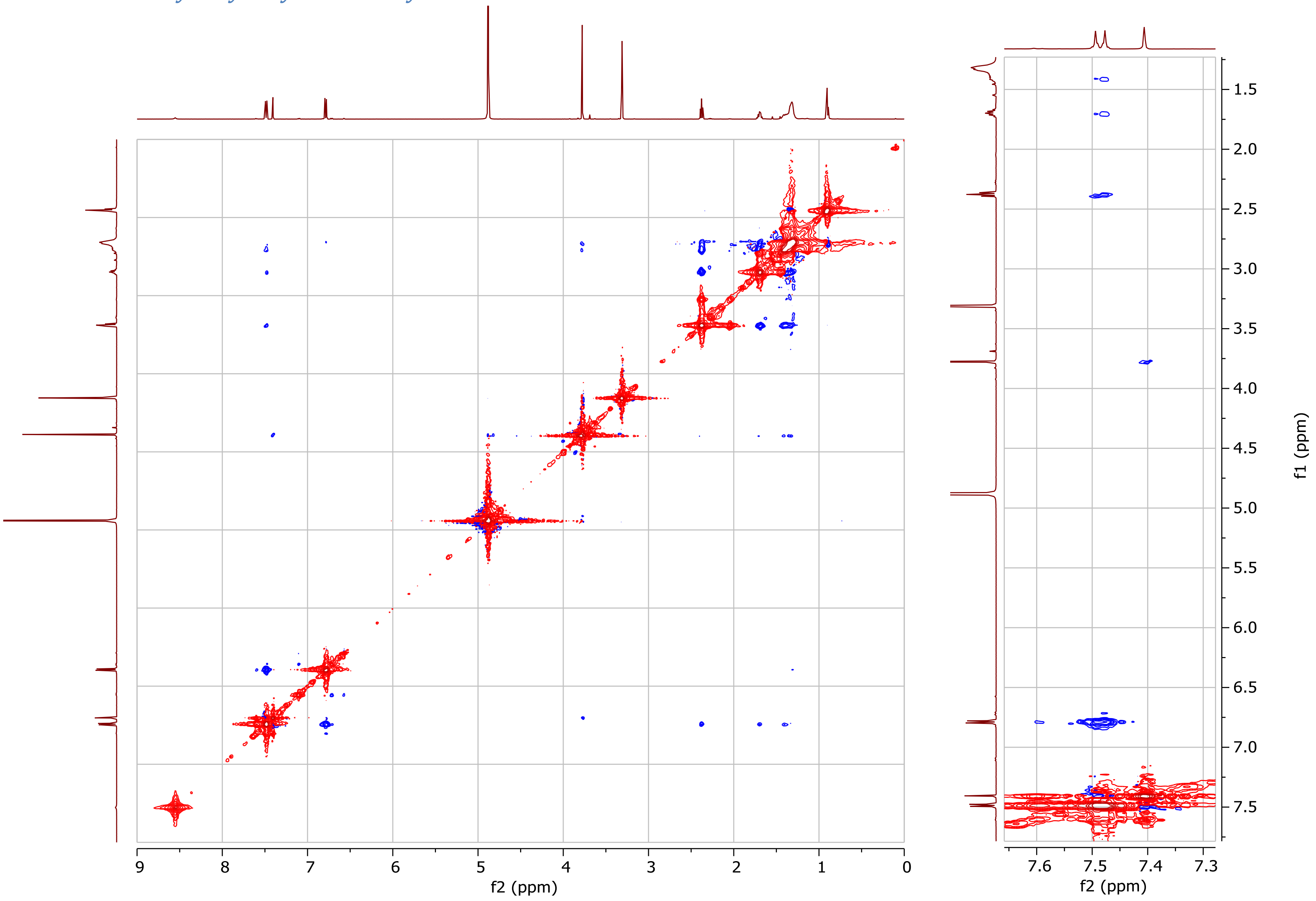

##### 4.8.3 N-octanoyldehydrophenylalanine methyl ester

###### *4.8.3.1 N-octanoyldehydrophenylalanine methyl ester 1H NOESY DMSO-d6 500MHz*

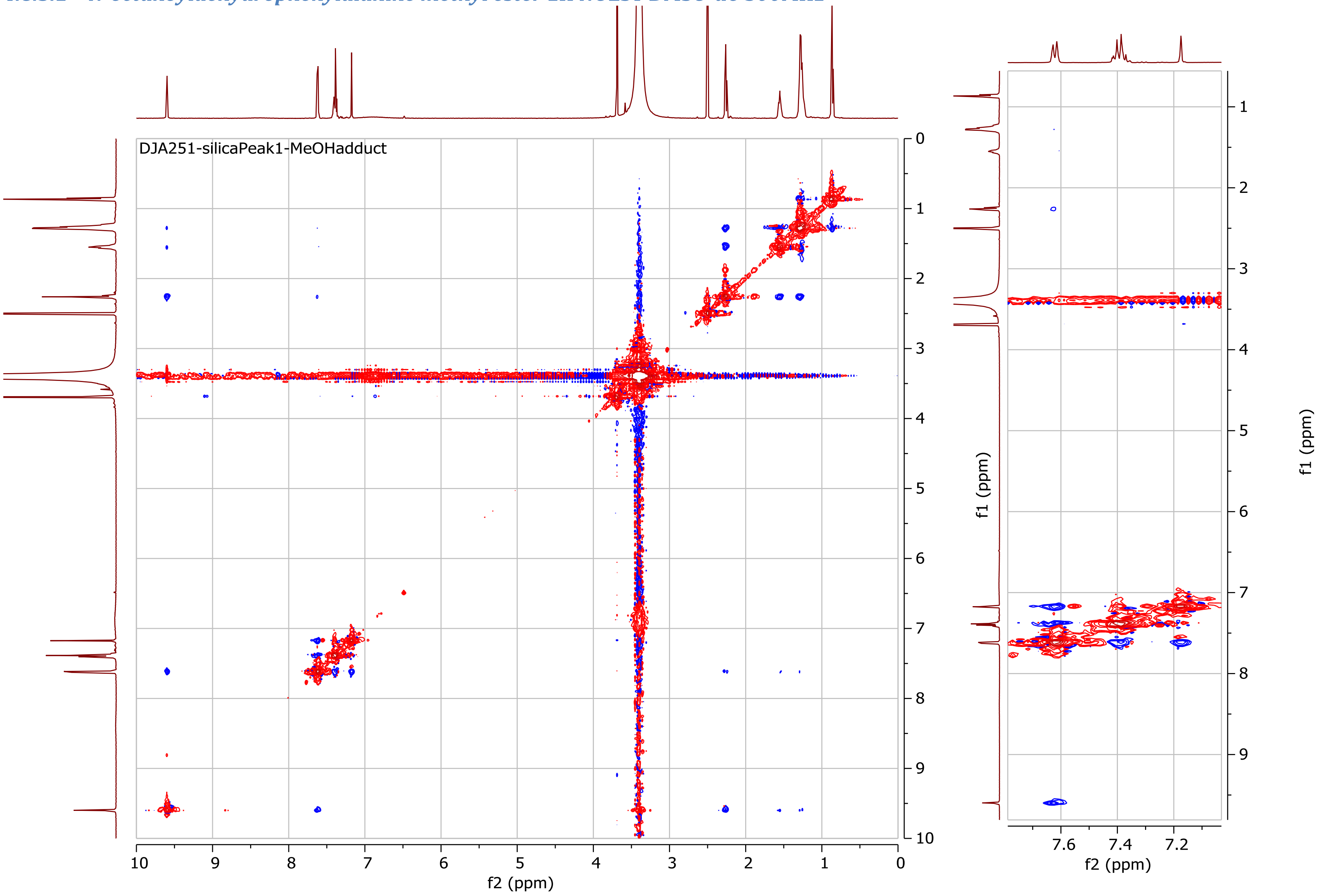

### 5 Other spectral data

#### 5.1 Infrared (IR) spectrum for nonyltyrazolone

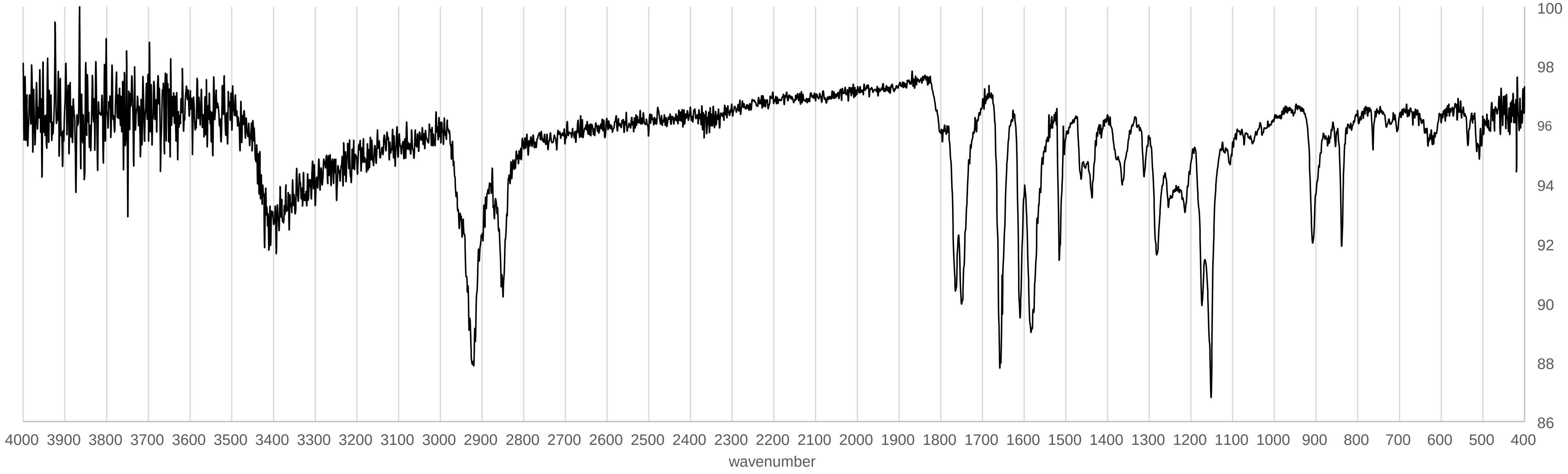

#### 5.2 Tandem mass spectra

Collision-induced dissociation tandem mass spectra have been deposited in the GNPS[9] library with the following spectrum IDs:

Hexyltyrazolone: CCMSLIB00005723541 (https://gnps.ucsd.edu/ProteoSAFe/gnpslibraryspectrum.jsp?SpectrumID=CCMSLIB00005723541)
Heptyltyrazolone: CCMSLIB00005723543 (https://gnps.ucsd.edu/ProteoSAFe/gnpslibraryspectrum.jsp?SpectrumID=CCMSLIB00005723543)
Octyltyrazolone: CCMSLIB00005723544 (https://gnps.ucsd.edu/ProteoSAFe/gnpslibraryspectrum.jsp?SpectrumID=CCMSLIB00005723544)
Nonyltyrazolone: CCMSLIB00005716822 (https://gnps.ucsd.edu/ProteoSAFe/gnpslibraryspectrum.jsp?SpectrumID=CCMSLIB00005716822)
Decyltyrazolone: CCMSLIB00005723362 (https://gnps.ucsd.edu/ProteoSAFe/gnpslibraryspectrum.jsp?SpectrumID=CCMSLIB00005723362)
Undecyltyrazolone: CCMSLIB00005723361 (https://gnps.ucsd.edu/ProteoSAFe/gnpslibraryspectrum.jsp?SpectrumID=CCMSLIB00005723361)
ω-6-undecenyltyrazolone: CCMSLIB00005723363 (https://gnps.ucsd.edu/ProteoSAFe/gnpslibraryspectrum.jsp?SpectrumID=CCMSLIB00005723363)
Hexylphenazolone: CCMSLIB00005723542 (https://gnps.ucsd.edu/ProteoSAFe/gnpslibraryspectrum.jsp?SpectrumID=CCMSLIB00005723542)
Heptylphenazolone: CCMSLIB00005723540 (https://gnps.ucsd.edu/ProteoSAFe/gnpslibraryspectrum.jsp?SpectrumID=CCMSLIB00005723540)
Octylphenazolone: CCMSLIB00005723545 (https://gnps.ucsd.edu/ProteoSAFe/gnpslibraryspectrum.jsp?SpectrumID=CCMSLIB00005723545)
Nonylphenazolone: CCMSLIB00005723546 (https://gnps.ucsd.edu/ProteoSAFe/gnpslibraryspectrum.jsp?SpectrumID=CCMSLIB00005723546)

Notes:

- These spectra tend to network with each other when performing spectral networking, except for ω-6-undecenyltyrazolone.
- (*E*)-and (*Z*)-tyrazolones produce identical tandem MS spectra. We suspect they isomerize during the CID process.
- Hydrolyzed oxazolones (i.e., acyldehydrotyrosines and acyldehydrophenylalanines, which appear sometimes if samples are sitting in acetonitrile:water for a long time before being analyzed by HPLC-UV-MS) show tandem MS very similar to their respective oxazolones. We suspect the former undergo spontaneous cyclodehydration to form their respective oxazolones during the CID process.

### 6 Bibliography

(1) Kautsar, S. A.; Blin, K.; Shaw, S.; Navarro-Muñoz, J. C.; Terlouw, B. R.; van der Hooft, J. J. J.; van Santen, J. A.; Tracanna, V.; Suarez Duran, H. G.; Pascal Andreu, V.; et al. MIBiG 2.0: a Repository for Biosynthetic Gene Clusters of Known Function. *Nucleic Acids Res* **2020**, *48*, D454–D458.

(2) Bentley, S. D.; Chater, K. F.; Cerdeño-Tárraga, A. M.; Challis, G. L.; Thomson, N. R.; James, K. D.; Harris, D. E.; Quail, M. A.; Kieser, H.; Harper, D.; et al. Complete Genome Sequence of the Model Actinomycete Streptomyces Coelicolor A3(2). *Nature* **2002**, *417*, 141–147.

(3) Udwary, D. W.; Zeigler, L.; Asolkar, R. N.; Singan, V.; Lapidus, A.; Fenical, W.; Jensen, P. R.; Moore, B. S. Genome Sequencing Reveals Complex Secondary Metabolome in the Marine Actinomycete Salinispora Tropica. *Proc Natl Acad Sci U S A* **2007**, *104*, 10376–10381.

(4) Paulsen, I. T.; Press, C. M.; Ravel, J.; Kobayashi, D. Y.; Myers, G. S. A.; Mavrodi, D. V.; DeBoy, R. T.; Seshadri, R.; Ren, Q.; Madupu, R.; et al. Complete Genome Sequence of the Plant Commensal Pseudomonas Fluorescens Pf-5. *Nat Biotechnol* **2005**, *23*, 873–878.

(5) UniProt Consortium, T. UniProt: The Universal Protein Knowledgebase. *Nucleic Acids Res* **2018**, *46*, 2699.

(6) Shannon, P.; Markiel, A.; Ozier, O.; Baliga, N. S.; Wang, J. T.; Ramage, D.; Amin, N.; Schwikowski, B.; Ideker, T. Cytoscape: a Software Environment for Integrated Models of Biomolecular Interaction Networks. *Genome Res* **2003**, *13*, 2498–2504.

(7) Blattner, F. R.; Plunkett, G.; Bloch, C. A.; Perna, N. T.; Burland, V.; Riley, M.; Collado-Vides, J.; Glasner, J. D.; Rode, C. K.; Mayhew, G. F.; et al. The Complete Genome Sequence of Escherichia Coli K-12. *Science* **1997**, *277*, 1453–1462.

(8) Xie, B.-B.; Shu, Y.-L.; Qin, Q.-L.; Rong, J.-C.; Zhang, X.-Y.; Chen, X.-L.; Zhou, B.-C.; Zhang, Y.-Z. Genome Sequence of the Cycloprodigiosin-producing Bacterial Strain Pseudoalteromonas Rubra ATCC 29570(T). *J Bacteriol* **2012**, *194*, 1637–1638.

(9) Wang, M.; Carver, J. J.; Phelan, V. V.; Sanchez, L. M.; Garg, N.; Peng, Y.; Nguyen, D. D.; Watrous, J.; Kapono, C. A.; Luzzatto-Knaan, T.; et al. Sharing and Community Curation of Mass Spectrometry Data with Global Natural Products Social Molecular Networking. *Nat Biotechnol* **2016**, *34*, 828–837.
